# Comprehensive molecular impact mapping of common and rare variants at GWAS loci

**DOI:** 10.1101/2025.06.05.658079

**Authors:** Brad Balderson, Sanjana Tule, Mei-Lin Okino, William JF Rieger, Sierra Corban, Jeff Jaureguy, Nathan Palpant, Kyle J. Gaulton, Loïc Yengo, Mikael Bodén, Graham McVicker

## Abstract

Deep learning sequence to function models can predict the molecular effects of genetic variants, but their predictions are limited to the cell types and assays they are trained on. Here we describe ***DNACipher***, a deep learning model that predicts the effects of genetic variants across diverse biological contexts—including those not directly measured.

DNACipher takes 524 kb of genome sequence as input and predicts variant effects across 38,582 cell type–assay combinations, > 6 times as many contexts as AlphaGenome. We also introduce DNACipher **Deep Variant Impact Mapping**, a method to prioritize variants within trait-associated loci based on elevated predicted molecular effects relative to non-trait-associated variants. Applied to 590,604 common and rare variants at GWAS loci across 10 complex traits, DVIM identified imVars at > 90% of GWAS loci for 9 out of 10 traits. Common imVars were significantly concordant with statistical fine-mapping, and rare imVars were enriched for prior trait associations. For type 1 diabetes (T1D), imVars also had predicted effects that were supported by single-nucleus ATAC-seq, luciferase assays, and massively parallel reporter assays. DNACipher DVIM prioritises common and rare variants at GWAS loci by predicting molecular effects across cellular contexts.

## Introduction

GWAS have identified thousands of variants associated with complex traits and diseases, however it remains difficult to distinguish causal variants of phenotype associations from non-causal variants in linkage disequilibrium (LD)^1^. Statistical fine-mapping can reduce the number of candidate causal variants substantially, but often leaves substantial ambiguity, especially within high LD regions. Additionally, GWAS are underpowered to detect associations with rare genetic variants, due to the small number of allele carriers^1^. Moreover, how the causal variants affect genome function in specific cellular contexts is largely unknown^2^. Methods that intersect variants with functional genomic data or evolutionary sequence conservation can prioritize likely causal variants and provide biological context for their potential molecular effects^3–9^. However, these approaches rely on overlap with genome annotations that are hundreds to thousands of bases long, and therefore cannot disentangle causal variants from non-causal variants that share annotations and are in high LD^10,11^.

Sequence to function (S2F) deep learning models predict genomic measurements from DNA sequence, enable single base-pair predictions, and have the potential to distinguish causal from non-causal GWAS variants at risk loci^12–19^. By comparing predicted signals from reference and alternative sequences, these models can, in principle, also infer the effects of a genetic variant for specific cell types and assays, and can do so for both common and rare genetic variants^20^.

Since most GWAS variants are non-coding^21^ and may affect the expression of genes that are over 1 million bp away^22,23^, recent improvements to S2F models have concentrated on expanding the maximum sequence length they take as input^15,17,19^. The largest models, AlphaGenome^24^, Borzoi^19^ and Enformer^17^, have sequence inputs (’receptive fields’) of 1 Mb, 524 kb and 196 kb, which can be used to predict the long range regulatory effects of noncoding variants.

A major limitation of current S2F models is that they can only generate predictions for the experiments (e.g. RNA-seq in heart tissue) that they have been trained on. While models such as AlphaGenome have been trained on a large number of experiments (5,930 human and 1,128 mouse)^24^, this represents only a small fraction of the experiments that could be performed on different cell types and tissues^25,26^. In addition, the available training data used by these models are imbalanced and have high ascertainment bias. For example, of the 12,869 human genomics experiments on ENCODE, 11% are DNase-seq experiments and 16% are from just two cell lines (K562 and HepG2)^27^.

The sparsity of experimental genomics data has motivated methods to impute missing experiments, such as ChromImpute^26,28^ and Avocado^25^. Avocado is a deep learning model that utilizes multiple one-hot-encoded inputs—including genome location, cell type, and assay—to predict experimental signals for these inputted biological contexts. By encoding cell type and assay as model input, Avocado can predict experimental signals for experiments that have not been performed^25^. However, Avocado does not utilise DNA sequences during prediction, and therefore cannot predict how genetic variants can affect the unobserved experiments that it imputes.

Recently, several S2F models have been developed to predict unobserved experimental signals including EpiGePT^29^, Enformer Celltyping^30^, and DeepCT^31^. Both EpiGePT and Enformer Celltyping take long sequence inputs (128 kb and 196 kb), but only predict signals for a small number of assays (8 and 6 assays) and require cell type-specific gene expression or chromatin accessibility as inputs. On the other hand, DeepCT predicts up to 31,760 cell type-assay combinations. However it does not predict gene expression or long-range variant effects because its input sequences are limited to 1 kb. Thus, current methods for predicting experimental signals for unobserved experiments from DNA sequences are limited because they either cannot take long sequences as input or cannot impute signals for many unobserved experiments.

Here, we develop a new model, DNACipher, which leverages DNA-sequence embeddings from any sequence model, including long-range models such as Borzoi^19^ or Enformer^17^, and combines these sequence embeddings with the imputation capability of Avocado^25^. Our approach allows long-range effect variant predictions across a broad range of observed and unobserved biological contexts (Figure 1A-C). After training on ENCODE data, using Borzoi sequence embeddings, “DNACipher-Borzoi” can predict functional variant effects in 38,582 distinct contexts (191 biological samples across 202 assays) from input sequences of 524 kb (Figure 1D-E). We validate DNACipher on held-out experiments and fine-mapped GTEx expression quantitative trait loci (eQTLs), and demonstrate that it can detect > 14-fold more molecular effects of high-confidence causal-eQTLs compared to AlphaGenome.

**Figure 1.**
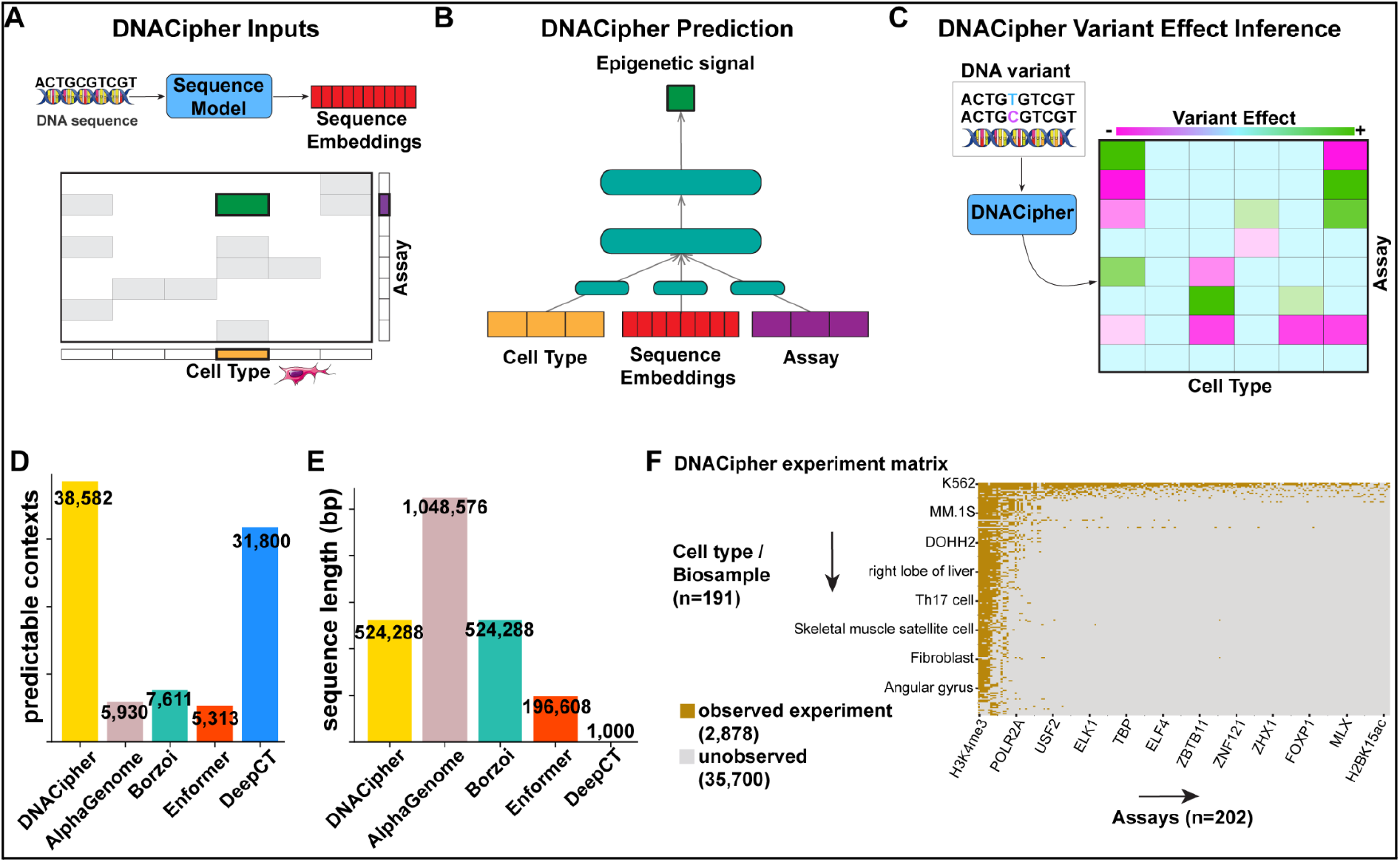
DNACipher inference of genetic variant effects for unobserved experiments. **A.** Schematic of DNACipher input, which includes sequence embeddings from any DNA sequence model, and experiments representing combinations of cell types (e.g. heart, fibroblast) and assays (e.g. RNA-seq, ATAC-seq, ChIP-seq). Cell type/assay combinations which have been experimentally observed are highlighted, while cell type/assay combinations which are imputable are blank. **B.** Schematic of the DNACipher model, which takes 3 inputs: one-hot encoded cell type, one-hot-encoded assay, and a DNA sequence embedding from a DNA sequence model. Linear layers parse each of the one-hot-encoded inputs independently, before they are concatenated and parsed through deeper neural network layers to predict the signal of the inputted sequence for the given cell type/assay combination. **C.** DNACipher infers variant effects for unobserved cell type/assay combinations by contrasting predicted signal predictions between DNA sequence variants for all cell type and assay combinations. **D.** The number of predicted cell type/assay contexts for DNACipher compared to other sequence models. **E.** The DNA sequence length which can be modelled using DNACipher with Borzoi sequence embeddings, compared to other sequence models. **F.** The cell type/assay matrix of observed and unobserved experiments from ENCODE, showing the high ascertainment bias for assays performed on particular cell types such as K562 cells. Most potential experiments have not been performed, and so imputation vastly increases variant effect predictions.

We also develop Deep Variant Impact Mapping (DVIM), a method to prioritize trait-associated variants based on their DNACipher-predicted effects in relevant cell types (‘impact’ variants, imVars). We apply DVIM to GWAS loci across 10 complex traits, and find that imVars have significantly higher causal probabilities from statistical fine-mapping.

Furthermore, imVars were detected at > 90% of GWAS loci for 9 of 10 complex traits. This indicates a significant advance for identifying potential trait-mediating molecular effects compared with eQTLs which only colocalise with a median of 21% of GWAS loci across traits^32^. DVIM can also be applied to both common and rare genetic variants, and we use it to identify 30,561 rare imVars across the 10 traits, with significant enrichment for prior trait associations.

## Results

The DNACipher model takes three inputs: embeddings for the genomic sequence, one-hot encoded cell type, and one-hot encoded assay. Latent representations of cell type and assay are learned by the model, whereas the sequence embeddings are provided from another pre-trained model. These inputs are then passed into a multi-layer perceptron with ∼1.3 million parameters (Figure 1A-B). This design makes it possible to predict variant effects for all combinations of cell types and assays, including those which have not been observed (Figure 1C). In addition, because DNACipher takes sequence embeddings from pre-trained models, different sequence models can be used, including those with long sequence contexts, without requiring high compute resources. Hence, DNACipher enables long-range variant effect predictions in unobserved biological contexts and it can be trained on a single A40 GPU (Figure 1D-E).

### DNACipher predicts genomic signals from DNA sequence for unobserved cell types and assays

After establishing prediction to unobserved experimental signals with short sequence models (Supp. Note 1), we next trained models that utilized either Enformer embeddings (“DNACipher-Enformer”) or Borzoi embeddings (“DNACipher-Borzoi”) (Figure 2A). Potential pre-training leakage was controlled using 5-fold cross-validation, where each test-fold represented re-allocation of test cell type/assays not included in the Borzoi/Enformer training dataset. The cell types chosen for testing for a given assay were those with the fewest number of other assays measured, to ensure our performance metrics best reflect the majority of cell types that have sparse experimental measurements (Figure 1F, Figure S2).

**Figure 2.**
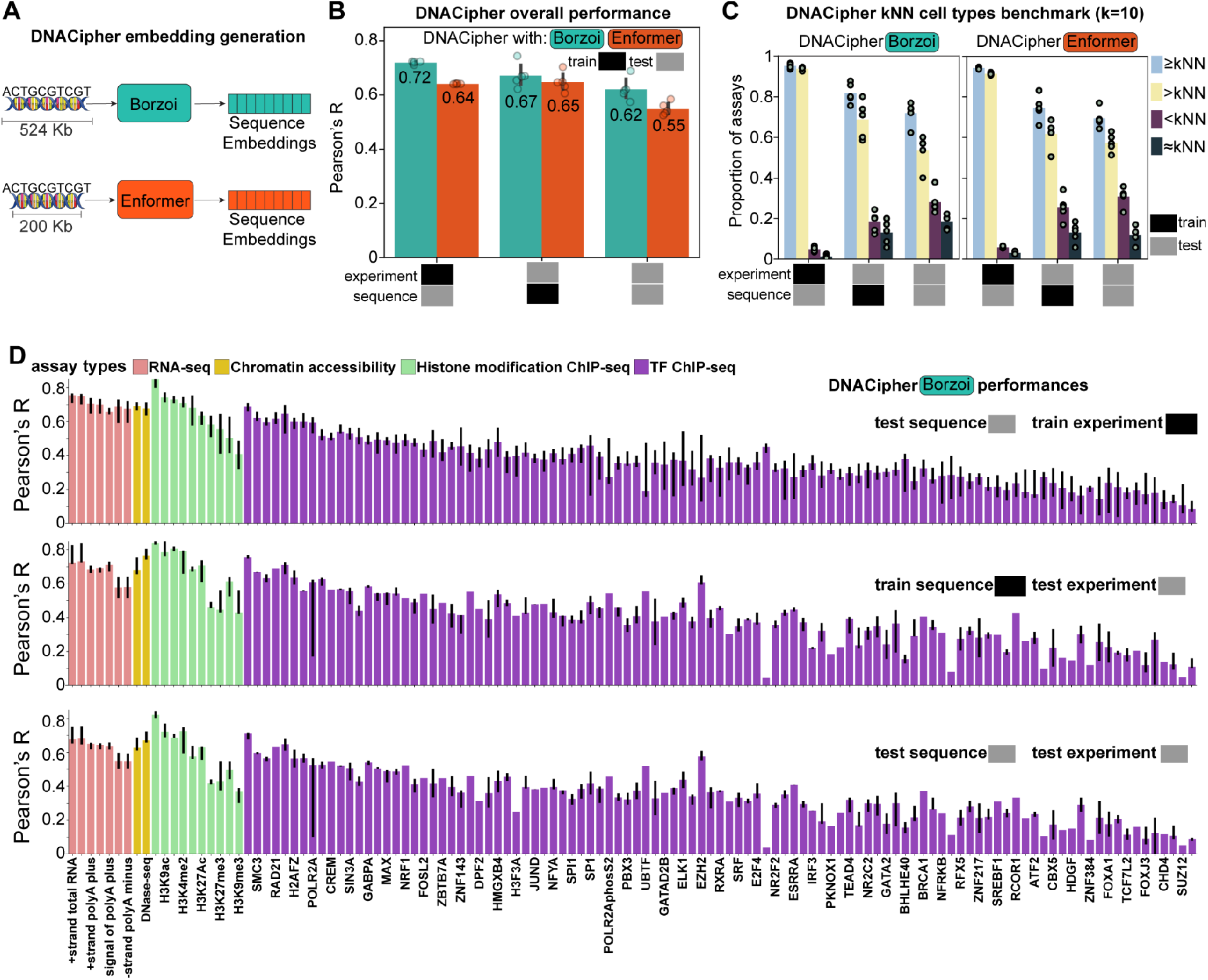
DNACipher predicts unobserved experiment signals with long sequences. **A.** Schematic depiction of long sequence embedding generation from Borzoi or Enformer, which were computed using the DNACipher training framework. **B.** Barcharts with mean Pearson’s correlation (R) for DNACipher trained with sequence embeddings from either Borzoi (‘DNACipher-Borzoi’) or Enformer (‘DNACipher-Enformer’) between the predicted and observed genomic signals. Correlations are stratified by test sequences (N = 345,371) in train experiments (observed, N = 2,759-2,847 across test folds), train sequences (N = 1,162,549) in test experiments (unobserved, N = 123-35 across test folds), and test sequences in test experiments. Points are correlations from 5 cross-validations of train/test experiments, and error bars are 95% confidence intervals. **C.** Barcharts for DNACipher-Borzoi and DNACipher-Enformer, which depict the mean proportion of assays where DNACipher imputation predicted assay signals equivalently or significantly better than a mean kNN cell type baseline (≥kNN; k = 10; two-sided paired Wilcoxon rank-sum test on Pearson’s R across N = 1,350 batches of 256 sequences, p < 0.05), significantly better than the kNN baseline (>kNN), significantly worse than the kNN baseline (<kNN), or non-significantly different to the baseline (≈kNN). Performances are stratified by train/test sequences and train/test experiments, and points correspond to the result for each of 5 cross-validations. **D.** Barcharts of median Pearson’s R for DNACipher-Borzoi predictions stratified by assay, train/test sequences and train/test experiments. Correlations are compiled across the 5 cross-validations and all cell/tissue types in the relevant assay, with error bars indicating 0.25-0.75 quantiles. Error bars are missing where only one experiment could be utilized for the relevant stratification. Sample sizes are listed in Table S1.

Allocation of train/test sequences was concordant with Borzoi/Enformer pre-training. With this rigorous evaluation, we found high and consistent Pearson correlations across the 5-fold cross validations between the observed and predicted signals (Figure 2B). For test experiments and test sequences, DNACipher-Borzoi achieved a mean R of 0.62, and DNACipher-Enformer achieved a mean R of 0.55 across folds (Figure 2B). Performance was also high across the vast array of assay types (Figure 2D, Ext. Data Fig 1, Table S1), though we note a slightly reduced overall performance for test sequence and training experiments between DNACipher-Borzoi and the original Borzoi model (Figure S3). Overall, this confirmed DNACipher based on long-sequence model embeddings can impute signals for missing experiments.

To further assess the validity of the long-sequence DNACipher models, we compared signal prediction performance against the mean of k-nearest-neighbour (kNN) training cell types (Figure 2C, Figure S4). We used a semantic definition of nearest neighbours, based on cosine similarity within openAI embeddings of the cell/tissue type names (Table S1). Both DNACipher-Borzoi and DNACipher-Enformer out-performed the mean kNN cell type baseline for most assays, with DNACipher-Borzoi performing as well or better than the mean kNN baseline for 72.58% of the assays for test cell types and test sequences (Figure 2C, Figure S4, Table S1). This shows that DNACipher outperforms using assay measurements of closely related cell types as a proxy. We also found DNACipher-Borzoi signal predictions are significantly correlated for cell type-specific signals in test sequences (Figure S5), DNACipher models with short sequences had comparable signal imputation performance to long sequences (Supp. Note 2, Figure S6, Table S1), and DNACipher latent representations of cell types and assays reflect known biological relationships (Supp. Note 3, Figure S7,S8, Table S2). Overall these results indicate that DNACipher integrates information from sequence, cell type, and assay embeddings to accurately predict unobserved molecular measurements.

### DNACipher long-range variant effect prediction detects a causal eQTL effect missed by Enformer and Borzoi

To evaluate whether DNACipher can predict long-range effects of genetic variants^17^, we first focused on the *WRN* locus as an example to highlight the benefit of imputation (Figure 3), before performing a systematic comparison across thousands of eQTLs in GTEx (Figure 4). We chose this locus specifically because it was held-out during training (allocated to test sequences), and because it intersects a high posterior inclusion probability (PIP = 0.9999) prostate eQTL from GTEx^23^.

**Figure 3.**
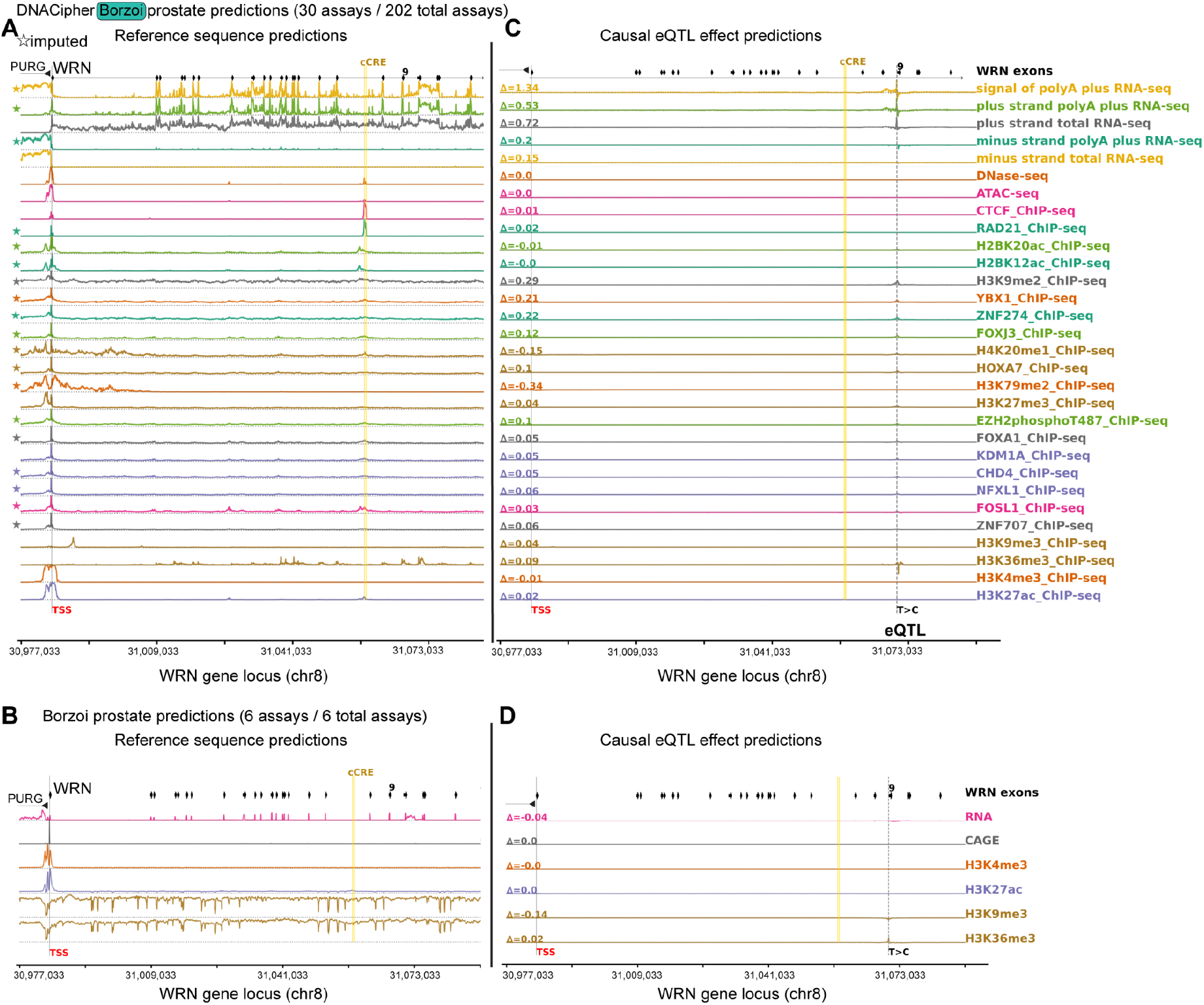
DNACipher-Borzoi predicts an eQTL effect that is not detected with Borzoi. **A.** DNACipher-Borzoi signal predictions for the top 30 prostate-specific assays (of 202 total) at the *WRN* locus reference genome sequence (test sequence). The x-axis is the genome position. Assays marked with stars are imputed. The *WRN* transcription start site (TSS), *WRN* exons, an ENCODE candidate Cis-Regulatory Element (cCRE), and opposite strand gene *PURG* are annotated. **B.** Equivalent to A, except with Borzoi prostate predictions, which are limited to 6 assays. **C.** Predicted change in assay signals between the reference and alternate alleles of a likely causal expression quantitative trait locus (eQTL, rs4733225) variant for *WRN*. Δ values are the predicted fold-change in signal (alternative-reference sequence predictions) across the WRN locus for the likely causal eQTL. **D.** Equivalent to C, except with Borzoi.

**Figure 4.**
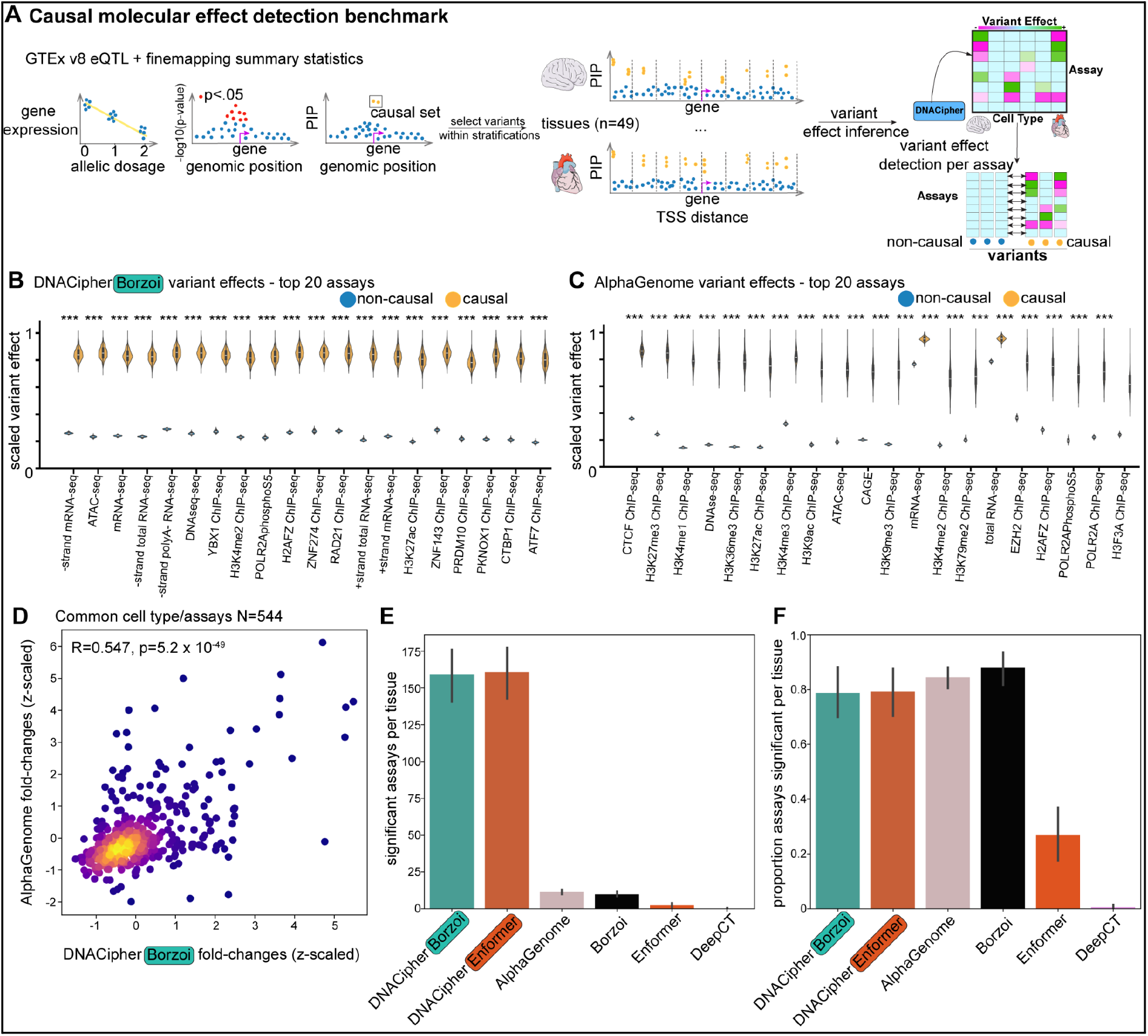
DNACipher improves detection of causal eQTL molecular effects. **A.** Schematic of balanced causal variant selection for benchmarking. Briefly, eQTLs and fine-mapping statistics were collected from GTEx v8. Likely causal (PIP > 0.9, N = 14,853) and non-causal (PIP < 0.1, N = 4,276) eQTL variants within tissues (N = 49) and 100 kb TSS-distance bins up to 1 Mb were selected for tissue-specific variant effect inference across all predictable assays. Causal and non-causal predicted assay effects for each tissue were then compared with a two-sided independent t-test with Benjamini-Hochberg False Discovery Rate (FDR) p-value correction. **B.** Violin plots with boot-strapped (N = 1,000) predicted variant effect means from DNACipher-Borzoi for likely causal and non-causal eQTL variants on the y-axis. Inner box-plots display the median value (white horizontal line), box edges are 25th and 75th percentiles, and whiskers are 1.5X the interquartile range. The top 20 most significant molecular assays are shown on the x-axis. **C.** Equivalent to B, except utilising AlphaGenome predictions. **D.** Scatter plot of z-scaled fold-changes between causal and non-causal eQTLs for each tissue and assay effect predicted by DNACipher-Borzoi (x-axis) and AlphaGenome (y-axis) (N = 544). **E.** Bar plots with the mean (across 10 random 90% re-samplings without replacement of variants) number of assays with predicted effect sizes that were significantly different between likely causal and non-causal eQTL variants for each tissue (N tissues = 49), using predictions from DNACipher-Borzoi (N assays = 202), DNACipher-Enformer (N assays = 202), AlphaGenome (N assays = 43), Borzoi (N assays = 39), Enformer (N assays = 37), and DeepCT (N assays = 40). Error bars are 95% confidence intervals. **F.** Equivalent to E, except the proportion of significant assays which were predictable by a given model for a given tissue is shown on the y-axis (N = 49).

We predicted signals across the DNA sequence surrounding this causal eQTL for 202 prostate assays using both DNACipher-Borzoi (Figure 3A) and DNACipher-Enformer (Ext. Data Fig. 2A). Because prostate tissue has sparse experimental data within ENCODE, Borzoi could only generate predictions for 6 prostate assays (Figure 3B). Enformer was similarly limited, but also lacked RNA-seq (Ext. Data Fig. 2B). Despite the underlying Enformer model not being trained on RNA-seq, DNACipher-Enformer predicted positive-strand RNA-seq expression that was specific to *WRN*’s exons and corresponded to the orientation of the gene (Ext. Data Fig. 2A), consistent with the ground-truth gene expression data (Ext. Data Fig. 3). DNACipher-Borzoi also exhibited accurate predictions on RNA-seq signals (Figure 3A), but had better strand-specificity. At the *WRN* transcription start site (TSS), both models predicted chromatin accessibility (DNAse-seq and ATAC-seq) and transcription-associated histone modifications (H3K4me3 and H3K27ac), and at an ENCODE candidate cis-regulatory element (cCRE), the models predicted CTCF and RAD21 ChIP-seq signals as well as nearby enhancer-associated histone modifications (H2BK20ac and H2BK12ac)^33^ (Figure 3A, Ext. Data Fig. 3A). Together, these observations illustrate that the long-sequence DNACipher models reconstruct complex regulatory landscapes even in unobserved experiments across an unobserved DNA sequence.

To test DNACipher’s ability to predict the long-range effects of genetic variants, we utilized the likely causal eQTL variant for *WRN* (rs4733225, PIP=0.9999, T>C substitution) and calculated a fold-change of predicted signals from the reference and alternative sequences (Δ; Figure 3C, Ext. Data Fig. 3C). Both models correctly predicted an overall increase in gene expression for the alternative allele across the *WRN* locus, which was specific to the RNA-seq assays (Δ0.17-Δ0.72 plus strand total RNA-seq; Figure 3C, Ext. Data Fig. 3C). This effect direction was consistent with the GTEx eQTL effect (β=0.34). In comparison, both Borzoi and Enformer’s prostate-specific predictions did not capture the eQTL effect on any assays (Figure 3D, Ext. Data Fig. 3D). This result is expected for Enformer which lacks RNA-seq training, but is surprising for Borzoi, and could be due to lack of strand-specificity for the prostate RNA-seq assay in this particular case. Overall, these findings illustrate that the long-range DNACipher models can predict variant effects over large genome regions.

### DNACipher predicts molecular QTL effect sizes with similar performance to AlphaGenome

We next evaluated the DNACipher models molecular QTL effect size prediction, in comparison with AlphaGenome and Borzoi. We benchmarked effect size prediction for eQTLs (Ext. Data Fig. 4), TF allele-specific binding (ASB) (Ext. Data Fig. 5), and DNase I hypersensitivity QTLs (dsQTLs) (Ext. Data Fig. 6). For eQTLs, we predicted effects of likely causal eQTLs (PIP > 0.99) on their associated eGene across all GTEx tissues (N = 14,196 eQTL-eGene-tissue effects). Overall, the average correlation across models between estimated and predicted effect sizes was R = 0.22 and was not significantly different across models (Ext. Data Fig. 4A,B). For non-zero predicted eQTL effects to account for potential false positive causal variants^34^, model correlations increased to a mean R = 0.54 but remained non-significantly different (Ext. Data Fig. 4A,C). The correlations for eQTLs at different distances to the eGene TSS were also similar (Ext. Data Fig. 4C,D), although for distal (> 12 kb) non-zero predicted eQTLs DNACipher-Borzoi (R = 0.70) and Borzoi (R = 0.66) had greater correlations than DNACipher-Enformer (R = 0.52) and AlphaGenome (R = 0.48).

For TF ASB prediction, we utilized the ADASTRA database^35^, which collates ASB effect sizes across 674 human TFs and 337 cell types. Correlations were also highly concordant amongst the models, and non-significantly different for overall TF binding effect sizes (Ext. Data Fig. 5). Because of missing data, most TF ASB events could not be predicted for the correct cell/tissue type using AlphaGenome or Borzoi, and in these cases using another tissue prediction as a proxy resulted in underperformance compared to the DNACipher models (Ext. Data Fig. 5). This demonstrates the importance of tissue-specific predictions for accurate effect size inference.

For dsQTL effect size benchmarking, we used dsQTLs from lymphoblastoid cell lines (LCLs, N = 574) used by previous benchmarks^36^. Models achieved a mean R = 0.695, and were non-significantly different (Ext. Data Fig. 6A,B). When we considered the unsigned (magnitude) of effect size prediction, DNACipher-Enformer (R = 0.28) significantly outperformed AlphaGenome (R = 0.095) and Borzoi (R = 0.1) (Ext. Data Fig. 6A,C). When mismatching the dsQTL tissue prediction, the DNACipher models outperformed AlphaGenome and Borzoi on both signed and unsigned dsQTL effect size prediction, again demonstrating the importance of our imputation approach to enable accurate molecular effect size prediction. Additionally, we performed saturated in silico mutagenesis^37^ to identify the motifs DNACipher-Enformer used for these dsQTL predictions (Figure S10). This identified high attribution at 219 motifs at the dsQTL location which could be matched to known TFs (Figure S10), confirming the model is utilizing biologically reasonable sequence features to make predictions.

Overall, the DNACipher models predict effect sizes across molecular phenotypes with similar performance to AlphaGenome and Borzoi, and exceed these models when the relevant cell type/tissue is missing.

### Inference to unobserved experimental contexts enhances detection of eQTL molecular effects

Based on our observations at the *WRN* locus, we hypothesized that the DNACipher models would outperform other S2F models at identifying context-specific molecular effects of genetic variants. We compared likely causal and likely non-causal eQTL variants from GTEx across 49 tissues. For each tissue, we binned eQTLs by their distance to the eGene’s TSS and selected 3 likely-causal eQTLs (PIP > 0.9) and 3 likely non-causal eQTLs (PIP < 0.1) within each bin (Figure 4A, Figure S9A). This yielded 23,058 eQTL variants with a balanced representation of tissue types, TSS distances, and expected regulatory impacts (Figure 4A, Figure S9B-D). We then matched each GTEx tissue to a corresponding DNACipher tissue and predicted variant effects across all 202 assays. For each assay, we compared the absolute predicted effect sizes of likely causal variants and non-causal variants (Student’s two-sided t-test) (Figure 4A-B, Table S3).

For comparison, we repeated this analysis with AlphaGenome, Borzoi, Enformer, and DeepCT (Figure 4). Consistent with our molecular effect size benchmarks, fold-changes between DNACipher-Borzoi and AlphaGenome between causal and non-causal eQTLs for common tissues and assays were highly concordant (R = 0.55, Figure 4D). Because DNACipher is able to impute molecular effects into unobserved experiments, the DNACipher models were able to detect many more tissue-specific molecular effects (mean N = 160), compared to AlphaGenome (mean N = 11.3), Borzoi (mean N = 9.9), Enformer (mean N = 2.23), and DeepCT (mean N = 0.15) (Figure 4E, Table S3). When comparing the proportion of the represented assays with significant effects for each tissue type, the DNACipher models were not significantly different to AlphaGenome and Borzoi (Figure 4F), but predicted a higher proportion of significant molecular effects compared to Enformer and DeepCT, suggesting greater sensitivity. This demonstrates that by imputing into unobserved biological contexts, DNACipher is able to detect a wider spectrum of molecular variant effects.

### DNACipher Deep Variant Impact Mapping (DVIM) identifies likely causal variants underlying Type 1 Diabetes (T1D) GWAS loci

We next developed an approach to utilize DNACipher for prioritizing likely causal variants at GWAS loci and determining their context-specific molecular effects, which we term Deep Variant Impact Mapping (DVIM). To develop our approach, we focused on genetic variants at GWAS hit loci for type 1 diabetes (T1D)^38^. Briefly, DVIM performs a statistical comparison between the predicted molecular effects of trait-associated (‘candidate’, p < 5 × 10^−7^) and confident non-trait associated (‘background’) variants at a given locus (common variants with minor allele frequency (MAF) > 5% and nominal p-value > 0.001), in order to call significant predicted effects (Figure 5A,B). DVIM uses null distributions from predicted background variant effects specific to each locus and molecular context (cell type/assay), since we observed large differences in this null distribution depending on the locus and molecular context (Figure S11). Furthermore, in-part because of a difference in the number of background variants at each locus, DVIM uses boot-strap resampling of background variants to re-calculate null distributions and p-values, since we observed locus and molecular context specific uncertainty in the estimated null distributions from boot-strap resampling (Figure S12,S13). Finally, because GWAS are under-powered to detect associations with rare variants and our approach is not dependent on MAF, we additionally used DVIM to test rare variants that occurred within the same region as the candidate common variants (MAF < 5%; Figure 5A). Candidate common or rare variants that had at least one significant predicted effect from our DVIM analysis we term as ‘impact variants’ (imVars). To avoid a high multiple hypothesis testing (MHT) burden, we focused on specific molecular effects in contexts related to T1D (immune cells and pancreatic tissue, N = 36).

**Figure 5.**
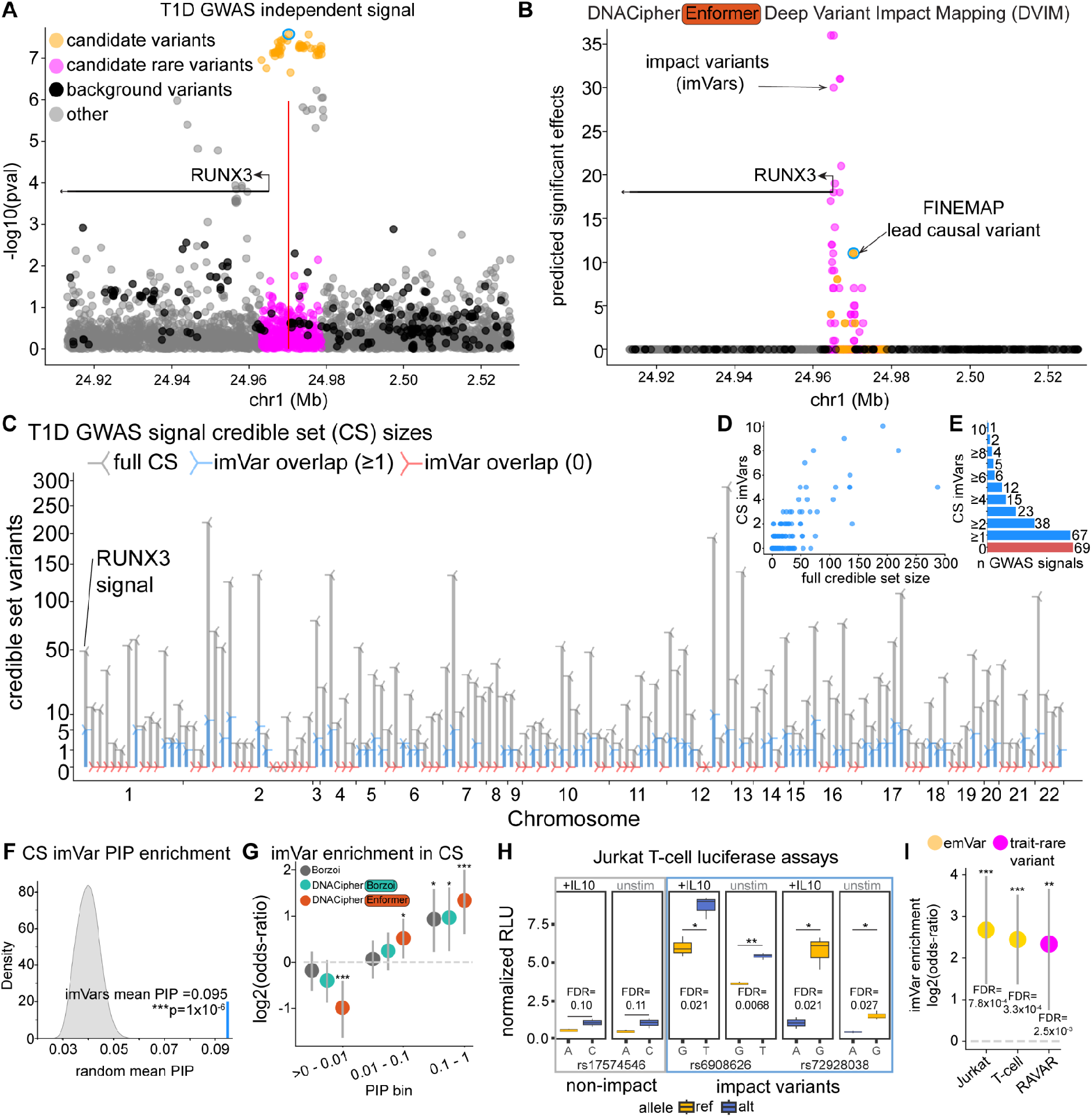
DNACipher Deep Variant Impact Mapping (DVIM) of Type 1 Diabetes (T1D) GWAS loci. **A.** GWAS associations with T1D (-log10 p-values, y-axis) for variants at the *RUNX3* locus (genome coordinates, x-axis). Variant categories are candidate common (trait-associated; orange), candidate rare (pink), background (non-trait associated; black), and other (gray) based on GWAS p-values, position and minor allele frequencies. The red vertical line marks the highest fine-mapping posterior inclusion probability (PIP) variant position. **B.** The number of significant predicted effects from DNACipher-Enformer DVIM (y-axis), which computes p-values from boot-strapping background variant predicted effects. The top-ranked common variant (blue outline) also had the highest PIP. Variants with significant predicted effects are termed impact variants (imVars). **C.** The y-axis is the CS size before (gray) and after (blue/red) imVar filtering. The x-axis is the genomic position of T1D GWAS signals. Red markers indicate imVars were not in the locus CS. **D.** Scatter plot of CS sizes before (x-axis) and after imVar filtering (y-axis). **E.** Ranked bar chart of N GWAS loci (x-axis) with N imVars in CS (y-axis). **F.** Distribution of mean PIPs from random CS sampling (grey; N = 1 x 10^6^) with observed mean PIP of CS imVars as a vertical blue line with empirical p-value. **G.** Log2-odds-ratio (y-axis) of overlap between common imVars and CS variants in different PIP bins (x-axis), with imVars called using either Borzoi, DNACipher-Borzoi, or DNACipher-Enformer predictions. The dotted line indicates the random expectation, and the error bars indicate 95% confidence intervals. Fisher’s exact test with FDR correction was used for statistical comparisons. Sample sizes are provided in Table S5. **H.** Box plots from Jurkat T cell luciferase assays for one non-imVar (grey box) and two imVars (blue box) with N = 3 replicates per condition. The y-axis is the normalized Relative Luciferase Unit (RLU) enhancer activity measurement, and the x-axis indicates the allele substitution for each variant. Tested conditions include T cell activation (+IL10 and CD3/CD28 Dynabeads) and un-stimulated. Box-plots display the median value, edges are 25th and 75th percentiles, and whiskers are 1.5X the interquartile range. Un-paired two-sided t-tests with Benjamini Hochberg false-discovery-rate (FDR) correction were used for statistical comparisons. **I.** Equivalent to G, except log2-odds-ratio (y-axis) are from overlap between imVars and expression modifying variants (emVars) from either Jurkat or T-cells, and rare imVars with rare trait-associated variants from the Rare Variant Association Repository (RAVAR). Sample sizes are provided in Table S5. ***FDR < 0.001, **FDR < 0.01, *FDR < 0.05.

We performed DVIM at all 136 T1D GWAS signals at 92 loci (Figure 5, Table S4). We highlight the *RUNX3* locus as an example application of DVIM (Figure 5A-B). 5 out of 49 (10.2%) of the candidate common variants associated with T1D were imVars at the *RUNX3* locus (Figure 5B), and the imVar with the highest number of predicted effects also had the highest causal PIP (Figure 5B). There were several more rare imVars but at approximately the same frequency given the total rare variants tested (N = 37 of 359, 10.3%). Rare imVars had more significant predicted variant effects than the common imVars (mean significant contexts N = 11.2 vs. N = 5.8), and rare imVars were concentrated at the *RUNX3* TSS (Figure 5B).

For this particular locus, we additionally performed a sensitivity analysis (Ext. Data Fig. 7A, Supp. Note 4), considering how the ranking of likely causal variants would change if we considered all possible molecular contexts from DNACipher or Borzoi compared with targeted analysis (Ext. Data Fig. 7B). We found using all contexts from DNACipher outperformed targeted analyses, although using all DNACipher contexts was highly similar to the targeted DNACipher analysis at the relevant MHT threshold (Ext. Data Fig. 7C).

Surprisingly, using all contexts from Borzoi reduced performance, and this was due to Borzoi largely ranking variants based on K562 and HEK293 effects, while DNACipher prioritized immune-related tissues consistent with the etiology of T1D (Ext. Data Fig. 7C,D). The imputation capability of the DNACipher model is hence highly favourable for the DVIM analysis, enabling prioritization of variant effects that do not reflect the ascertainment of the training data on potentially irrelevant cell lines.

Considering all 132,244 variants across the 136 T1D signals from our targeted DVIM analysis, we detected a similar number of loci with at least one common or rare imVar using DNACipher-Enformer, DNACipher-Borzoi, or Borzoi (63.4-70.2% and 96.3% of loci with common and rare imVars, respectively; Ext. Data Fig. 8A). We next examined the intersection of imVars from our targeted analysis with 95% credible set (CS) variants across all signals^38^ (Figure 5C-F, Ext. Data Fig. 8B-E), finding imVars within CSs at a similar number of loci across models (42.6-47.8% and 12.5-14.7% of loci with common and rare imVars, respectively; Ext. Data Fig. 8B). Taking imVars as a filter, the average credible set size could be reduced from 28.2 variants to 1.2 variants, with additional imVars called using Borzoi predictions (Figure 5C-D, Ext. Data Fig. 8C). For DNACipher-Enformer, 67 of 136 (49.2%) of the GWAS signals had imVars in the CS, with 29 (21%) of credible sets containing a single imVar and 52 credible sets (38.2%) containing 3 or less imVars (Figure 5E).

DNACipher-Borzoi and Borzoi had similar results (Ext. Data Fig. 8D). DNACipher-Enformer imVars within credible sets had significantly higher causal PIPs (mean PIP = 0.095) compared to randomly selecting the equivalent number of variants from the same credible sets (empirical p = 1 x 10^-6^) (Figure 5F). DNACipher-Borzoi imVars also gave significant mean causal PIPs (Ext. Data Fig. 8E), while Borzoi imVars were not statistically different from random sampling (Ext. Data Fig. 8E). We also considered enrichment of common imVars from the different models at different PIP bins, finding greater enrichment for the DNACipher-Enformer imVars than DNACipher-Borzoi, and then Borzoi (Figure 5G, Table S5). This indicates that DNACipher imVars are likely to be causal variants from statistical fine-mapping, and show greater correspondence to statistical fine-mapping than DVIM with Borzoi.

To test whether DNACipher accurately predicted the cell type context of molecular effects, we tested whether imVars for each molecular effect were enriched in accessible chromatin of the predicted cell type using independent single nucleus ATAC-seq (snATAC-seq) that was not within the DNACipher training data^38^. ImVars from DNACipher models were significantly enriched within accessible chromatin of the cell types predicted to be affected across molecular modalities, however for Borzoi this test was only significant for H3K27ac and RNA-seq due to missing experiments (Figure S13, Table S5). This demonstrates DNACipher provides context aware predictions across molecular modalities that are corroborated by independent snATAC-seq measurements.

We also experimentally validated the regulatory function of DNACipher-Enformer imVars, by performing luciferase assays in Jurkat T cells for two impact variants (rs6908626 and rs72928038 at the *BACH2* locus) and one non-impact variant (rs17574546 at the *RASGRP1* locus) (Figure 5H, Table S5). The sequences containing the imVars drove higher luciferase expression compared to non-imVar sequences (Figure 5H). In addition, both imVars had significant allele-specific differences (Figure 5H, Table S5). Consistent with these results, we tested for enrichment of imVars in expression modifying variants (emVars) determined from massively parallel reporter assays (MPRAs) of Jurkat T cells^39^ or primary T cells^40^, finding the strongest and most significant enrichment for both sets of emVars with DNACipher-Enformer imVars (Figure 5I) compared to DNACipher-Borzoi and Borzoi imVars (Ext. Data Fig. 8F).

### Rare imVars at T1D loci are enriched for rare variants associated with immunity traits

GWAS are underpowered to detect associations with rare variants^41^, and most rare variants are not included in CSs. Nonetheless, 17 of the 136 T1D GWAS signals (12.5%) contained rare (MAF < 5%) imVars in CSs (Ext. Data Fig. 8B). When all rare variants were considered, DNACipher-Enformer DVIM found 6,009 rare imVars (4.74% of the 126,742 rare variants tested) at 131 of the 136 T1D signals (96%) (Ext. Data Fig. 8B). Rare imVars had greater magnitudes of predicted effects compared with common imVars and non-imVars (Ext. Data Fig. 9), consistent with purifying selection that prevents large-effect alleles from reaching high population frequencies^42,43^. We next assessed whether rare imVars were previously trait-associated in the Rare Variant Association Repository (RAVAR)^44^. While only 30 of the 6,451 trait-associated RAVAR variants intersected with T1D GWAS loci (Table S5), 6 of these were rare imVars, which is significantly more than expected by chance (odds-ratio = 5, FDR = 2.5 x 10^-3^, Figure 5I, Table S5). The traits associated with rare imVars were related to immune cell abundance, such as platelet counts (rs754838420^45^ MAF=0.001, rs149765041^43,46^ MAF=0.02), monocyte counts (rs540639423^46,47^ MAF=0.008), and eosinophil/granulocyte counts (rs12722502^43,46^ MAF=0.02) (Table S5).

Autoimmune-diagnostic blood measurements such as IgA, IgM, ILR5, SLAM7 and Albumin protein measurements were also associated with one rare imVar (rs188468174^43,48^ MAF=0.01). Notably, DNACipher-Borzoi and Borzoi exhibited greater and more significant enrichments between rare imVars and trait-associated rare variants (Ext. Data Fig. 8F, Table S5). Overall, these results show that DNACipher DVIM can detect trait-relevant rare variants.

### Multi-trait DVIM prioritizes likely causal variants across thousands of GWAS loci

We applied DNACipher-Enformer with DVIM to whole-genome-sequence (WGS) GWAS of an additional 9 complex traits^49^, representing a total of 654 loci and 458,045 tested variants (Figure 6, Table S6). Tested variants included 33,564 trait-associated common variants (p < 5 × 10^−8^), and 424,481 rare variants within the same genomic regions. Variant effect predictions were also generated for 108,549 background variants. We tested for effects on RNA-seq (both strands) and chromatin accessibility for all 191 cell types represented in DNACipher (N = 573 molecular contexts). ImVars were detected across all traits analysed (1,858 common imVars, 24,552 rare imVars), and 8 of 9 complex traits (except Number of Children, NC) had imVars at > 90% of GWAS loci (Figure 6A). On average, 68% of loci for a given trait had either common or rare imVars, 25% had only rare imVars, and 1.3% had only common imVars (Figure 6A). Of the total tested variants per trait, an average of 5.2% were rare imVars, and 0.4% were common imVars (Figure 6B). For CS variants, an average of 5.5% were common imVars, and an average of 0.24% were rare imVars (Figure 6C). When considering the distribution of imVars across loci within each trait, we observe a relatively consistent frequency of common or rare imVars, with the means of rare imVars ranging from 3-5% of total tested variants per locus (median of means = 4.5%), and 0.0-0.33% for common imVars (median of means = 0.21%) (Figure 6D). Importantly, the difference between common and rare imVar frequencies per locus were largely mitigated if we considered the total number of common or rare variants tested (mean 0.0-4.8% common imVars per locus, median of means = 3.7%; mean 3.3-5.7% rare imVars per locus, median of means = 4.8%; Ext. Data Fig. 10A). CS variants were predominately common imVars (mean 0-4.9% common CS imVars per locus, median of means = 3.0%) with negligible rare imVars (mean 0% rare CS imVars), and this was consistent when considering the number of common or rare imVars tested (Ext. Data Fig. 10B).

**Figure 6.**
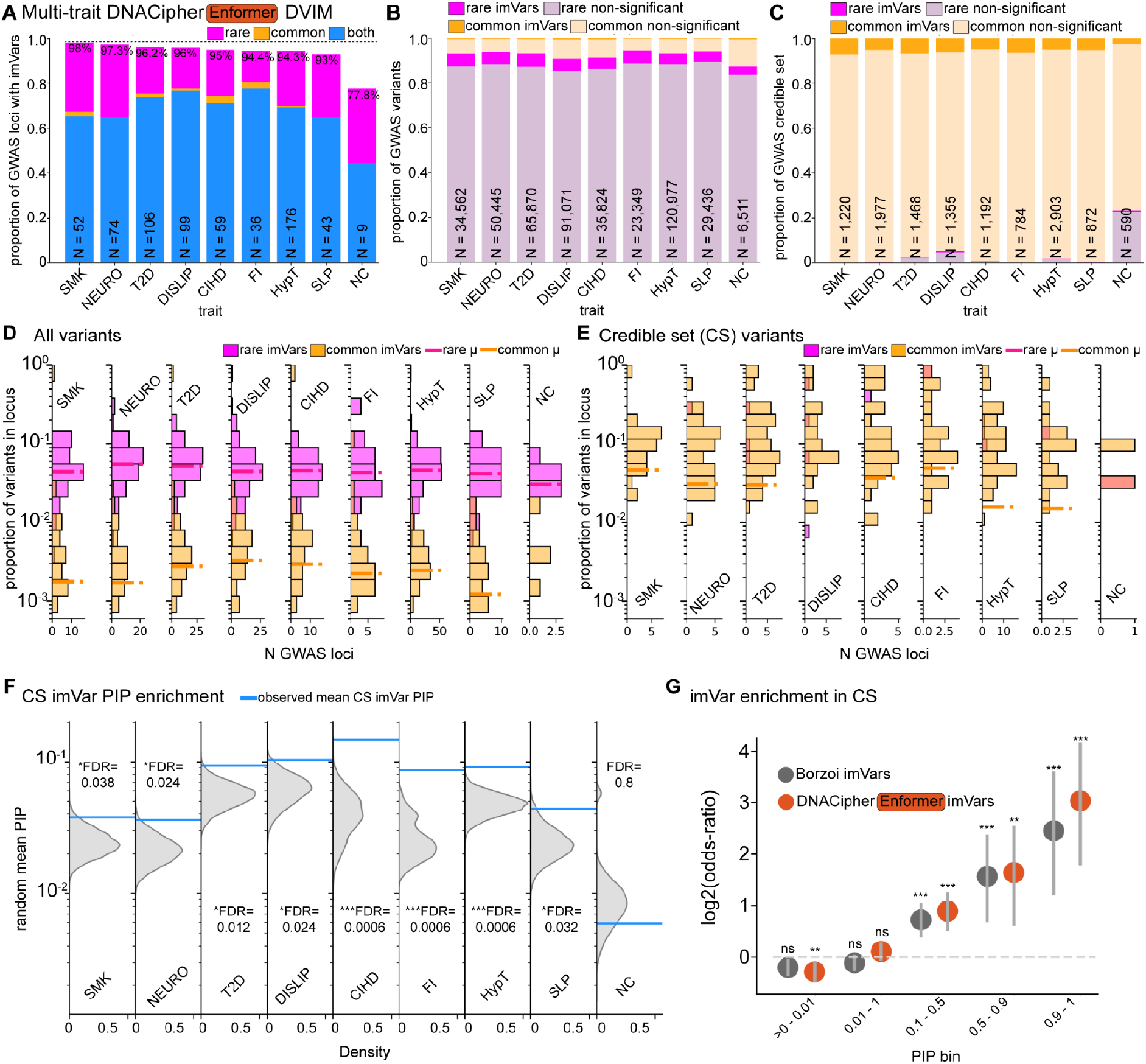
DNACipher Deep Variant Impact Mapping (DVIM) across 9 complex traits. **A.** Stacked bar chart with the proportion of GWAS loci (y-axis) across traits with at least one detected common and/or rare imVar (‘both’ indicates common and rare imVars detected) across 9 complex traits (x-axis). The number (N) of GWAS loci for each trait and total percentage of loci with ≥1 imVar are overlaid. Traits are abbreviated as SMK = ‘Ever smoked’, NEURO = ‘Neuroticism score’, T2D = ‘Type 2 diabetes’, DISLIP = ‘Dislipidaemia’, CIHD = ‘Chronic ischaemic heart disease’, FI = ‘Fluid intelligence score’, HypT = ‘Hypertension’, SLP = ‘Sleep duration’, NC = ‘Number of children’. **B.** Equivalent to A, except the y-axis indicates the total proportion of tested GWAS variants that were rare or common variants that were imVars or non-significant. N of the total variants tested at GWAS loci for each trait are overlaid. **C.** Equivalent to B, except for fine-mapping credible set (CS) GWAS variants. N indicates the number of CS DVIM tested variants for each trait. **D.** Histograms for each trait, showing the distribution across loci of the proportion of DVIM analysed variants that are rare or common imVars. Dashed lines indicate the mean imVar proportion of each type across loci. **E.** Equivalent to D, except subsetted to CS variants. **F.** Distributions of mean PIPs for random CS variants (N = 10,000) for each trait, with blue lines indicating the mean PIP for imVars. Benjamini-Hochberg false discovery rate (FDR) corrected empirical p-values indicate significance of mean imVar PIP. **G.** Log2-odds-ratio (y-axis) of overlap between common imVars and CS variants in different PIP bins from all traits (x-axis), with imVars called using either Borzoi or DNACipher-Enformer predictions. The dotted line indicates the random expectation, and the error bars indicate 95% confidence intervals. Fisher’s exact test with FDR correction was used for statistical comparisons. Sample sizes and p-values are provided in Table S7. ***FDR < 0.001, **FDR < 0.01, *FDR < 0.05.

For 8 of the 9 traits, imVars within CSs had significantly higher causal PIPs compared to randomly selecting the equivalent number of variants from the same CSs (Figure 6F). The trait without significantly higher imVar PIPs was NC, which had the lowest heritability and missing SNP-based heritability of the traits analysed^49^. Performing the DVIM analysis using Borzoi also showed non-significance for NC, but was additionally non-significant for 3 other traits (only 5 of 9 traits with significantly higher PIPs for CS imVars; Ext. Data Fig. 10C).

Comparing Borzoi and DNACipher-Enformer, both did have significant associations between common imVars and non-imVars within different PIP bins across all traits, with an apparent but non-significantly higher enrichment for DNACipher-Enformer (Figure 6G, Table S7). For traits with sufficient rare imVars within the RAVAR database (N = 4 traits), we found significant enrichment for rare trait-associated variants and DNACipher-Enformer rare imVars at hypertension (HypT) loci (Ext. Data Fig. 10D, Table S7). Borzoi rare imVars for HypT, chronic ischaemic heart disease (CIHD) and dislipidemia (DISLIP) were enriched for prior trait-associations (Ext. Data Fig. 10D, Table S7). Overall, applying DVIM to an additional 9 complex traits yielded similar results to our DVIM analysis of T1D, showing concordance between statistical fine-mapping and enrichment of rare imVars for prior trait-associated rare variants.

## Discussion

DNACipher DVIM identifies likely functional GWAS variants consistent with statistical fine-mapping while predicting their specific biological contexts and molecular effects across both common and rare variants. Crucially, DNACipher’s imputation resolves ascertainment bias in public genomics databases, preventing the accuracy drops that occur when S2F models would otherwise rely on experiments from mismatched cell types for molecular effect prediction.

By predicting functional impacts for rare variants, which are usually underpowered in GWAS, DNACipher DVIM addresses a major gap in human genetics. Although common variants account for most complex trait heritability^49,50^, rare variants constitute > 97% of all human genetic variants^51^, contribute substantially to missing heritability^49,50^, are important risk factors for common diseases^51^, and drive extreme disease phenotypes^43,52^. Across 10 complex traits, DVIM identified 30,561 rare imVars at GWAS loci, which is 4.7-fold more than previously catalogued trait-associated rare variants^44^. This lays the foundation for future high-throughput screening studies, such as MPRAs^10,40,53^, to validate functional predictions, and polygenic risk scoring (PRS) studies to utilize rare imVars to improve extreme phenotype prediction^43,52,54^.

By evaluating variants independently of linkage disequilibrium, DNACipher DVIM overcomes key limitations of observational genetics studies; estimation of causal variant frequency (common vs. rare)^55,56,57,58^, estimation of the number of causal variants per locus^59,54^, and pinpointing molecular mechanisms that could mediate trait-effects^32,60^.

Regarding causal variant frequency, this has historically sparked debate regarding the presence of potential ‘synthetic associations’ in GWAS studies^55^, which was refuted based on simulation showing trait-associated variants at GWAS loci would have a lower average MAF if they were driven primarily by rare variants^56^. Our results align with common causal variants, with most GWAS loci having common imVars detected, though we note almost all loci also had rare imVars detected, and a fraction of loci only had rare imVars. Loci without common imVars could indicate a lack of sensitivity for common causal variants, which is possible given a recent MPRA study found Enformer could be insensitive to common trait-associated emVars if these fell within low-information positions of motifs^61^. Furthermore, causal variants may have not been analysed, since they may not have been observed or imputed, and we did not consider structural variants or protein effects^62^.

In terms of estimating the number of causal variants, statistical fine-mapping accuracy depends heavily on prior assumptions, with infinitesimal models often outperforming strong assumptions of few causal variants^59^. While imVars were enriched in credible sets from fine-mapping techniques that assume few causal variants, we note DVIM significance calls are based on non-trait associated variants, and these could have under-powered small trait effects. Under an infinitesimal framework, virtually all variants altering a mediating molecular phenotype may exert non-zero trait effects, and therefore our approach likely calls high effect variants. Future work focusing on identifying these trait-mediating molecular effects could enable the identification of genetic variants with small effects that are under-powered with GWAS.

Observational eQTL mapping has had limited results for identifying GWAS-mediating molecular mechanisms^32,60^, with eQTL colocalization intersecting a median of only 21% of GWAS loci across traits^32^, though this can be improved by incorporating other types of molecular QTLs^63^. This has been attributed to a lack of power^60^ resulting in a bias toward genes that tolerate expression variation^32,64^. In contrast, DNACipher DVIM identified an imVar at > 90% of GWAS loci across 9 of 10 complex traits, indicating the DNACipher model predicted an outlier molecular effect relative to non-trait associated variants in most cases. Though additional experimental validation is needed, DNACipher DVIM offers a powerful framework to uncover potential trait-mediating molecular mechanisms that remain unexplained by conventional molecular QTL studies.

While prior sequence-to-function (S2F) advances focused on expanding the “receptive field” (input sequence length)^17,19,24^, DNACipher advances the “context field”, the range of predictable biological contexts. By combining long-range sequence embeddings with cell type/assay latent spaces, DNACipher evaluates long DNA sequences across tens of thousands of cell type/assay pairs. This scale surpasses previous models that are restricted to very short sequence contexts^31^, or lack the ability to represent DNA sequence directly^25^. Alternatives to DNACipher are more limited because they require cell type-specific inputs like gene expression^29,30,65^ or they can only make predictions for the same contexts that they are trained on^24^. Intriguingly, unlike smaller-scale imputation frameworks that generalize better across unobserved experiments than unobserved sequences^29,31^, DNACipher performs better on unobserved sequences than unobserved experiments. This shift likely reflects two factors: the vast scale of our experiment space (202 assays across 191 cell types) and strong sequence priors from pre-trained DNA embeddings.

Consequently, generalizing across the high-dimensional cell-type latent space presents a greater challenge than generalizing across sequences in our framework, which likely enabled the increase in the scale of our imputation approach.

In our molecular effect size prediction benchmarks, we did not observe a clear increase in performance for AlphaGenome compared to DNACipher or Borzoi. This could be because the benchmark in the AlphaGenome study considered binary classification of causal and matched non-causal eQTL variants, rather than the effect size predictions that we examined in this study. Another possible explanation is the contexts used to predict the eQTL effect sizes. AlphaGenome was trained directly on GTEx data, and where Borzoi lacked GTEx RNA-seq for a given tissue the AlphaGenome study averaged across tissues^24^, and Enformer was not directly trained on RNA-seq data at all^17^. In our benchmarks, we were able identify RNA-seq data matching the correct tissue type in Borzoi outputs (even if not GTEx RNA-seq data), and our DNACipher-Enformer model performed as well as the other models given training on relevant RNA-seq data. This raises the possibility that the improved eQTL detection for AlphaGenome reflects contexts used in the benchmark, rather than its longer sequence length. Consistent with this, DNACipher-Borzoi had higher signal prediction performance than DNACipher-Enformer, yet this did not translate into greater molecular effect size prediction accuracy. Further, using mis-matched tissue predictions significantly reduced the performance of AlphaGenome and Borzoi compared with the DNACipher models. Together, these observations raise the possibility that the task of predicting within-genome signals from within-genome sequence variation may be approaching a performance ceiling for genetic variant molecular effect prediction, such that further improvements in signal prediction performance do not necessarily translate into improved variant effect prediction. Determining whether such a performance ceiling exists, and whether tissue context alignment can explain the improvements in variant effect prediction in recent studies^24^, will require further systematic investigation.

While DNACipher on held-out experiments and sequences showed high signal prediction performance, there is still a performance gap compared to training experiments depending on assay type. We anticipate that the DNACipher model trained on new data as it becomes available will reduce this performance gap, since 1) there will be less data sparsity, and 2) the imputation will become more accurate with more information. Additionally, recent models^3466^ have made significant advances on training S2F models on single cell data, which when combined with DNACipher could improve predictions with a more comprehensive cell type embedding. The model can also be re-trained with richer sequence-embeddings from new open-source sequence models as they become available. Notably, there are still some assay types that DNACipher does not represent, such as chromatin contact maps and splicing, that are modeled by AlphaGenome^24^. Because of the specific output heads utilized to model these assay types, our framework does not currently represent them, and future work will be needed for imputation of these modalities. Given these possibilities, we envision continuous updating of DNACipher, which may yield a greater detection of imVars at GWAS loci.

Looking ahead, additional biological contexts—beyond cell type and assay—could be modeled as inputs to S2F-like frameworks, such as developmental stage or perturbation conditions. This would enable inference of how genetic variants influence phenotypes across development or in response to perturbation. Recent efforts have sought to impute missing molecular measurements during development^67^ and predict perturbation responses in unobserved cell types^68^. However, these models typically overlook DNA sequence as an input for variant effect prediction. Extending such models to incorporate DNA sequence could make it possible to pinpoint when, and under what conditions, genetic variants exert their effects. In this way, the high ascertainment bias of experimental genomics data could be overcome^69^, enabling the discovery of highly context-specific variants such as response eQTLs^70^.

## Methods

### ENCODE data preprocessing

We accessed the ENCODE^27^ data portal (https://www.encodeproject.org) in September of 2023, and obtained metadata for 748,860 processed human (*Homo sapiens*, GRCh38) files which had “released” status, no quality control issues, and which were derived from untreated biosamples. We restricted the dataset to biosample types “cell line”, “tissue”, or “primary cell”, thereby excluding *in-vitro* differentiated and cell free samples. We included the following assay types: “ATAC-seq”, “Histone ChIP-seq”, “Mint-ChIP-seq”, “TF ChIP-seq”, “DNase-seq”, “polyA minus RNA-seq”, “polyA plus RNA-seq”, and “total RNA-seq”. For RNA-seq assays, we retained files that provided strand-specific or overall “signal of unique reads” normalized by reads-per-million. If duplicate files existed for a given experiment, we selected the most recent file or the file with the largest size as a proxy for sequencing depth. For non RNA-seq assays, we used bigBed peak files. A single peak set per experiment was selected by prioritizing peaks files that aggregated the most replicates or that had the most recent release date. When files could not be prioritized with these criteria for a given experiment, we prioritized them by their “Output type” in the following order: “pseudoreplicated peaks”, “IDR thresholded peaks”, “replicated peaks”, and “optimal IDR thresholded peaks”. For DNAse-seq, “peak” files were selected. This data preprocessing methodology resulted in a single bigBed file per cell type-assay combination.

For ChIP-seq, ATAC-seq, and DNase-seq assays, we generated continuous “peak signal” profiles by taking the-log10(p-value) of each peak and assigning this value to every base pair within the genomic peak coordinate range. This approach reduced experimental background noise and enabled us to represent peaks as continuous signal tracks. When p-values were unavailable (e.g., DNase-seq or “IDR thresholded peaks”), we substituted the “signal value” instead. Next, we converted each track into bigWig format^71^, generating one “peak signal” file per cell type-assay combination.

### Train / test experiment allocation

Cell type labels were derived from the ENCODE “Biosample term name” metadata field, while assays were labeled according to the “Assay annotation” field. For ChIP-seq experiments we appended the target name (e.g., a specific histone modification or TF) to the assay label to ensure that each assay mapped to a clearly defined molecular target.

Following criteria similar to Avocado^25^, we only retained cell types that had ≥ 5 assays, and assays present in ≥ 3 cell types. For our initial assignment of train / test experiments for DNACipher utilizing Nucleotide Transformer^72^ (NT) embeddings (Supp. Note 1), for each assay, we designated the cell type represented in the most assays as the test cell type for that assay, to avoid depleting training data for cell types with extremely sparse experimental measurements. This selection process was iterative: once a test cell type was chosen for an assay, we updated the training set counts before proceeding to the next assay.

For the DNACipher models using sequence embeddings from models trained on genomic signal data (DNACipher-Borzoi and DNACipher-Enformer), we re-allocated train / test experiments to ensure the test experiments did not overlap with the training data of these models. We downloaded the train / test experiment metadata for Borzoi (https://raw.githubusercontent.com/calico/borzoi/main/examples/targets_human.txt), and since we only utilized ENCODE data, we removed any Borzoi training experiments with matching ENCODE accession IDs as potential test experiments. We then re-formatted the cell type and assay metadata to match our cell type/assay labels for each experiment, and removed from potential test experiments any experiments that matched these labels, yielding N = 1,060 experiments non-overlapping with the Borzoi training data. From these potential test experiments, for each assay, we designated the cell type with the *least* number of other assays measured as the test cell type for that assay, to ensure our performance metrics reflected sparsely sampled cell types as is the most common case (Figure S2). This selection process was iterative: once a test cell type was chosen for an assay, we updated the training set counts before proceeding to the next assay. To avoid depleting training data for assays or cell types with extremely limited experimental measurements, we restricted allocating an experiment to testing unless either the cell type or assay had at least 3 remaining experiments present in the training set. We repeated this process of train / test allocation 5 times with no overlap between test experiment allocations within each fold.

Because of the constraints on allocating test experiments to control for pre-training data leakage and ensure adequate training data, for each fold we could not allocate a unique test experiment for every assay, such that fold-1 had 123 assays of 202 with a test experiment, fold-2 had 93, fold-3 had 65, fold-4 had 45, and fold-5 35 assays with a test experiment.

### Signal normalization

For each train / test region (see section “**Train / test sequence allocation**” for details), we extracted the average signal of the central 294 bp using bwtool^73^. We then applied the arcsinh transformation to mitigate the influence of outliers, similar to Avocado^25^. All normalization parameters were derived exclusively from the training sequences and experiments to prevent data leakage, and the same normalization steps were then applied to the test sequences and experiments. For RNA-seq assays, we first applied a log2-transformation to non-zero signals, then subtracted the minimum log2 signal value to ensure all signals were positive. Across all assays, we then computed the mean non-zero signal for each assay across all training cell types and training regions. From these per-assay means (µ *_i_*), we calculated the median of those means (*m*). Using this median as a reference, we defined a scaling factor (*s _i_*) for each assay *i* as per equation 1.

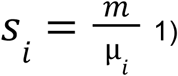

Multiplying each assay’s signals by *s_i_*, we aligned their average magnitudes, resulting in a uniform mean non-zero signal across assays (Figure S15). This normalization approach allowed for more direct comparisons of signals from diverse assays and cell types, and was essential to minimise systematic scaling differences between assays that otherwise dominated the loss function.

### Train / test sequence allocation

To define genome regions for training and testing, we used bedtools^74^ to tile the hg38 reference genome into 5,994 bp windows, with a 1,998 bp step size to obtain 3-fold coverage. This window size matches the 5,994 bp input length for NT embeddings that were used for the initial DNACipher models (Supp. Note 1). We excluded regions overlapping the ENCODE blacklist^75^. For the initial DNACipher models that utilized NT embeddings, chromosomes 2 and 16 were allocated as test sequences (N = 163,745), leaving a total of 1,344,175 train sequences. For details on how sequence embeddings for NT were for these sequences, see Methods section “**Nucleotide Transformer (NT) embeddings”**.

For the DNACipher models that utilized long sequence embeddings (Borzoi or Enformer), train / test sequences were allocated by intersecting the tiled regions with the train / test sequences allocated with Borzoi (https://github.com/calico/borzoi/blob/main/data/sequences_human.bed.gz). Borzoi used 5-fold cross validation, where the train sequences were re-allocated for each fold but the test sequences were constant between folds. We therefore utilized the tiled regions that intersected the Borzoi test regions as our test sequences (N = 345,371) and allocated all of the tiled regions that overlapped any of the Borzoi train regions across folds as our train sequences (N = 1,162,549). For details on how sequence embeddings were generated for these sequences, see Methods section “**Long sequence embeddings”**.

For the alternative DNACipher models which utilized genome location on model input (Supp. Note 1,2), we utilized a different train / test region allocation, to avoid leakage between train and test sets due to intersecting sequences (which were originally tiled at ⅓ overlaps across the genome). For these models, the train and test regions were generated by tiling at 5,994 bp across the genome using bedtools^74^ with a 5,994 bp step size. Within each genome location encoded at 12 kb resolution on chr2 and chr16, the first 6 kb sequence was allocated to the train set and the second sequence was allocated to the test set (Supp. Note 1,2). This resulted in a total of 54,440 test sequences and 1,399,036 train sequences for the genome-location encoding models.

### Long sequence embeddings

We extracted Enformer and Borzoi embeddings by tiling across the hg38 reference genome with a window size matching the model sequence input (196,608 bp for Enformer, 524,288 bp for Borzoi) and a step size matching the output embedding window in the centre of the input sequence (114,688 bp for Enformer, 196,608 bp for Borzoi)^17^. For each tile, we extracted the corresponding reference genome sequence, and passed the sequence through the corresponding model to extract the latent features from the layer immediately prior to the output heads. We introduced “N” padding at chromosome boundaries to maintain the input sequence lengths for tiles at chromosome boundaries. For Enformer, this resulted in 896 x 3,072 embedding per sequence, where the 896 dimension refers to 128 bp sequence bins, and 3,072 represents the latent feature dimension. For Borzoi, this resulted in a 6,144 x 1,920 embedding, where the 6,144 dimension refers to 32 bp sequence bins, and 1,920 represents the latent feature dimension. To generate sequence embeddings for each train and test region, the sequence embeddings at bins that intersected the genome regions were averaged (295 bp regions), to generate 3,072 and 1,920 latent sequence features per region for Enformer and Borzoi, respectively. We used the Pytorch-based implementations for both models to extract sequence embeddings^76^.

### Nucleotide Transformer (NT) embeddings

We obtained Nucleotide Transformer (NT) embeddings from the last layer (layer 24) by passing the full 5,994bp train / test region through the hg38 trained, 500 million parameter model^72^. This produced a 999 × 1,280 feature matrix per region, where the 999 dimension represents 6 bp sequence bins, and 1,280 denotes the latent sequence feature dimension. To obtain a single embedding, we averaged the middle 48 rows of this matrix (representing the central 288bp of the region) generating a 1,280-dimensional sequence embedding per region.

### DNACipher model architectures

We implemented the DNACipher model in Pytorch^77^, resulting in approximately 1.3 million parameters. DNACipher has 3 input layers: (1) a cell type embedding layer (input size = 191, output size = 128), (2) an assay embedding layer (input size = 202, output size = 256), and (3) a fully-connected layer that projects embeddings (3,072 for Enformer, 1,920 for Borzoi, 1,280 for NT) down to 128 dimensions. The outputs from the 3 input layers (cell type = 128, assay = 256, sequence = 128) are concatenated to one tensor of size 512, and then passed through 3 fully-connected layers with 512 input sizes and 512 output sizes. Layer normalization was applied after each of the deep layers^78^. All fully connected linear layers included learnable bias parameters. Gelu was used as the activation function for all fully connected linear layers. The final output layer projects to a single continuous signal value.

For models that included genome location, there were an additional 3 input embedding layers to encode genome location at different resolutions: 11,988 bp, 47,952 bp, and 119,880 bp. At each bin size, the bin within which the corresponding input sequence occurred was one-hot encoded and passed into the model at the corresponding input layer. Bin size 11,988 bp had 257,628 bins, bin size 47,952 bp had 64,417 bins, and the largest bin size of 119,880 bp had 25,772 bins across the hg38 genome. Each genome location input layer was projected to 64 dimensions. Models utilizing NT embeddings and genome location had 23.3 million parameters, models with NT embeddings and without genome location had 1 million parameters, and models with genome location and no sequence input had 23.14 million parameters (Supp. Note 2).

### Training strategy

Training batches consisted of 256 regions with the full set of signals across training experiments. We utilized Adam as the optimizer^79^ with a fixed learning rate of 0.0014526, which was determined from hyper-parameter testing of different learning rates ranging from 0.001 to 0.01 using RayTune^80^. For regularisation, we used a 5% dropout rate across all layers^81^. DNACipher models that included genome location were prone to overfitting and therefore a more stringent dropout regularization^81^ was employed specifically at the genome location input layers at a rate of 20%. For the initial DNACipher models utilizing NT embeddings, training was conducted for 25 epochs with MSE as the loss function and without ReLU activation applied to the model output, followed by a secondary round of training for an additional 25 epochs with ReLU activation on the model output and where each experiment was weighted in the loss function. The ReLU activation was to enforce non-negativity. For the long-sequence models, we updated this approach to use SoftPlus activation instead of ReLU, to avoid 0 gradients for negative values that occur with ReLU. Experiments representing rarer cell types or assays were more heavily weighted for the secondary round of training. In this case, the squared error between the predicted and observed signals across the training experiments was multiplied by the weight assigned to each experiment. The mean of the weighted squared error was the loss for this second round of training.

The weight of each experiment was determined using a sampling strategy that aimed to reduce the inherent bias in the ENCODE dataset due to assays which were overrepresented across cell types (e.g. DNase-seq) and cell types which were overrepresented across assays (e.g. K562 cells) (Figure 1F). To achieve a uniform distribution of experiments for each cell type would require over-sampling some assays.

Similarly, to achieve a uniform distribution of experiments for each assay would require oversampling some cell types. To reduce the influence of this ascertainment bias on the loss, we empirically tested a range of assay (*p_assay*) and cell type (*p_cell = 1 - p_assay*) sampling strategies. For a given assay bias (*p_assay*), we conducted 100,000 samplings of the available experiments. At each sampling, we chose whether to prioritize reducing assay bias or cell type bias with probability *p_assay* or *p_cell*, respectively. If reducing assay bias was selected, a random assay was selected with a probability inversely proportional to its frequency within the dataset. Of the available cell types for the chosen assay, a cell type was then chosen with probability inversely proportional to the cell type’s frequency within the dataset. Conversely, if reducing cell type bias was selected for the sampling, the cell type was chosen first, and then the assay. The number of times a given experiment was chosen using this sampling process was counted. For different values of *p_assay* chosen uniformly between 0 and 1, the bias of chosen assays and cell types was quantified using the standard deviation of cell type counts across assays (assay bias) and assay counts across cell types (cell type bias), revealing an optimal *p_assay=0.428* that minimized both the cell type and assay bias (Figure S16). The sampling count of each training experiment *c_i_* was then used to determine the weight of the experiment (*w_i_*) as per equation 2.

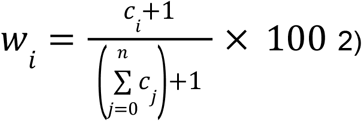

Where *n* is the total number of training experiments.

### kNN baseline comparison

To assess DNACipher’s predictive accuracy relative to a simple baseline, we compared its prediction against a mean-based estimate across kNN cell types for a given assay. We did not use the observed signal values for the ‘missing’ experiment to define neighbors because this would require using data that is supposed to be unobserved, resulting in data leakage. Therefore, we identified using OpenAI text embeddings for the cell type / tissue names. This semantic definition of neighbours aligns with how a user may reasonably use predictions from other tissues for an assay if the relevant tissue is missing. Specifically, we computed 1,536 text embeddings by inputting the cell type/tissue string into the OpenAI API with the ‘text-embedding-3-small’ model, and identified the 10-nearest neighbor cell types based on cosine distance in this embedding space. We found these neighbors to be reasonable. For example ‘T-cells’ had ‘T-helper 17 cell’, ‘CD4-positive, alpha-beta T cell’, ‘CD8-positive, alpha-beta T cell’ within the top 10 nearest neighbours; 8 of the 10 nearest neighbours were T-cell subtypes and the remaining 2 neighbours were B-cell subtypes. The nearest neighbours for each cell/tissue type are provided in Table S1.

To compare DNACipher to the kNN baseline statistically, we used evaluation batches of 256 train or test sequences, ran DNACipher predictions for these, and then generated Pearson correlations with the observed signals for train or test cell types for each batch (N = 1,350 batches, N = 345,600 train or test sequences). We also computed equivalent correlations for the kNN baseline, where we used the mean signal for a given sequence across the k-nearest neighbour cell types for a given assay as the prediction. Because of missing experimental data, the mean was only taken for the cell types that had a measurement for that particular assay, and therefore at most 10 neighbors were used. We then used a paired two-sided Wilcoxon rank-sum test to compare DNACipher predictions and the kNN baseline in terms of their Pearson’s correlations across batches for a given assay, followed by Benjamini-Hochberg FDR correction for multiple hypothesis testing across assays. This was performed for both DNACipher-Enformer and DNACipher-Borzoi across all 5-folds of train / test experiment allocation.

### Cell type specific signal prediction

To select test sequences with cell type specific signals across assays represented for a given cell type, we first summed the signals for all available assays for a given cell type across all test sequences (N = 345,371). We then calculated the sum and standard deviation across cell types for each test region, and took all regions with a sum greater than 150 and less than 1500, since this range had an approximately linear magnitude/variance relationship. Ordinary least square regression (statsmodels.regression.linear_model.OLS) was then applied to regress the observed sums to the standard deviation across test regions, and we selected outliers as those with a residual variance greater than 0.5 (N = 11,016). From these test regions with a higher variability than expected given the overall signal magnitude, we then selected regions with high cell type specificity. Specifically, we constructed a one-hot-encoded vector for each cell type, and took the top 100 test regions with signals across cell types that had the smallest cosine distance to the one-hot-encoding of each respective cell type. Because some cell types did not have unique signals, but shared some signals with others, this resulted in a unique set of 162 test regions with the highest cell type specificity (Fig. S5A). We then used DNACipher-Borzoi to impute these test region signals across the same set of assays for each cell type, and also took the sum across these assays to get a single signal per cell type (Fig. S5B), as equivalently performed for the observed data (Fig. S5A). We then calculated Spearman (scipy.stats.spearmanr) and Pearson’s (scipy.stats.pearsonr) correlations between the predicted and observed signals across the test regions for each cell type, taking significant correlations as cases where both correlation statistics were significant after Benjamini-Hochberge false-discovery-rate (FDR) correction (FDR < 0.05).

### Latent embedding annotation

We obtained latent representations of cell types and assays directly from DNACipher’s input layers. We passed one-hot encoded vectors through their respective embedding layers and extracted the resulting latent features (128 dimensions for cell type and 256 for assay embeddings). We used Scanpy^82^ to apply UMAP to these latent representations, constructing k-nearest-neighbour graphs with cosine distance. For cell type and assay embeddings, we used k = 3, while for sequence embeddings we used k = 15. After dimensionality reduction with UMAP, we applied Leiden clustering^83^ to identify assay clusters using default parameters. Clusters were annotated by examining the biological themes of the grouped assays and cell types, with the exact memberships of each cluster shown in Table S2.

### GTEx eQTL effect size benchmark

Significant eQTLs and DAPG fine-mapping summary statistics for GTEx version 8 were downloaded from the data portal^23^ (https://gtexportal.org/home/). We selected all variants with a fine-mapping PIP > 0.99 and <35 kb from the eGene TSS for the associated eGene across all 49 GTEx tissues as likely causal eQTL variants. We additionally filtered for eQTL-eGene-Tissue combinations with absolute effect sizes >0.47 to reduce the total. We matched the GTEx tissues to the tissues that were predictable by the benchmarked models. AlphaGenome was trained on the GTEx tissues except for lung tissue, and so we used predictions corresponding to these outputs but excluded lung predictions. For DNACipher, we manually matched the tissues (e.g. matching GTEx ‘Pancreas’ with ‘pancreas’ as an ENCODE biosample). For Borzoi, matching was first established using semantic similarity, followed by manual curation to ensure precise ontology matching, and calling five most similar tissue types to the corresponding GTEx tissue. For Borzoi, if RNA-seq was not present for a given match tissue type, since these were ranked by similarity to the GTEx tissue, we used the subsequent tissue match preference and were able to making tissue matches predictions for all eQTL-eGene-tissue triplets. Exact output predictions used for each model and the corresponding GTEx tissue are provided in Table S3. For DNACipher, we used plus strand polyA+ RNA-seq and minus-strand polyA+ RNA-seq as the assays. Effect sizes were predicted from the models by comparing the reference and alternative allele sequence predictions for the corresponding output, with the query sequences centred on the eQTL. To ensure our predictions were specific for the eGene, we only considered strand-specific and output signal regions that intersected the eGene location and for the strand-specific RNA-seq predictions that corresponded to the strand the gene was encoded on. To score the effect of the eQTL on the eGene, we used the fold-change between the alternative and reference sequence predictions from each model for the relevant output signals (sum(alt)-sum(ref)). We then z-scored these signal predictions across all predictions for a given model, in order to enable a consistent cutoff for considering non-zero signal prediction correlations.

### Transcription factor allele-specific binding (TF ASB) effect size benchmark

TF ASB events were downloaded from the ADASTRA^35^ database (https://adastra.autosome.org/). ADASTRA’s raw cell-type labels were aligned to DNACipher’s label set. Each raw label was compared with every DNACipher label using the cosine similarity of the 768-dimensional embedding produced by Google’s gemini embedding model (gemini-embedding-2-preview, accessed April 2026)^84^. The 5 most similar DNACipher labels for each ADASTRA label were manually curated to a single match. In total, 189 of the 603 labels were mapped and ADASTRA records whose labels could not be mapped were excluded. Because alleles that intersect ASB events may not be causative, we filtered for events that had concordant changes between the ASB effect size and the position weight matrix score for the TF. We took TF ASB effect sizes as the difference between the mean alternate allele specific effect size and the mean reference allele specific effect size.

We then matched the corresponding TF outputs from each model with the listed TFs for each ASB event, and likewise for the tissue types, with the exact outputs used for each model listed in Table S3. In cases of missing tissues for AlphaGenome and Borzoi, a random other tissue with ChIP-seq output for the corresponding TF was used. We measured the variant effect as the difference between the alternative and reference sequence prediction, with the sequence centred on the variant position, and the signal prediction that overlapped the variant location. This corresponded to the 128 bp signal prediction that overlapped the variant position for DNACipher-Enformer and AlphaGenome, and 32 bp signal prediction for Borzoi. We then correlated the model predictions for each tissue-specific and tissue-mismatched TF ASB prediction against the observed TF ASB effect sizes.

### dsQTL effect size benchmark

Summary statistics for the LCL dsQTLs were the same as used by ChromBPNet^36,85,86^ and were specifically called using DNAse-seq data from Yoruba LCLs (https://static-content.springer.com/esm/art%3A10.1038%2Fng.3331/MediaObjects/41588_2015_BFng3331_MOESM26_ESM.xlsx). We followed the same pre-processing as previously described^36^. Because coordinates were reported in hg19 and the AlphaGenome API does not support changing reference genomes, we used pyLiftOver to convert to hg38 coordinates and used hg38 as the reference genome. For the DNACipher models and AlphaGenome, we used outputs corresponding to ATAC-seq and GM12787, which is a common LCL. Borzoi lacked ATAC-seq for the same cell line, and so we used the corresponding DNAse-seq output. For AlphaGenome and Borzoi, we also simulated missing this context, by taking a random other tissue as the dsQTL prediction to measure the effect of this on the dsQTL effect size prediction. All dsPeak reported were 100 bp in length, and all dsQTLs were within 50 bp from the dsPeak start or end position. Therefore we measured variant effects as the alternative sequence predictions minus the reference sequence predictions for the relevant LCL output for each model at signal predictions +/-150 bp from the variant position.

### dsQTL saturated *in silico* mutagenesis (ISM)

We performed attribution analysis based on a saturated in silico mutagenesis of 50 bp sequences (hg38) surrounding each of the 574 LCL dsQTLs. For each variant generated, effect size was measured across the total output window (114 kb) for DNACipher-Enformer with ATAC-seq as the assay and GM12787 as the cell type. Variant effects were then summed at each position across the 50 bp sequences, and tangermeme^37^ was used to call high attribution ‘seqlets’ between 4 and 25 bp (default values). This analysis identified at least one seqlet for 551 of the 574 dsQTL sequences (96%) (Figure S10A). These seqlets were concentrated at the position of the dsQTL within sequences (Figure S10B), with 377 of the 574 dsQTL positions (66%) intersecting with at least one seqlet in the sequence. 220 (53.6%) seqlets intersected dsQTLs, and motif matching was performed on these against non-redundant core JASPAR TF motifs^87^ with tangermeme^37^. 219 TFs matched to at least one seqlet (Figure S10C). For the top 10 TFs with the highest number of dsQTL seqlet matches, we reconstructed position frequency matrices from the seqlets, ensuring to match the strand with the highest TF match of the seqlet (Figure S10D).

### GTEx likely causal and non-causal eQTL molecular effect detection

GTEx^23^ variants were stratified by tissue and distance from the TSS of the associated eGene in bins of 100 kb. Within each stratification, the top and bottom 3 variants with the highest posterior inclusion probability (PIP) which had a PIP > 0.9 or PIP < 0.1 were chosen as example likely causal and likely non-causal variants, respectively (Figure S9A). This resulted in an equal representation of variants across TSS distances and tissues, with a range of effect sizes (Figure S9B-D).

Selected eQTLs were scored by the sum of absolute signal prediction difference between predictions from the alternative and reference sequences centred at each eQTL location, with sequences corresponding to the input lengths for the corresponding model; 1,000,000 bp for AlphaGenome, 524,288 bp for DNACipher-Borzoi and Borzoi, 196,608bp for DNACipher-Enformer and Enformer, and 1,000 bp for DeepCT. The experiments scored corresponded to the full set of assay predictions for biosamples matching the corresponding GTEx tissue type. For AlphaGenome, Borzoi, and Enformer, since cell/tissue labels could across amongst assays but still be relevant for the given GTEx tissue, we determined the top 5 most relevant cell / tissue labels present across the experiments and used all assays that matched these labels for the corresponding GTEx tissue. If a given assay was present for multiple relevant tissues, the tissue most semantically similar was used. All of the matching biosamples used are listed in Table S3. Within each assay type, tissue-specific causal-and non-causal eQTL predicted effects were then compared using a two-side independent t-test.

P-values across assay types were then adjusted using the Benajamini-Hochberg false-discovery-rate correction^88^. Assays with an adjusted p-value < 0.05 were taken as having significantly different predicted effects between causal-and non-causal eQTLs for each model.

### Deep Variant Impact Mapping variant allocation

For T1D, we downloaded summary statistics of variant associations, FINEMAP posterior probabilities of variant causality, and variant chromatin accessibility peak annotations from the supplemental data provided by Chiou, *et al.*^38^. For the additional 9 complex traits, the SuSIE-SS^2^ fine-mapping summary statistics were downloaded from the supplemental data^49^, and the authors provided the full summary statistics for all variant-trait associations tested from the WGS genotypes. When allocating variants to each category (candidate common, rare, background) we analysed each independent signal determined by fine-mapping for each respective study, such that a given variant may be re-analysed if within the sequence window of more than one independent signal. Variants were stratified as background, candidate common variants, candidate rare variants, and other rare variants. Variants within ±57 kb from the GWAS signal center were considered relevant for testing, since this falls within the 114 kb range where signal predictions can be made with DNACipher-Enformer. GWAS signal center was defined as the lead causal variant position from fine-mapping. To analyse a broader set of common variants, we considered ‘candidate common variants’ as those with a nominal p-value of association an order of magnitude greater than p-value used to nominate the GWAS locus in the original studies. In particular, for T1D p < 5 × 10^−8^ was used to determine GWAS loci, and so we considered candidate common variants as those with p < 5 × 10^−7^. For the other 9 complex traits, p < 5 × 10^−9^ was used by the original study, and so we used p < 5 × 10^−8^ to select candidate common variants. Candidate rare variants were taken as any variant with MAF < 5% that were within the same genomic region as the selected candidate common variants (defined as the minimum and maximum positions of the selected candidate common variants). Background variants were variants which did not meet the criteria of candidate common or rare variants, that were non-significant (nominal p-value > 0.001), and had a MAF > 5%. Because these initial criteria resulted in some of the GWAS signals having few background variants, if less than 100 background variants were selected with these criteria, the top 100 non-significant variants with the highest MAFs were used as the background variants instead.

### Deep Variant Impact Mapping statistical testing

For each of the variants selected (candidate common, rare, and background variants), we used DNACipher-Enformer to predict the effect of the variants for T cells, B cells, natural killer cells, and pancreas. Predictions were made for a total of 9 assays, which included minus strand polyA plus RNA-seq, plus strand polyA plus RNA-seq, minus strand total RNA-seq, plus strand total RNA-seq, ATAC-seq, H3K27ac ChIP-seq, H2BK20ac ChIP-seq, H3K27me3 ChIP-seq, and CTCF ChIP-seq. Variant effects were measured similarly to the GTEx eQTL molecular effect detection benchmark, except the overall effect was signed by the overall change of the locus (signed sum of absolute differences). To ensure the variant effects at each GWAS signal were comparable, variants associated with each GWAS signal had fixed query sequence locations so that each variant was compared against the same reference sequence. Hg38 was used as the reference genome for both the T1D and multi-trait analysis. The reference sequence for each signal was fixed to the centre of the GWAS signal. Because the T1D study utilized hg19 mapped variants and lifted over all coordinates to hg38, we replaced the reference allele with the corresponding hg19 nucleotide in the cases where the hg19 and hg38 reference sequences did not match. For the T1D DVIM analysis, we predicted variant effects for 36 different cell type and assay combinations for each of the 197,148 variants across 136 T1D GWAS signals, resulting in 7.097 million variant effect predictions. For the multi-trait DVIM, we predicted across all cell types (N = 191) for plus/minus strand total RNA-seq and ATAC-seq (N = 3 assays), making 573 effect predictions per variant and 324.66 million variant effect predictions for the 566,594 variants analysed (with some duplication due to variant sharing across independent GWAS signals).

Within each T1D GWAS signal, we tested whether the predicted effect of each candidate (common or rare) variant differed significantly from a null distribution of variant effects. To account for uncertainty in the estimated null distribution due to a limited number of background variants^89,90^, we performed bootstrap resampling of the background variant effects.

For each molecular context—defined as a cell type–assay combination *m*—we resampled the predicted effects of background variants *B* times (*B* = 1,000,000). In each bootstrap iteration *b*, we calculated the mean (*μ*_bm_) and standard deviation (σ_bm_) of the resampled background effects, enforcing a minimum standard deviation of 0.01 to prevent spurious significance from zero-variance. We then counted the number of bootstraps in which each candidate variant’s predicted effect was not statistically significant (*N_vm_*) when compared to the bootstrapped null using a two-sided one-sample z-test (equation 3).

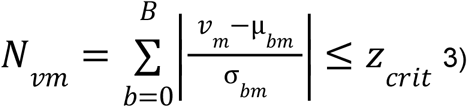

The critical z-score threshold (*z_crit_*) was derived using a Bonferroni correction for the total number of variant effects tested within the GWAS signal (equation 4).

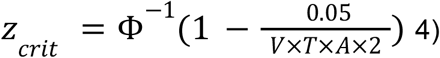

Where ɸ^−1^ is the inverse cumulative density function of the normal distribution, 0.05 is the un-adjusted critical p-value, *V* is the total number of variants tested at the GWAS signal, *T* is the total number of cell types tested, *A* is the total number of assays tested, and the 2 is because it is a two-sided test.

We then estimated a p-value per variant effect that considered the uncertainty in the null distribution^89,90^ (*p_vm_*), as the proportion of times a given candidate variant effect (*vm*) was non-significant across the boot-strapped null distributions (equation 5, pseudocounts added to prevent 0 p-values).

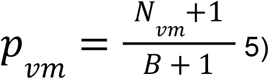

To determine the number of GWAS signals with imVars, while accounting for multiple hypothesis testing, we took a hierarchical approach typical for eQTL studies^91^. For each GWAS signal, we recorded the minimum observed Bonferroni-corrected p-value as the p-value for that signal, and then adjusted p-values across GWAS signals using the Benjamini-Hochberg (BH) False Discovery Rate method. GWAS signals with an adjusted BH p-value less than 0.05 were considered as having at least one imVar. This was repeated for rare and common variants separately (Figure 5,6), and also for variants that intersected the credible set variants (Figure 5C), to determine the number of signals containing imVars of each type. For details regarding the DVIM sensitivity analysis, see Supp. Note 4. In addition to DVIM with DNACipher-Enformer predictions, we also performed DVIM using Borzoi with the equivalent targeted outputs, and additionally DNACipher-Borzoi for the T1D analysis. Between these analyses, the exact same variants were considered.

### Deep Variant Impact Mapping comparison with statistical fine mapping

After calling imVars at each GWAS signal, we intersected them with credible set variants for each signal. To determine if the imVars were more likely to be causal, we calculated an empirical p-value for the fine-mapping PIP for the imVars. We randomly sampled the same number of credible set variants from each GWAS signal as the number of called imVars at each signal, and took the mean PIP of these randomly selected variants 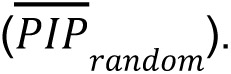 We repeated this 1,000,000 times (*R* = 1,000,000) for the T1D analysis to generate a random background distribution of mean PIPs, and R = 10,000 times for each of the traits in the multi-trait DVIM analysis. We used more random selections for T1D, because at R = 10,000 we did not observed any cases of the 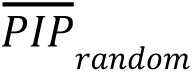 exceeding the mean imVar PIP, and so we increased the sampling to get a more accurate p-value. We calculated the empirical p-value as the number of random mean PIP values greater than or equal to the observed mean PIP (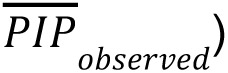 of the imVars, divided by the number of random selections performed (*R*) (equation 6).

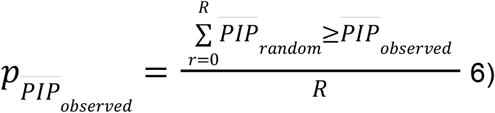

In addition to determining if the CS imVars had higher PIPs than expected, we also considered binary enrichment of imVars within CSs. We constructed contingency tables of imVars intersected with CS variants within bins of causal PIP. A Fisher’s Exact Test to calculate p-values for non-random association between the imVars from each model and PIP bins was performed, with odds-ratios with two-sided 95% confidence intervals calculated using Scipy. Benjamini-Hochberg FDR was applied for multiple hypothesis testing. The exact counts for the contingency tables are provided in Table S5 for the T1D analysis, and Table S7 for the multi-trait analysis.

### Deep Variant Impact Mapping comparison with independent snATAC-seq data

To evaluate whether the predicted cell type-specific variant effects were consistent with functional elements in the same cell types, we examined the overlap between imVars and chromatin accessibility annotations from Chiou *et al.*^38^. These variant annotations included cell types determined from PBMCs and pancreatic snATAC-seq. Because DNACipher does not represent the cell types at the same level of resolution as the snATAC-seq data, we collapsed cell types into broader categories including pancreas, T cells, B cells, and natural killer cells.

We then constructed contingency tables to represent the intersection between predicted variant effects and overlap with accessible chromatin in the same cell types. The contingency tables were constructed across all variants at all GWAS signals, counting the number of variants in the following categories: accessible in cell type and predicted to affect the cell type by the given assay, accessible in cell type and not predicted to affect the cell type by the given assay, not accessible and predicted to affect the cell type by the given assay, and not accessible in the cell type and not predicted to affect the cell type by the given assay (Table S5). We then performed Fisher’s exact test to determine if variants predicted to affect particular cell types were enriched in accessible chromatin of the corresponding cell types. The p-values were adjusted for multiple hypothesis testing using the Bonferroni method (Table S5).

### Luciferase assays

We tested three variants for enhancer activity using luciferase assays. We previously published the luciferase assay corresponding to imVar rs72928038^92^. The assays for imVar rs6908626 and non-impact variant rs17574546 are new to this study (Table S5). We cloned short sequences containing the reference alleles for rs72928038, rs6908626, and rs17574546 and alternate alleles for rs6908626, rs17574546 from human DNA samples (Coriell) into the luciferase reporter vector pGL4.23 (Promega) in the forward direction with restriction enzymes SacI-HF (NEB) and KpnI-HF (NEB). The alternate allele for rs72928038 was introduced via site-directed mutagenesis using the Q5 SDM kit (NEB). Plasmids were prepared from competent DH5alpha *E.coli* (NEB) using the Qiaspin Miniprep Kit (Qiagen) and sequences were verified via Sanger sequencing. Sequences used for cloning and Sanger sequencing are listed in Table S5.

Jurkat cells from ATCC were maintained in culture according to manufacturer’s recommendations at a concentration of 1×10^5^/mL-1×10^6^/mL and tested every 10 passages to confirm lack of mycoplasma contamination. Approximately 0.5×10^6^ cells per replicate (3 replicates) were co-transfected with 500 ng of firefly luciferase vector containing either the reference or alternate allele or an empty pGL4.23 vector as a control, and 50 ng pRL-SV40 Renilla luciferase vector (Promega), using the Lipofectamine LTX reagent. Media was refreshed 24 hours after transfection with standard culture media containing 30U/mL IL-2 and Human T-Activator CD3/CD28 Dynabeads (Invitrogen) at a 1:1 ratio with cell counts where indicated. Cells were collected 48 hours post transfection and assayed using the Dual-Luciferase Reporter system (Promega). Firefly activity was normalized to the Renilla activity and expressed as fold change compared to the luciferase activity of the empty vector (RLU). A two-sided t-test was used to compare the luciferase activity between the two alleles in the stimulated and unstimulated treatment conditions, and separately to compare normalized enhancer activity for each allele in the stimulated and unstimulated conditions. Benjamini-Hochberg FDR was used to correct for MHT.

### imVar comparison with MPRAs

MPRA summary statistics for T1D variants from Jurkat T cells^39^ and primary T cells^40^, were downloaded from the supplementary table from each respective publication. Based on these summary statistics, we determined emVars using the same criteria as the authors of each respective study, which yielded the same number of previously reported emVars for Jurkat cells (N = 313) but an additional 2 emVars for the T cells (N = 547). The variant locations from the Jurkat study were lifted from hg19 to hg38 to match the T1D DVIM variants analysed. We overlapped variants tested by DVIM and each respective MPRA by position to define variants tested by both methods (N = 1,359 for both studies). We then constructed contingency tables of overlap between imVars, non-imVars, emVars, and non-emVars, with emVars defined by either study (Table S7). We then calculated odds ratios, confidence intervals, and performed Fisher’s exact test using Scipy^93^, and corrected for multiple hypothesis testing with Benjamini-Hochberg FDR.

### imVar comparison RAVAR

RAVAR^44^ variant-trait associations were downloaded from http://www.ravar.bio/api/download/static/snp_fulltable.txt. RAVAR variants were intersected with all DVIM tested rare variants for each respective trait GWAS by the position of the genetic variants, since not all of the rare variants had associated reference SNP cluster identifiers (RSIDs). For traits where at least 20 RAVAR variants had been tested with DVIM, we constructed contingency tables overlapping the trait rare imVars and rare non-imVars with the RAVAR variants, taking all tested rare variants not within the RAVAR database as non-RAVAR variants (Table S5,S7). We then calculated odds ratios, confidence intervals, and performed Fisher’s exact test using Scipy^93^, and corrected for multiple hypothesis testing with Benjamini-Hochberg FDR.

### Data and code availability

All data utilised are publicly available. ENCODE^27^ data tracks used are publicly available at https://www.encodeproject.org/. GTEx^23^ v8 data utilised is available at https://gtexportal.org/home/. Independent validation ATAC-seq data^38^ is available via the Gene Expression Omnibus at accession GSE163160. DNACipher and DVIM are available with a Python interface and command-line tool at https://github.com/BradBalderson/DNACipher.

## Contributions

BB conceived the project, secured funding, designed the approach and analysis, developed software, performed benchmarking, made figures, and wrote the manuscript. ST consulted on DNACipher training strategy, implemented initial code for Enformer embedding generation, benchmarked DeepCT for the eQTL analysis, edited the manuscript, and tested software. MO and SC designed, performed, and analyzed the luciferase assay experiments in Jurkat cells. WJFR benchmarked Enformer for the causal eQTL analysis, compiled ADASTRA data, edited the manuscript, and tested software. JJ edited the manuscript and tested software. NP edited the manuscript and secured funding. KJG secured funding and supervised the luciferase experiments. LY secured funding, supervised the multi-trait DVIM analysis, ideated many Discussion points, and edited the manuscript. MB conceived the project, supervised the project, secured funding, and edited the manuscript. GM conceived the project, supervised the project, secured funding, and edited the manuscript.

## Acknowledgements

BB was supported by a Hewitt Foundation Fellowship and Ekhart Scholarship, as well as a Medical Research Future Fund (MRFF) grant to NP and MB (grant number 2016033), and a Snow Medical Fellowship to LY. ST was also supported by the same MRFF grant. MO was supported by NIGMS/NIH T32 award GM008666. WJFR was supported by an Australian Research Training Program (RTP) scholarship. JJ was supported by NIH F31 fellowship HG013262. KJG was supported by NIH grants HG012059 and DK138512, and a grant from the Larry L Hillblom Foundation. GM was supported by NIH grant HG011315 and a grant from the Larry L Hillblom Foundation. We thank Qiongyi Zhao, Weston Elison, Chris Duroiu, Han Chen, and David Laub for testing the DNACipher software and providing feedback. We also thank Johannes C. Hingerl for providing performance summary statistics for Borzoi.

## Extended Data Figures

**Extended Data Figure 1.**
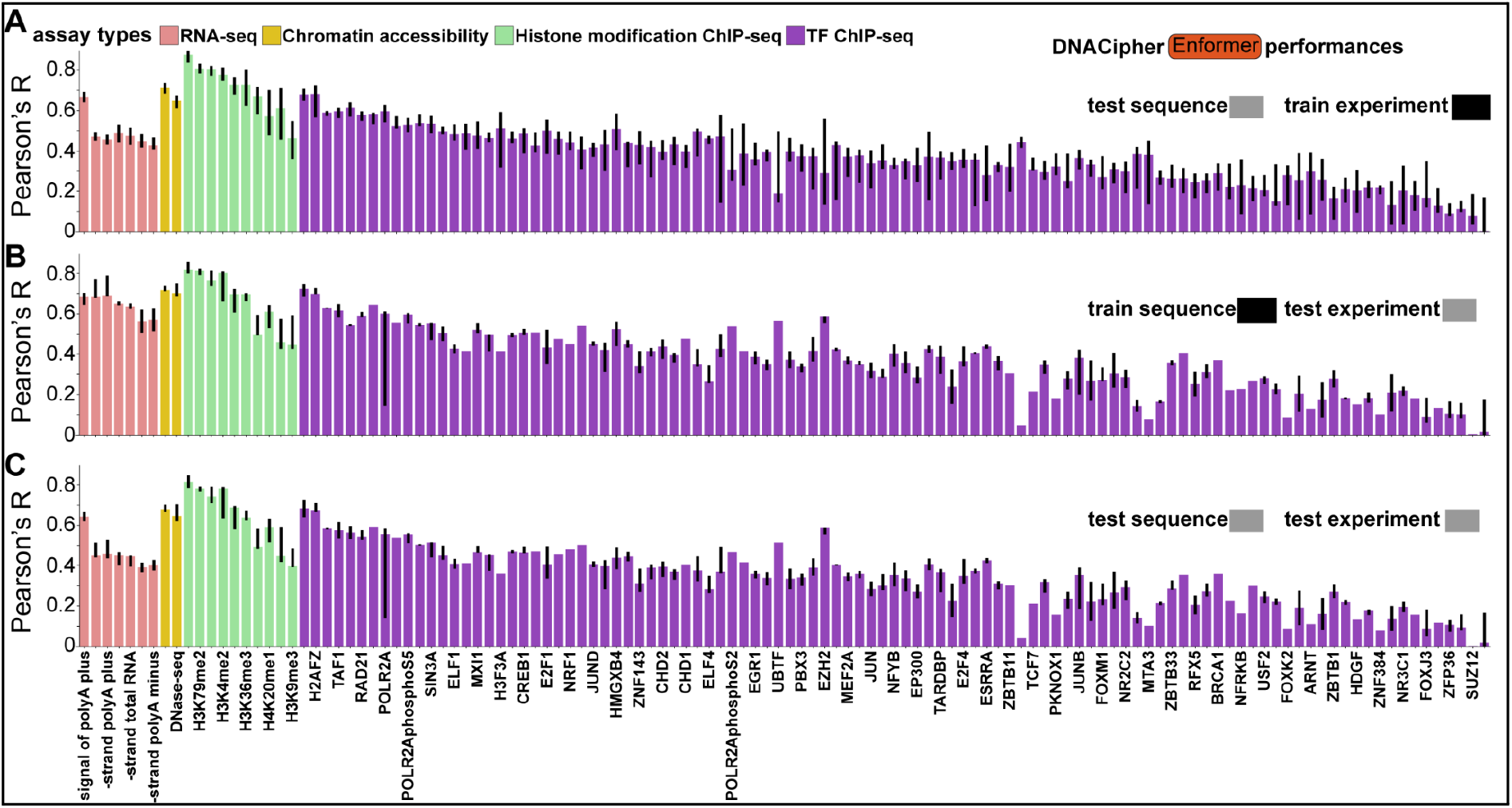
DNACipher-Enformer performance metrics by assay type. A-C. Barcharts of median Pearson’s R for DNACipher-Enformer predictions against actual experimental signals stratified by assay, for (**A**) test sequences in train experiments, (**B**) train sequences in test experiments, (**C**) test sequences in test experiments. Correlations are compiled across 5 cross-validations of train/test cell/tissue types for each assay. Error bars indicate 0.25-0.75 quantiles. Error bars are missing where only one experiment could be utilized for the relevant stratification. Sample sizes are listed in Table S1.

**Extended Data Figure 2.**
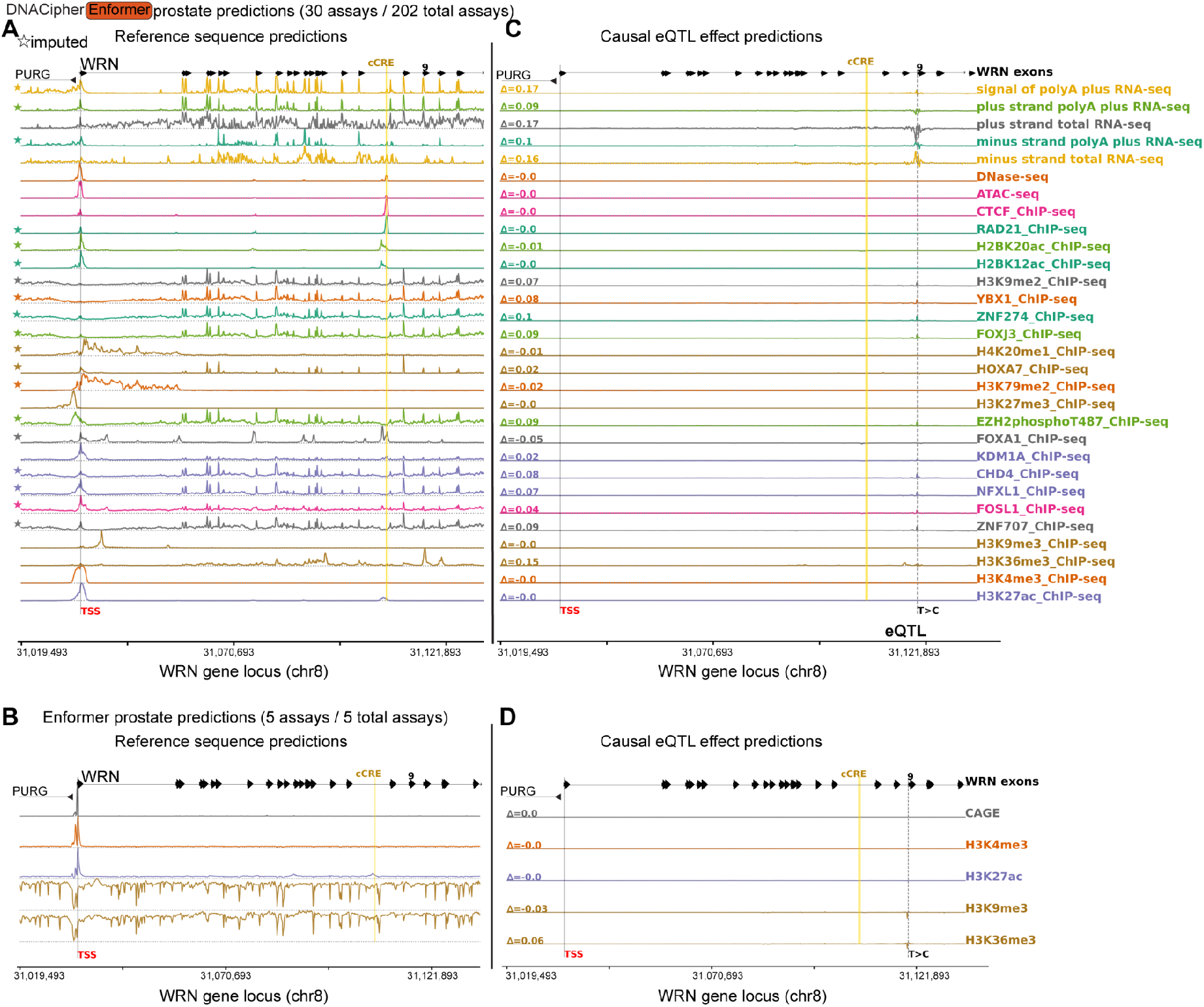
DNACipher-Enformer predicts an eQTL effect that is not detected with Enformer. **A.** DNACipher-Enformer signal predictions for the top 30 prostate-specific assays (of 202 total) at the *WRN* locus reference genome sequence (test sequence). The x-axis is the genome position. Assays marked with stars are imputed. The *WRN* transcription start site (TSS), *WRN* exons, an ENCODE candidate Cis-Regulatory Element (cCRE), and opposite strand gene *PURG* are annotated. **B.** Equivalent to A, except with Enformer prostate predictions, which are limited to 5 assays. **C.** Predicted change in assay signals between the reference and alternate alleles of a likely causal expression quantitative trait locus (eQTL, rs4733225) variant for *WRN*. Δ values are the predicted fold-change in signal (alternative-reference sequence predictions) across the WRN locus for the likely causal eQTL. **D.** Equivalent to C, except with Enformer.

**Extended Data Figure 3.**
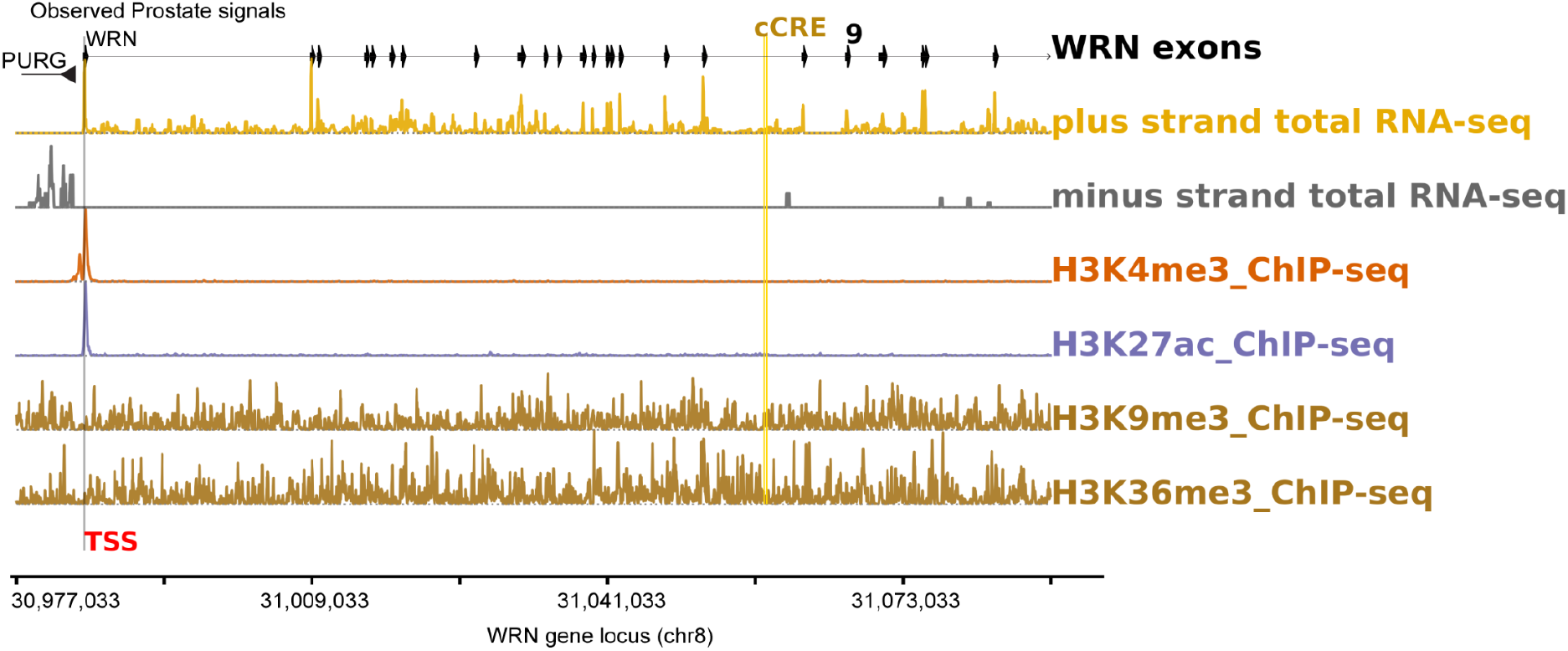
ENCODE genomic signals at the *WRN* gene locus. Signal tracks from ENCODE prostate experiments at the *WRN* locus reference genome sequence (test sequence). The x-axis is the genome position. The *WRN* transcription start site (TSS), *WRN* exons, an ENCODE candidate Cis-Regulatory Element (cCRE), and opposite strand gene *PURG* are annotated.

**Extended Data Figure 4.**
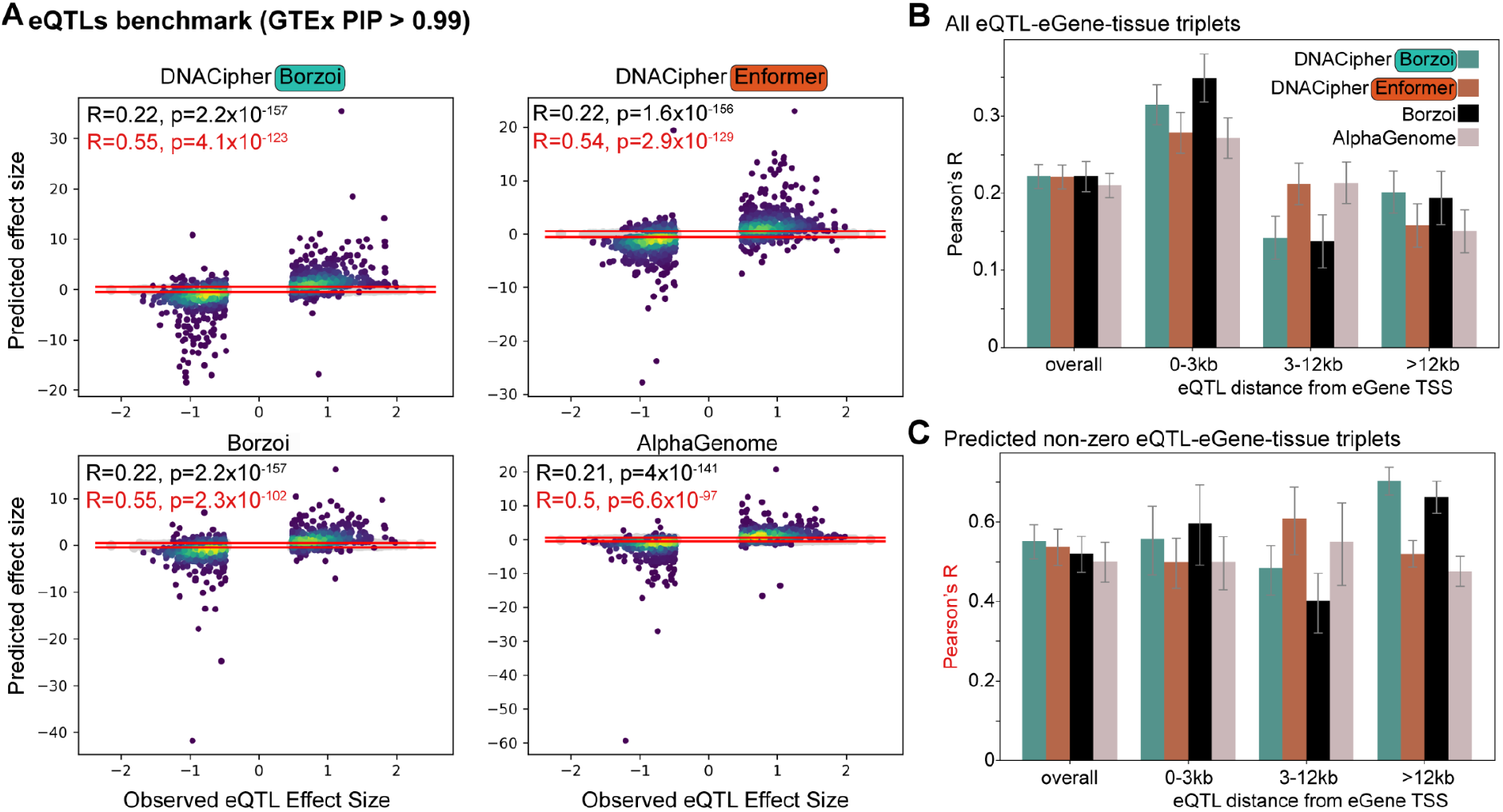
DNACipher predicts GTEx eQTL effects with similar performance to AlphaGenome. **A.** Scatter plots where each point corresponds to a likely causal eQTL-eGene-tissue combination (N = 14,196 in each plot). The x-axis is the measured eQTL effect size (β) and the y-axis is the predicted eQTL effect sizes (z-scaled) from DNACipher-Borzoi, DNACipher-Enformer, Borzoi, or AlphaGenome. Points are coloured by density estimated using a Gaussian kernel density estimator^93^. Pearson’s correlations (R) and beta-distribution p-values indicate correlation between predicted and actual eQTL effect sizes. R is also calculated for non-zero predicted effect sizes above the horizontal red lines (|z|>0.5). **B.** Bar plots with Pearson’s R between predicted and actual eQTL-eGene-tissue effect sizes with error bars indicating 95% confidence intervals. Performances are further stratified by all eQTL-eGene-tissue effects (‘overall’), eQTLs within 0-3 kb of the eGene TSS (N = 4,732), 3-12 kb from eGene TSS (N = 4,770), and >12 kb (N = 4,694). **C.** Equivalent to B, except for non-zero predicted effect size correlations.

**Extended Data Figure 5.**
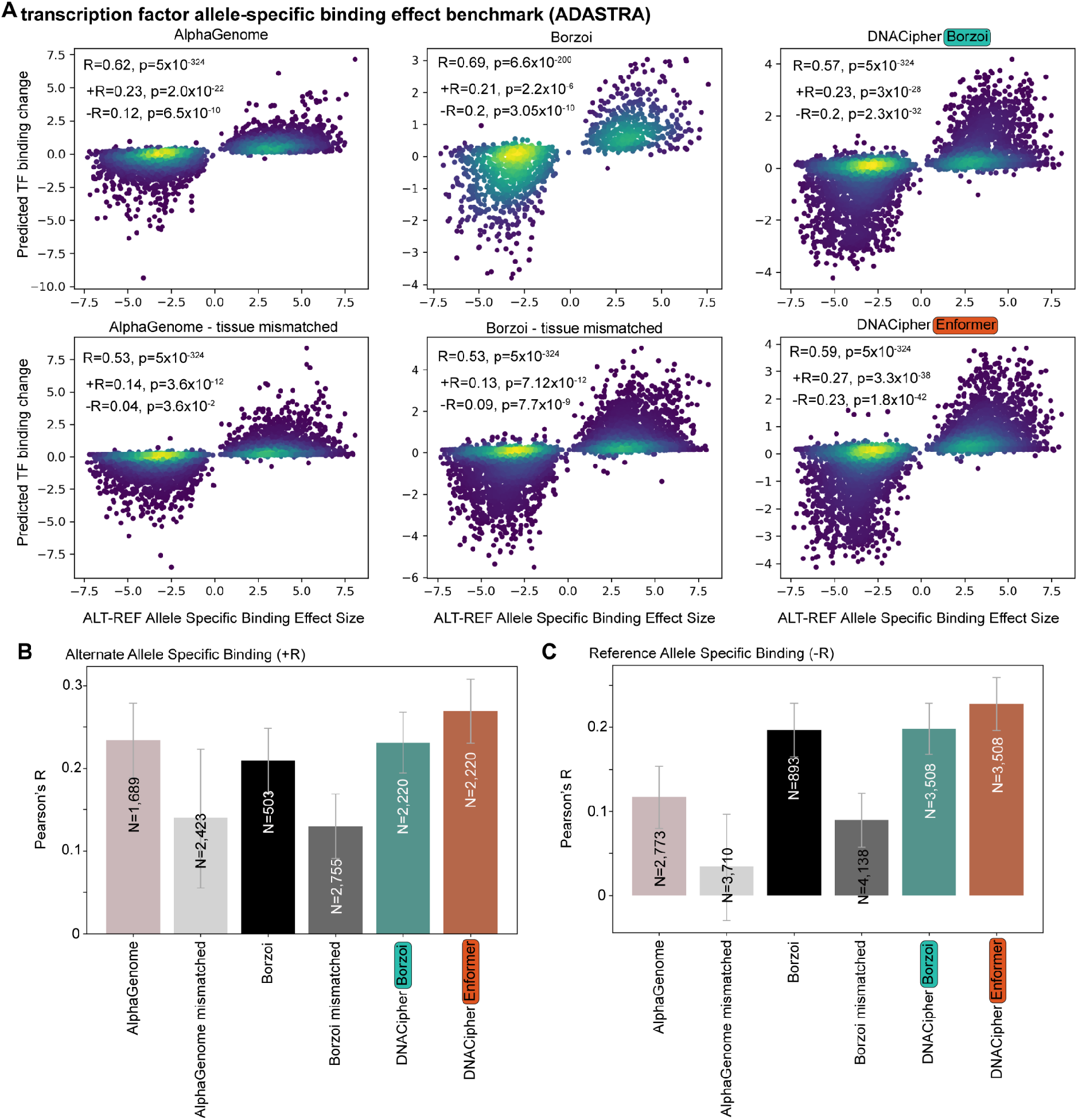
DNACipher predicts transcription factor (TF) allele-specific binding (ASB) with greater performance than AlphaGenome for missing tissues. **A.** Scatter plots where each point corresponds to a TF ASB event in a particular cell/tissue from the ADASTRA database (N varies for each plot due to differences in events that can be predicted by each model, and are listed in panel B). The x-axis is the measured allele-specific effect size (>0 indicates preference for alternative allele) and the y-axis is the predicted effect sizes (z-scaled) from AlphaGenome, Borzoi, DNACipher-Borzoi, or DNACipher-Enformer. For TF ASB events that could not be predicted for AlphaGenome/Borzoi, predictions are shown for a random tissue where predictions could be generated for the corresponding TF. Points are coloured by density estimated using a Gaussian kernel density estimator^93^. Overall Pearson’s correlation (R) are indicated with beta-distribution p-values. Additionally, correlations are shown for alternative allele binding preference (+R, TF ASB effect size > 0) or reference allele binding preference (-R, TF ASB effect size < 0). **B.** Bar plots with Pearson’s R values between model predicted and actual TF ASB effect sizes with preference for the reference allele (-R). Error bars indicate 95% confidence intervals. **C.** Equivalent to B, except for TF ASB events with alternative allele binding preference (+R).

**Extended Data Figure 6.**
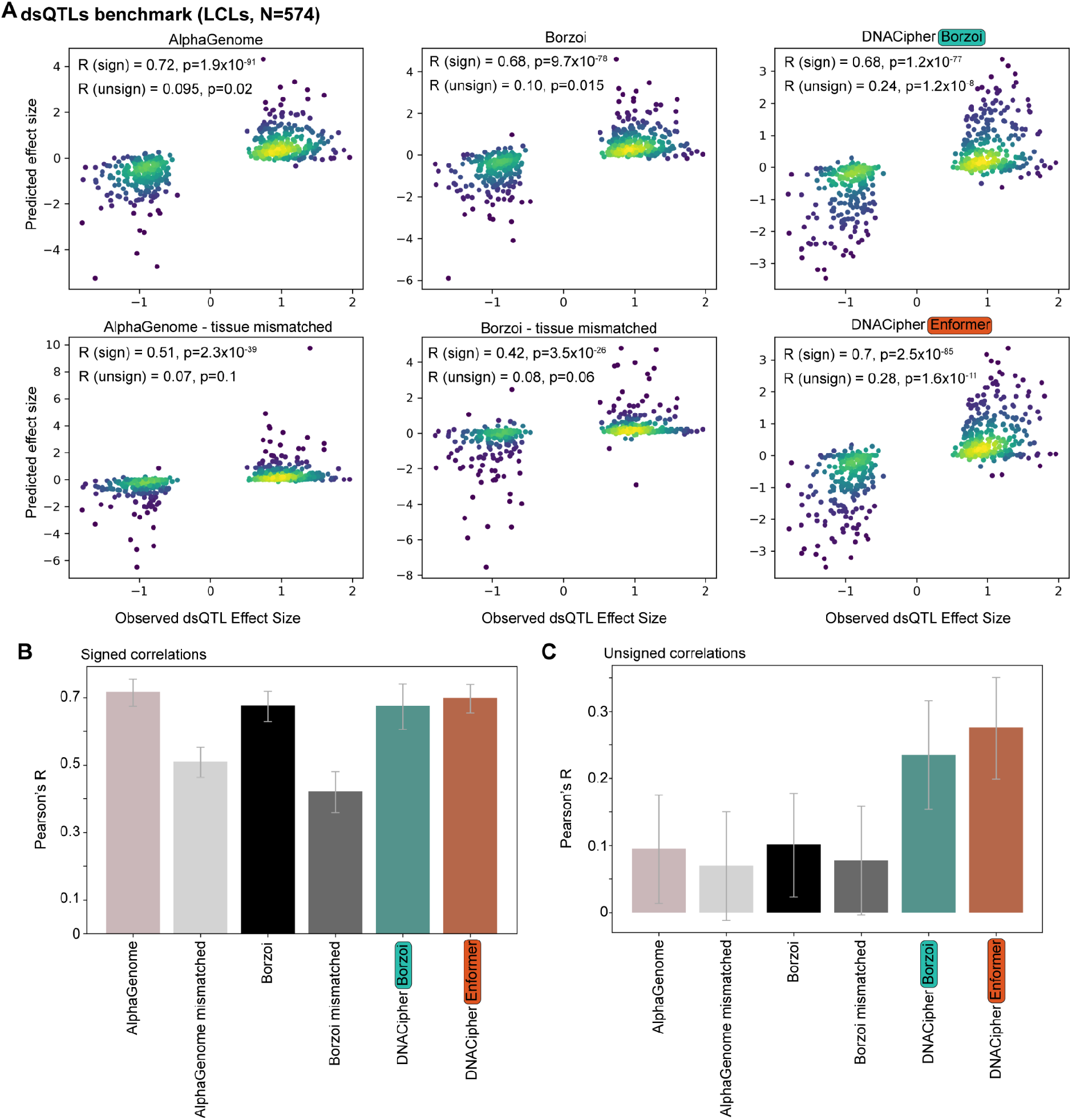
DNACipher predicts dsQTL effect sizes with similar performance to AlphaGenome. **A.** Scatter plots where each point corresponds to a dsQTL-dsPeak effect size in Yoruba lymphoblastoid cell lines (LCLs) (N = 574). The x-axis is the measured dsQTL effect size (>0 indicates preference for alternative allele) and the y-axis is the predicted effect sizes (z-scaled) from AlphaGenome, Borzoi, DNACipher-Borzoi, or DNACipher-Enformer. To simulate performance if mis-matched tissues were used because of missing experiments with AlphaGenome/Borzoi, predictions are also shown for using random cell type/tissues. Points are coloured by density estimated using a Gaussian kernel density estimator^93^. Overall Pearson’s correlation (‘R (sign)’) are indicated with beta-distribution p-values. Correlations are also shown for the absolute effect size prediction (‘R (unsign)’). **B.** Bar plots with the overall Pearson’s R (sign) values between model predicted and actual dsQTL effect sizes. Error bars indicate 95% confidence intervals. **C.** Equivalent to B, except for the correlations for magnitude (‘R (unsigned)’).

**Extended Data Figure 7.**
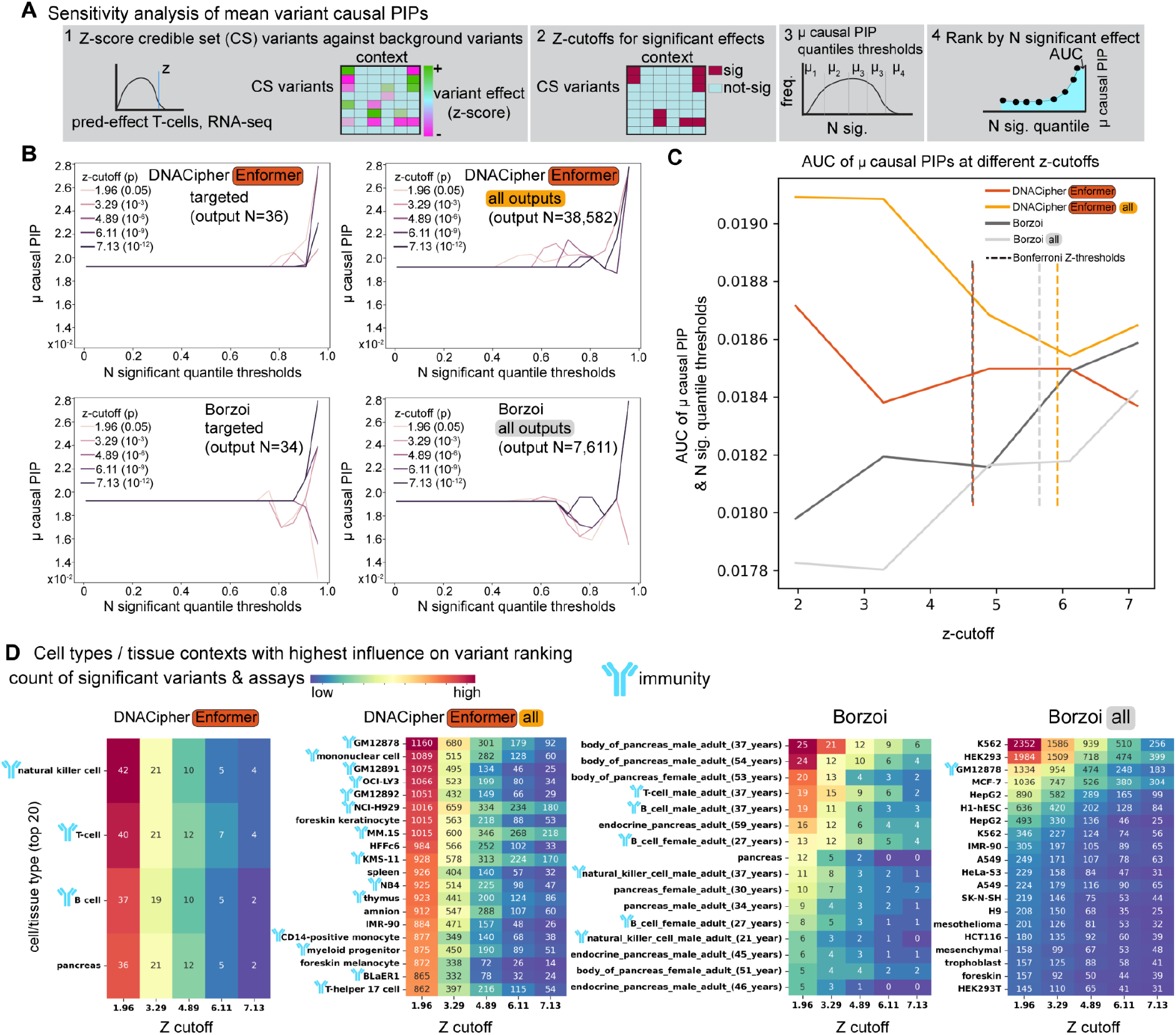
DNACipher comprehensive molecular effect prediction outperforms DVIM targeted analysis and overcomes ascertainment bias of predictable contexts. **A.** Overview of sensitivity analysis; 1) z-scoring of credible set (CS) variants against background (non-trait associated) variant effects for each predicted context, 2) z-cutoffs to threshold significant/non-significant effects, 3) ranking of CS variants by the number of significant effects, and taking the mean posterior inclusion probability (PIP) of variants with N significant effects above different quantile thresholds of N significant effects, 4) Area under the curve (AUC) of mean PIPs (y-axis) against N significant effect quantile thresholds to measure ranking of causal variants. **B.** Line plots of the N significant quantile thresholds (x-axis) and the mean PIP for variants above the quantile threshold (y-axis) at different z-cutoffs (colored) for DNACipher-Enformer targeted analysis (N contexts outputs = 36), DNACipher-Enformer all outputs (N context outputs = 38,582), Borzoi targeted (N context outputs =34) and Borzoi all outputs (N context outputs = 7,611). This analysis was conducted for the T1D *RUNX3* locus (N = 43 CS variants). **C.** AUC of N significant effects quantile threshold and mean PIP curves (y-axis) at different z-cutoffs (x-axis) for the different models and contexts. Vertical dashed lines indicate the Bonferroni z-threshold for multiple hypothesis testing correction for the number of contexts tested for each relevant model/context output (Bonferroni p-value threshold of < 0.05). **D.** Heatmaps displaying the count of significant assays across variants for each cell/tissue type (rows; maximum top 20) at different significant thresholds (z-cutoffs, columns) for each model/context output. Higher values indicate more variants were ranked highly due to the corresponding cell/tissue type at the z-cutoff. Immunity-related cell/tissue types are indicated with an antibody schematic, showing DNACipher-Enformer with all outputs ranks immunity-related tissues highly (of 191 cell/tissue types) while using all Borzoi outputs ranked cancer cell lines highly (K562, HEK293, HepG2) reflecting ascertainment bias of training experiments.

**Extended Data Figure 8.**
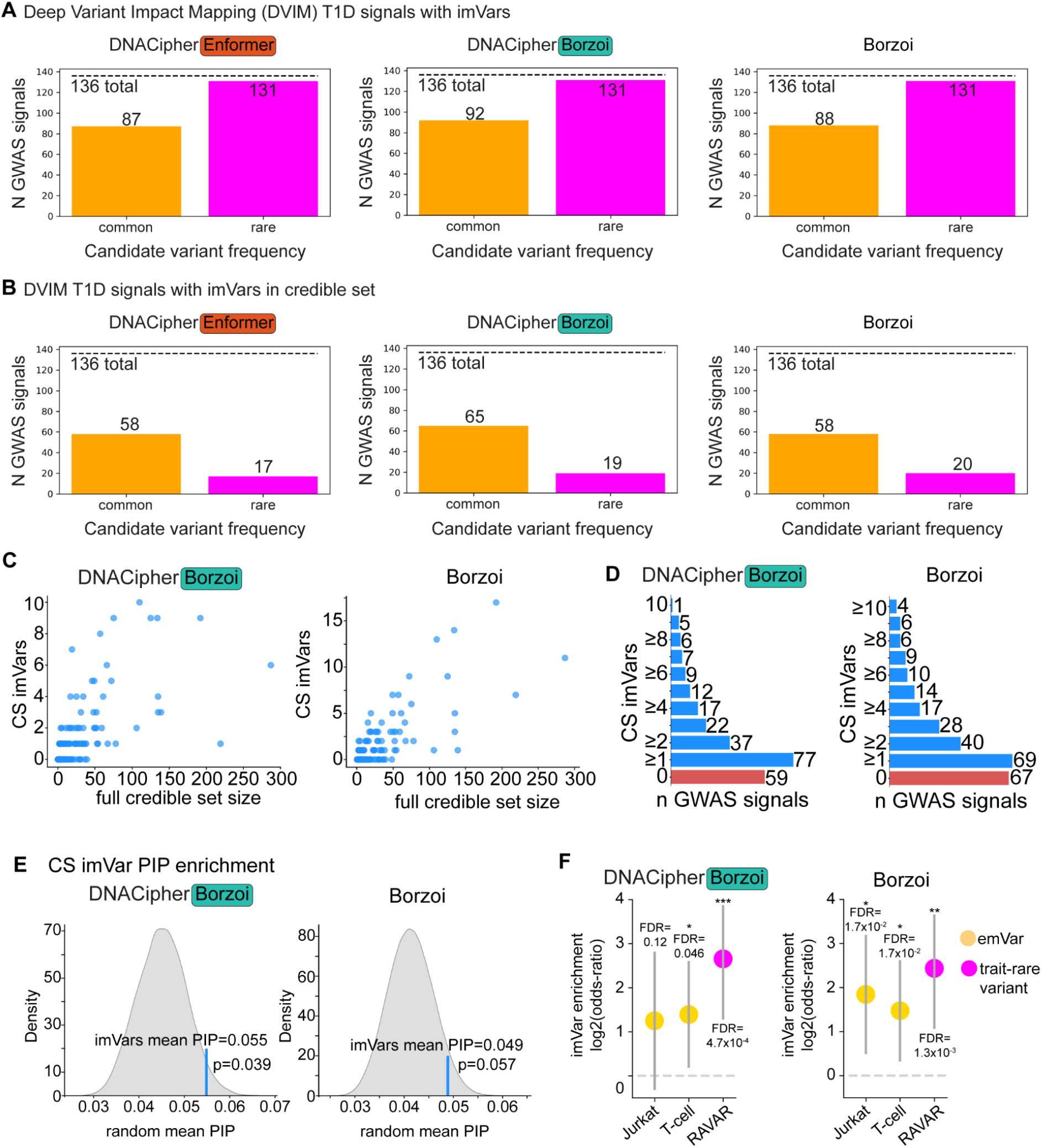
Comparison of T1D Deep Variant Impact Mapping (DVIM) of T1D between models. **A.** Barplot showing the number of T1D GWAS signals that had at least one common or rare imVar by DVIM using different variant effect predictions from either DNACipher-Enformer, DNACipher-Borzoi, or Borzoi. **B.** Equivalent to A, except only considering fine-mapping credible set (CS) variants. **C.** Scatter plots of CS sizes before (x-axis) and after imVar filtering (y-axis), using imVars from DVIM using variant effect predictions from DNACipher-Borzoi or Borzoi. **D.** Ranked bar chart of N GWAS loci (x-axis) with N imVars in CS (y-axis) from DVIM with either DNACipher-Borzoi or Borzoi. **E.** Distribution of mean PIPs from random CS sampling (grey; N = 1 x 10^6^) with observed mean PIP of CS imVars as a vertical blue line with empirical p-value for either DNACipher-Borzoi or Borzoi. **F.** Log2-odds-ratio (y-axis) of overlap between imVars and expression modifying variants (emVars) from either Jurkat or T-cells, and rare imVars with rare trait-associated variants from the Rare Variant Association Repository (RAVAR) using imVars from DNACipher-Borzoi or Borzoi variant effect predictions. The dotted line indicates the random expectation, and the error bars indicate 95% confidence intervals. Fisher’s exact test with FDR correction was used for statistical comparisons. Sample sizes are provided in Table S5.

**Extended Data Figure 9.**
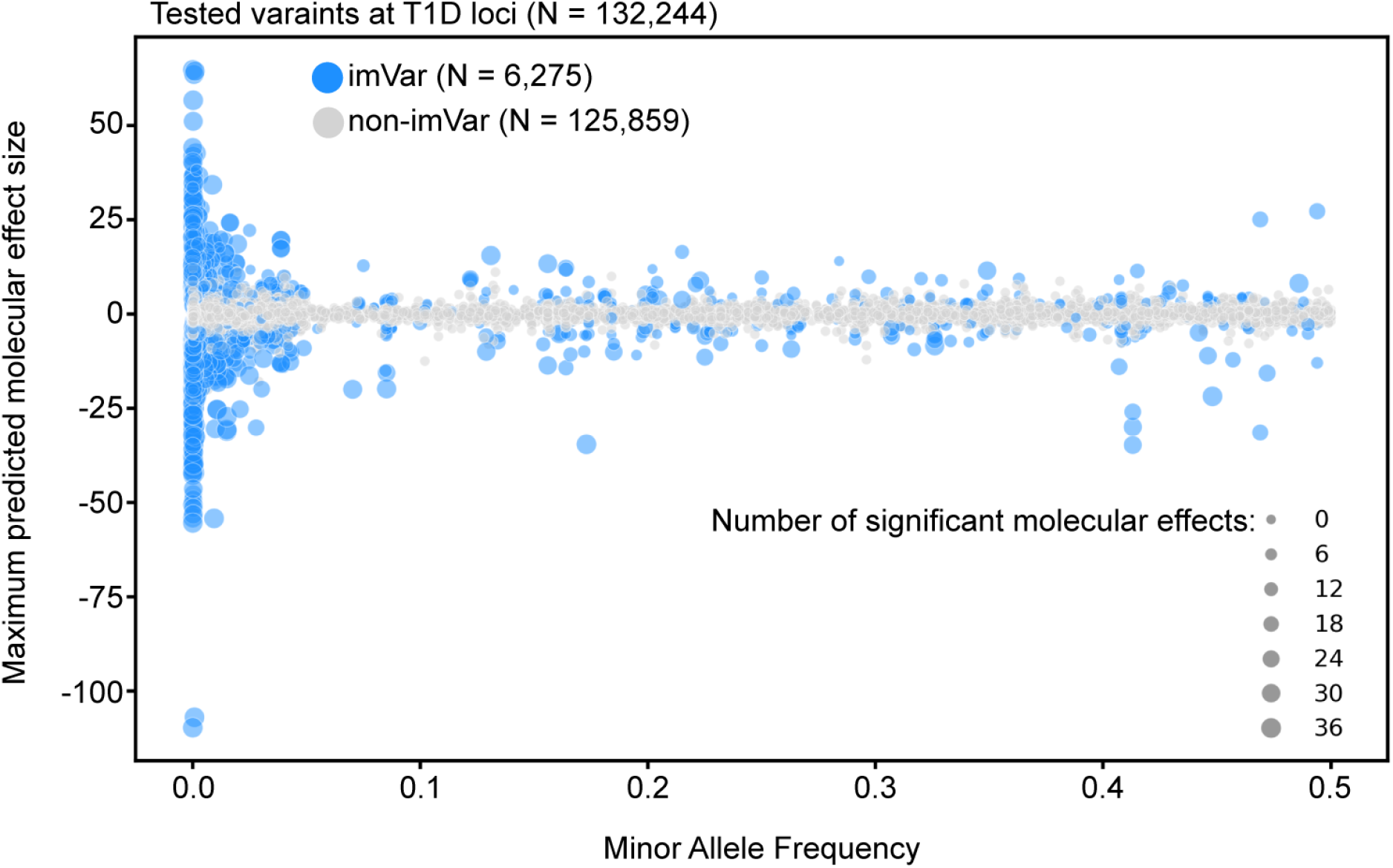
DNACipher-Enformer predicted effect sizes and minor allele frequency. Scatter plot of the maximum predicted effect size from DNACipher-Enformer across all targeted T1D contexts (N = 36, y-axis) for Deep Variant Impact Mapping (DVIM) tested common and rare variants at T1D loci and the minor allele frequency (x-axis). Point sizes correspond to the number of significant effects compared to non-trait associated variants at the same genome regions (boot-strapped one-sample two-sided z-test, N boot-straps = 1,000,000).

**Extended Data Figure 10.**
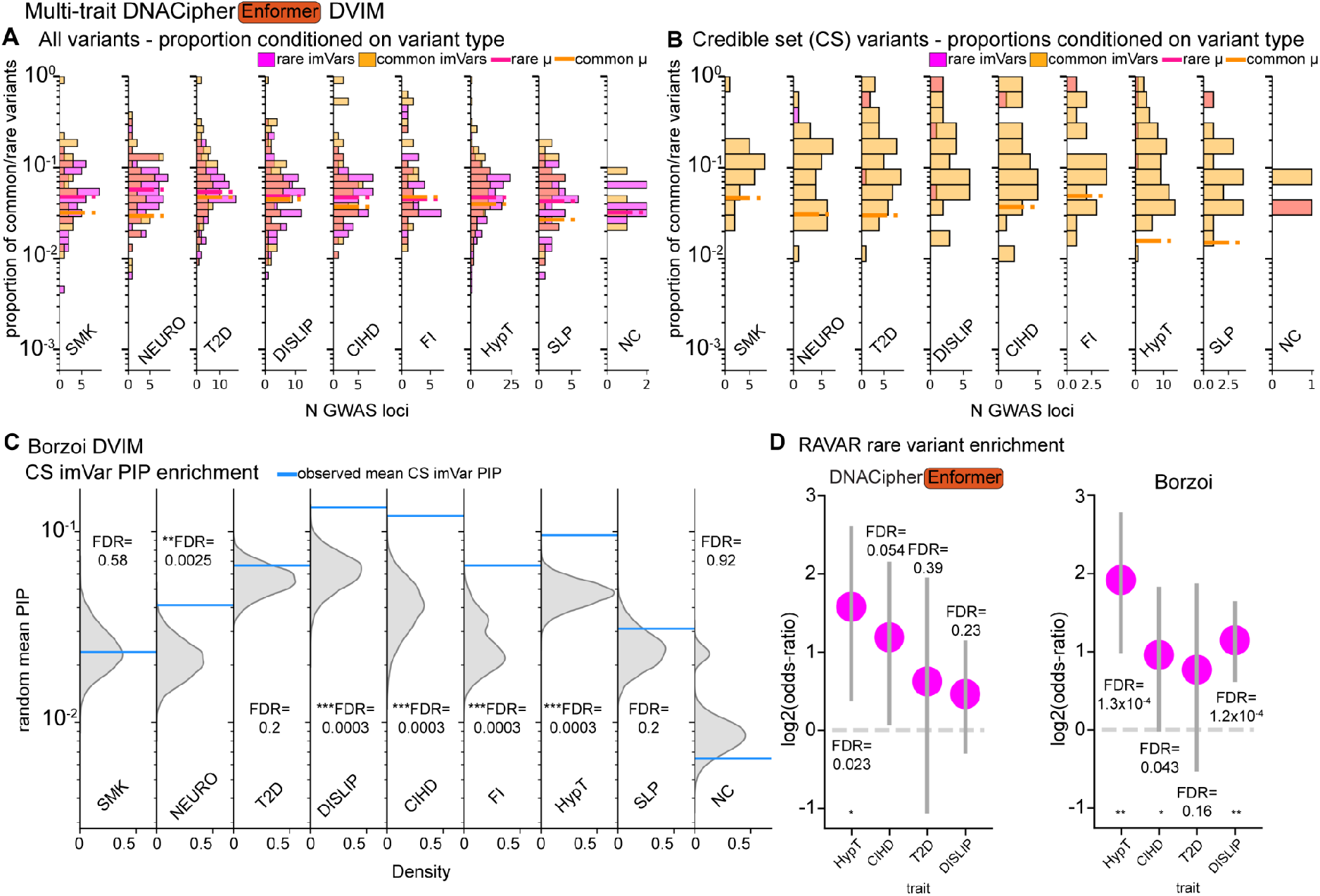
Additional data of Deep Variant Impact Mapping (DVIM) across 9 complex traits. **A.** Histograms for each trait, showing the distribution across loci of the proportion of rare variants that were rare imVars, and the proportion of common variants that were common imVars. Dashed lines indicate the mean imVar proportion of each type across loci. Figure 6D shows the equivalent for the proportion of all variants analysed at each locus. **B.** Equivalent to A, except subsetted to CS variants. Figure 6E shows the equivalent for the proportion of all CS variants analysed at each locus. **C.** Distributions of mean PIPs for random CS variants (N = 10,000) for each trait, with blue lines indicating the mean PIP for imVars called using Borzoi variant effect predictions. Benjamini-Hochberg false discovery rate (FDR) corrected empirical p-values indicate the significance of the mean imVar PIP. **D.** Log2-odds-ratio (y-axis) of overlap between rare imVars with rare trait-associated variants from the Rare Variant Association Repository (RAVAR) using imVars from DNACipher-Enformer or Borzoi variant effect predictions. The dotted line indicates the random expectation, and the error bars indicate 95% confidence intervals. Fisher’s exact test with FDR correction was used for statistical comparisons. Sample sizes are provided in Table S7.

## References

1. McCarthy, M. I. et al. Genome-wide association studies for complex traits: consensus, uncertainty and challenges. Nat. Rev. Genet. 9, 356–369 (2008).

2. Zou, Y., Carbonetto, P., Wang, G. & Stephens, M. Fine-mapping from summary data with the ‘Sum of Single Effects’ model. PLoS Genet. 18, e1010299 (2022).

3. Dong, S. et al. Annotating and prioritizing human non-coding variants with RegulomeDB v.2. Nat. Genet. 55, 724–726 (2023).

4. Dong, S. & Boyle, A. P. Predicting functional variants in enhancer and promoter elements using RegulomeDB: DONG and BOYLE. Hum. Mutat. 40, 1292–1298 (2019).

5. Ritchie, G. R. S., Dunham, I., Zeggini, E. & Flicek, P. Functional annotation of noncoding sequence variants. Nat. Methods 11, 294–296 (2014).

6. Rentzsch, P., Witten, D., Cooper, G. M., Shendure, J. & Kircher, M. CADD: predicting the deleteriousness of variants throughout the human genome. Nucleic Acids Res. 47, D886–D894 (2019).

7. Rentzsch, P., Schubach, M., Shendure, J. & Kircher, M. CADD-Splice-improving genome-wide variant effect prediction using deep learning-derived splice scores. Genome Med. 13, 31 (2021).

8. Gao, H. et al. The landscape of tolerated genetic variation in humans and primates. Science 380, eabn8153 (2023).

9. Frazer, J. et al. Disease variant prediction with deep generative models of evolutionary data. Nature 599, 91–95 (2021).

10. McQuerry, J. A. et al. Massively parallel identification of functionally consequential noncoding genetic variants in undiagnosed rare disease patients. Sci. Rep. 12, 7576 (2022).

11. Liu, L. et al. Biological relevance of computationally predicted pathogenicity of noncoding variants. Nat. Commun. 10, 330 (2019).

12. Zhou, J. & Troyanskaya, O. G. Predicting effects of noncoding variants with deep learning-based sequence model. Nat. Methods 12, 931–934 (2015).

13. Kelley, D. R., Snoek, J. & Rinn, J. L. Basset: learning the regulatory code of the accessible genome with deep convolutional neural networks. Genome Res. 26, 990–999 (2016).

14. Zhou, J. et al. Deep learning sequence-based ab initio prediction of variant effects on expression and disease risk. Nat. Genet. 50, 1171–1179 (2018).

15. Kelley, D. R. et al. Sequential regulatory activity prediction across chromosomes with convolutional neural networks. Genome Res. 28, 739–750 (2018).

16. Kelley, D. R. Cross-species regulatory sequence activity prediction. PLoS Comput. Biol. 16, e1008050 (2020).

17. Avsec, Ž., et al. Effective gene expression prediction from sequence by integrating long-range interactions. Nat. Methods 18, 1196–1203 (2021).

18. Chen, K. M., Wong, A. K., Troyanskaya, O. G. & Zhou, J. A sequence-based global map of regulatory activity for deciphering human genetics. Nat. Genet. 54, 940–949 (2022).

19. Linder, J., Srivastava, D., Yuan, H., Agarwal, V. & Kelley, D. R. Predicting RNA-seq coverage from DNA sequence as a unifying model of gene regulation. Nat. Genet. (2025) doi:10.1038/s41588-024-02053-6.

20. Wesolowska-Andersen, A. et al. Deep learning models predict regulatory variants in pancreatic islets and refine type 2 diabetes association signals. Elife 9, (2020).

21. Lee, P. H. et al. Principles and methods of in-silico prioritization of non-coding regulatory variants. Hum. Genet. 137, 15–30 (2018).

22. Smemo, S. et al. Obesity-associated variants within FTO form long-range functional connections with IRX3. Nature 507, 371–375 (2014).

23. GTEx Consortium. The GTEx Consortium atlas of genetic regulatory effects across human tissues. Science 369, 1318–1330 (2020).

24. Avsec, Ž., et al. Advancing regulatory variant effect prediction with AlphaGenome. Nature 649, 1206–1218 (2026).

25. Schreiber, J., Durham, T., Bilmes, J. & Noble, W. S. Avocado: a multi-scale deep tensor factorization method learns a latent representation of the human epigenome. Genome Biol. 21, 81 (2020).

26. Boix, C. A., James, B. T., Park, Y. P., Meuleman, W. & Kellis, M. Regulatory genomic circuitry of human disease loci by integrative epigenomics. Nature 590, 300–307 (2021).

27. ENCODE Project Consortium et al. Expanded encyclopaedias of DNA elements in the human and mouse genomes. Nature 583, 699–710 (2020).

28. Ernst, J. & Kellis, M. Large-scale imputation of epigenomic datasets for systematic annotation of diverse human tissues. Nat. Biotechnol. 33, 364–376 (2015).

29. Gao, Z., Liu, Q., Zeng, W., Jiang, R. & Wong, W. H. EpiGePT: a pretrained transformer-based language model for context-specific human epigenomics. Genome Biol. 25, 310 (2024).

30. Murphy, A. E., Beardall, W., Rei, M., Phuycharoen, M. & Skene, N. G. Predicting cell type-specific epigenomic profiles accounting for distal genetic effects. Nat. Commun. 15, 9951 (2024).

31. Sindeeva, M. et al. Cell type-specific interpretation of noncoding variants using deep learning-based methods. Gigascience 12, (2023).

32. Mostafavi, H., Spence, J. P., Naqvi, S. & Pritchard, J. K. Systematic differences in discovery of genetic effects on gene expression and complex traits. Nat. Genet. 55, 1866–1875 (2023).

33. Narita, T. et al. Acetylation of histone H2B marks active enhancers and predicts CBP/p300 target genes. Nat. Genet. 55, 679–692 (2023).

34. Hingerl, J. C., et al. scooby: Modeling multi-modal genomic profiles from DNA sequence at single-cell resolution. bioRxivorg (2024) doi:10.1101/2024.09.19.613754.

35. Abramov, S. et al. Landscape of allele-specific transcription factor binding in the human genome. Nat Commun 12, 2751 (2021).

36. Pampari, A. et al. ChromBPNet: bias factorized, base-resolution deep learning models of chromatin accessibility reveal cis-regulatory sequence syntax, transcription factor footprints and regulatory variants. bioRxivorg (2025) doi:10.1101/2024.12.25.630221.

37. Schreiber, J. tangermeme: A toolkit for understanding cis-regulatory logic using deep learning models. bioRxiv (2025) doi:10.1101/2025.08.08.669296.

38. Chiou, J. et al. Interpreting type 1 diabetes risk with genetics and single-cell epigenomics. Nature 594, 398–402 (2021).

39. Ho, C.-H. et al. Genetic and epigenetic screens in primary human T cells link candidate causal autoimmune variants to T cell networks. Nat Genet 57, 2536–2545 (2025).

40. Mouri, K. et al. Prioritization of autoimmune disease-associated genetic variants that perturb regulatory element activity in T cells. Nat. Genet. 54, 603–612 (2022).

41. McCarthy, M. I. & Hirschhorn, J. N. Genome-wide association studies: potential next steps on a genetic journey. Hum Mol Genet 17, R156–65 (2008).

42. Quintana-Murci, L. Understanding rare and common diseases in the context of human evolution. Genome Biol. 17, 225 (2016).

43. Fiziev, P. P. et al. Rare penetrant mutations confer severe risk of common diseases. Science 380, eabo1131 (2023).

44. Cao, C. et al. RAVAR: a curated repository for rare variant-trait associations. Nucleic Acids Res. 52, D990–D997 (2024).

45. Barton, A. R., Sherman, M. A., Mukamel, R. E. & Loh, P.-R. Whole-exome imputation within UK Biobank powers rare coding variant association and fine-mapping analyses. Nat. Genet. 53, 1260–1269 (2021).

46. Vuckovic, D. et al. The polygenic and monogenic basis of blood traits and diseases. Cell 182, 1214–1231.e11 (2020).

47. Astle, W. J. et al. The Allelic landscape of human blood cell trait variation and links to common complex disease. Cell 167, 1415–1429.e19 (2016).

48. Pietzner, M. et al. Mapping the proteo-genomic convergence of human diseases. Science 374, eabj1541 (2021).

49. Wainschtein, P. et al. Estimation and mapping of the missing heritability of human phenotypes. Nature 649, 1219–1227 (2026).

50. Wainschtein, P. et al. Assessing the contribution of rare variants to complex trait heritability from whole-genome sequence data. Nat Genet 54, 263–273 (2022).

51. Taliun, D. et al. Sequencing of 53,831 diverse genomes from the NHLBI TOPMed Program. Nature 590, 290–299 (2021).

52. Souaiaia, T. et al. Distinct genetic architecture in the tails of complex traits. Nature 655, 676–684 (2026).

53. Klein, J. C. et al. A systematic evaluation of the design and context dependencies of massively parallel reporter assays. Nat. Methods 17, 1083–1091 (2020).

54. Wu, Y. et al. Genome-wide fine-mapping improves identification of causal variants. Nat Genet 58, 940–951 (2026).

55. Dickson, S. P., Wang, K., Krantz, I., Hakonarson, H. & Goldstein, D. B. Rare variants create synthetic genome-wide associations. PLoS Biol 8, e1000294 (2010).

56. Wray, N. R., Purcell, S. M. & Visscher, P. M. Synthetic associations created by rare variants do not explain most GWAS results. PLoS Biol 9, e1000579 (2011).

57. Goldstein, D. B. The importance of synthetic associations will only be resolved empirically. PLoS Biol 9, e1001008 (2011).

58. Chang, D. & Keinan, A. Predicting signatures of ‘synthetic associations’ and ‘natural associations’ from empirical patterns of human genetic variation. PLoS Comput Biol 8, e1002600 (2012).

59. Cui, R. et al. Improving fine-mapping by modeling infinitesimal effects. Nat Genet 56, 162–169 (2024).

60. Rosen, J. D., Broadaway, K. A., Brotman, S. M., Mohlke, K. L. & Love, M. I. Higher eQTL power reveals signals that boost GWAS colocalization. Am J Hum Genet 113, 868–876 (2026).

61. Siraj, L. et al. Functional dissection of complex trait variants at single-nucleotide resolution. Nature (2026) doi:10.1038/s41586-026-10121-6.

62. Cheng, J. et al. Accurate proteome-wide missense variant effect prediction with AlphaMissense. Science 381, eadg7492 (2023).

63. Arthur, T. D. et al. Multiomic QTL mapping reveals phenotypic complexity of GWAS loci and prioritizes putative causal variants. Cell Genom 5, 100775 (2025).

64. Chen, M. et al. Cell-type-specific eQTLs underlie the genetic architecture of complex traits. Nature (2026) doi:10.1038/s41586-026-10577-6.

65. Nair, S., Kim, D. S., Perricone, J. & Kundaje, A. Integrating regulatory DNA sequence and gene expression to predict genome-wide chromatin accessibility across cellular contexts. Bioinformatics 35, i108–i116 (2019).

66. Lal, A. et al. Decoding sequence determinants of gene expression in diverse cellular and disease states. Nat Methods 23, 1138–1151 (2026).

67. Zhang, R., et al. Multi-condition and multi-modal temporal profile inference during mouse embryonic development. bioRxivorg (2024) doi:10.1101/2024.03.03.583179.

68. Ji, Y. et al. Scalable and universal prediction of cellular phenotypes. BioRxiv (2024) doi:10.1101/2024.08.12.607533.

69. Umans, B. D., Battle, A. & Gilad, Y. Where are the disease-associated eQTLs? Trends Genet. 37, 109–124 (2021).

70. Kim-Hellmuth, S. et al. Genetic regulatory effects modified by immune activation contribute to autoimmune disease associations. Nat. Commun. 8, 266 (2017).

71. Kent, W. J., Zweig, A. S., Barber, G., Hinrichs, A. S. & Karolchik, D. BigWig and BigBed: enabling browsing of large distributed datasets. Bioinformatics 26, 2204–2207 (2010).

72. Dalla-Torre, H. et al. Nucleotide Transformer: building and evaluating robust foundation models for human genomics. Nat. Methods 22, 287–297 (2025).

73. Pohl, A. & Beato, M. bwtool: a tool for bigWig files. Bioinformatics 30, 1618–1619 (2014).

74. Quinlan, A. R. & Hall, I. M. BEDTools: a flexible suite of utilities for comparing genomic features. Bioinformatics 26, 841–842 (2010).

75. Amemiya, H. M., Kundaje, A. & Boyle, A. P. The ENCODE blacklist: Identification of problematic regions of the genome. Sci. Rep. 9, 9354 (2019).

76. Hingerl, J. C., Karollus, A. & Gagneur, J. Flashzoi: an enhanced Borzoi for accelerated genomic analysis. Bioinformatics 41, (2025).

77. Paszke, A., et al. PyTorch: An imperative style, high-performance deep learning library. arXiv [cs.LG] (2019).

78. Ba, J. L., Kiros, J. R. & Hinton, G. E. Layer Normalization. arXiv [stat.ML] (2016).

79. Kingma, D. P. & Ba, J. Adam: A method for stochastic optimization. arXiv [cs.LG] (2014).

80. Liaw, R. et al. Tune: A research platform for distributed model selection and training. arXiv [cs.LG] (2018).

81. Srivastava, N., Hinton, G., Krizhevsky, A., Sutskever, I. & Salakhutdinov, R. Dropout: A Simple Way to Prevent Neural Networks from Overfitting. Journal of Machine Learning Research 15, 1929–1958 (2014).

82. Wolf, F. A., Angerer, P. & Theis, F. J. SCANPY: large-scale single-cell gene expression data analysis. Genome Biol. 19, 15 (2018).

83. Traag, V. A., Waltman, L. & van Eck, N. J. From Louvain to Leiden: guaranteeing well-connected communities. Sci. Rep. 9, 5233 (2019).

84. Lee, J. et al. Gemini Embedding: Generalizable embeddings from Gemini. arXiv [cs.CL] (2025) doi:10.48550/arXiv.2503.07891.

85. Lee, D. et al. A method to predict the impact of regulatory variants from DNA sequence. Nat Genet 47, 955–961 (2015).

86. Degner, J. F. et al. DNase I sensitivity QTLs are a major determinant of human expression variation. Nature 482, 390–394 (2012).

87. Ovek Baydar, D., et al. JASPAR 2026: expansion of transcription factor binding profiles and integration of deep learning models. Nucleic Acids Res 54, D184–D193 (2026).

88. Benjamini, Y., Drai, D., Elmer, G., Kafkafi, N. & Golani, I. Controlling the false discovery rate in behavior genetics research. Behav. Brain Res. 125, 279–284 (2001).

89. Efron, B. Large-scale simultaneous hypothesis testing: The choice of a null hypothesis. J. Am. Stat. Assoc. 99, 96–104 (2004).

90. Johansson, R., Strålfors, P. & Cedersund, G. Combining test statistics and models in bootstrapped model rejection: it is a balancing act. BMC Syst. Biol. 8, 46 (2014).

91. Ongen, H., Buil, A., Brown, A. A., Dermitzakis, E. T. & Delaneau, O. Fast and efficient QTL mapper for thousands of molecular phenotypes. Bioinformatics 32, 1479–1485 (2016).

92. Benaglio, P. et al. Mapping genetic effects on cell type-specific chromatin accessibility and annotating complex immune trait variants using single nucleus ATAC-seq in peripheral blood. PLoS Genet. 19, e1010759 (2023).

93. Virtanen, P. et al. SciPy 1.0: fundamental algorithms for scientific computing in Python. Nat. Methods 17, 261–272 (2020).

