## Supplementary for "Comprehensive molecular impact mapping of common and rare variants at GWAS loci"

### Supplementary Information

#### Supplementary Notes

Supplemental Note 1. DNACipher-NT-Loc predicts genomic signals from DNA sequence for unobserved cell types and assays

Supplemental Note 2. Long-range sequences or genome locations do not enhance prediction of unobserved experiments

Supplemental Note 3. DNACipher's latent embeddings reflect biological relationships

Supplemental Note 4. DNACipher's imputation enables variant scoring that overcomes cell line ascertainment present in other models

#### Supplementary Figures

Figure S1. DNACipher-NT-Loc signal prediction performance across diverse assay types.

Figure S2. DNACipher train and test experiment allocation for fold 1 of 5 cross validations.

Figure S3. DNACipher-Borzoi signal prediction performance compared with Borzoi.

Figure S4. DNACipher signal imputation outperforms a k-nearest neighbour (kNN) baseline.

Figure S5. DNACipher-Borzoi predicts cell type specific signals.

Figure S6. Benchmarking alternative DNACipher implementations.

Figure S7. DNACipher learns cell type relationships to predict specific genomic signals.

Figure S8. DNACipher learns assay relationships to predict specific genomic signals.

Figure S9. Selected GTEx variants for molecular effect detection benchmark have equal representation across tissues and distances from associated gene (eGene) transcription start sites (TSS).

Figure S10. Saturated In silico mutagenesis (ISM) of caQTL sequences highlights high attribution TF motifs used by DNACipher-Enformer.

Figure S11. Variation in background distribution of cell type, assay, and locus predicted effect sizes from DNACipher-Enformer.

Figure S12. Variation in estimated background distribution with 100 boot-straps.

Figure S13. Example volcano plots showing context-specific significant effects against background variant effect confidence intervals.

Figure S14. T1D impact variants are enriched in accessible chromatin of cell types predicted to be significantly impacted by DNACipher across assays.

Figure S15. Scaling factors and a custom RNA-seq transformation normalize signal measurements to the same mean non-zero signal.

Figure S16. Sampling optimization in ENCODE data identifies optimal experiment weights to minimize cell type and assay ascertainment bias.

**Supplementary References**

**Additional Files**
Table S1. DNACipher training metadata and performance metrics

Table S2. Cell type and assay cluster memberships from latent embedding analysis.

Table S3. Summary statistics from causal and non-causal eQTL comparisons.

Table S4. DNACipher-Enformer DVIM Type 1 Diabetes GWAS summary statistics.

Table S5. T1D DVIM luciferase assay and enrichment statistics (PIP, emVar, snATAC-seq, RAVAR)

Table S6. DNACipher-Enformer DVIM summary statistics across 9 complex traits.

Table S7. DVIM enrichment statistics (PIP, RAVAR)

### Supplementary Notes

#### Supplemental Note 1.
#### DNACipher-NT-Loc predicts genomic signals from DNA sequence for unobserved cell types and assays

We trained several versions of the DNACipher model that take different sequence embeddings as input, with and without genome location information as utilized by the Avocado model[1]. For the initial version of the DNACipher model, to validate the approach while ensuring no possibility of inflated performance metrics due to pre-trained data leakage[2,3,4], we utilized short (6 kb) sequence embeddings from Nucleotide Transformer (NT)[5] which is not trained on genomic signal data. In addition, we included genome locations as model inputs for this initial model, which we named "DNACipher-NT-Loc" (Figure S1A-B). We evaluated the overall performance of DNACipher-NT-Loc by computing the Pearson correlations (R) between predicted and actual experimental values, considering unobserved (test) experiments and unobserved (test) sequences. In this initial model, allocation of test sequences included every second 6 kb sequence within 12 kb genome tiles across chromosomes 2 and 16. This allocation was performed because including genome position as input required training the latent position embedding, requiring example sequences for each position, and therefore we encoded positions at a 12 kb resolution, including additional location encodings at 48 kb and 120 kb resolutions (Figure S1A). Furthermore, in this initial model we did not perform cross-validation, but had a single allocation of train/test sequences and train/test experiments. Test experiments were chosen randomly amongst the cell types measured for each assay. We obtained Pearson correlations of R = 0.68 for predicting training experiments (N = 2,676 experiments) from test sequences (N = 54,440) and R=0.67

for predicting unobserved test experiments (N = 206) from training sequences (N = 1.3 x $10^6$) (Figure S1C. We next evaluated the performance of DNACipher-NT-Loc on test sequences and unobserved (test) experiments, and obtained a Pearson correlation of R=0.60 (Figure S1C). Overall, this demonstrates that DNACipher-NT-Loc can impute experimental signals even when the sequence and experiment has not been observed.

We also considered performance of DNACipher-NT-Loc by assay type (Figure S1D, Table S1). For chromatin accessibility assays (ATAC-seq and DNase-seq), we observed consistently high Pearson correlations (R≈0.50) across observed/unobserved experiment and sequence stratifications (Figure S1D, Table S1). For RNA-seq assays (total RNA-seq, mRNA-seq, and polyA depleted mRNA-seq), we observed correlations of R≈0.70 across testing stratifications (Figure S1D, Table S1). Histone modification ChIP-seq assays were more variable. Certain marks (H3K4me3, H3K79me3, H3K9ac, H3K4me2, H3K27ac, and H3K36me3) achieved Pearson correlations of 0.40-0.64, while others achieved lower correlations of ~0.20 (Figure S1D, Table S1). TF ChIP-seq prediction performance was generally consistent across stratifications (Figure S1D, Table S1), however, certain TFs had substantially reduced performance when they were predicted in unobserved cell types and sequences (PATZ1, ZBTB8A, TRIM22, ZNF274, REST, GATA3, MYNN, ZNF707, and YBX1) (Figure S2D, Table S1). These results show that DNACipher-NT-Loc was able to infer signal measurements in unobserved cell types across a diverse range of assays, including for unobserved sequences, and has consistently high performance for chromatin and gene expression assays.

**Supplemental Note 2.**

**Long-range sequences or genome locations do not enhance prediction of unobserved experiments**

We systematically compared DNACipher implementations with different combinations of short or long sequence or genome location on model inputs (Figure S6A). We also examined whether there was a difference in performance when the genome location inputs for the test chromosomes had example training sequences or not (Figure S6A). This latter condition acted to isolate whether the performance difference due to genome location was because of improving learnt relationships between cell types and assays via the genome location input or from the sequence information.

For test sequences and train experiments, DNACipher models with long sequences did significantly outperform DNACipher with short sequences and genome location input or only short sequence or only genome location (DNACipher-NT-Loc, $p > 0.01$, paired t-test) (Figure S6B, Table S1). However, for test experiments using train sequences DNACipher with long sequence context did not outperform short sequences with genome location, but did outperform short sequences alone (Figure S6B, Table S1). For test experiments and test sequences, the longer sequence models did not outperform the short sequence models with or without genome location information (Figure S6B, Table S1). DNACipher-NT-Loc had better performance for test sequences in train experiments compared with DNACipher-NT-UntrainedLoc, suggesting the genome location input improved signal prediction for unobserved sequences at similar genome locations. However there was no significant difference between DNACipher-NT-Loc and DNACipher-NT-UntrainedLoc when comparing results for test experiments, suggesting the genome location information did not provide additional information to improve imputation of unobserved experiments (Figure S6B, Table S1). Comparing the short sequence models with or without genome location information, genome location did improve signal prediction for train experiments and test sequence, as well as test experiments and train sequences, but did not improve signal prediction for test experiments and test sequences (Figure S6B, Table S1). All sequence models outperformed genome location only models (Figure S6B, Table S1).

While these results are consistent with Avsec *et al.*[3] showing that long sequence does improve signal prediction for test sequences in train experiments, it does not outperform encoding short sequences with genome location when performing imputation (test sequences and test experiments). Furthermore, neither long sequence context or genome location context outperforms encoding short 6 kb sequences for inferring signals for test sequences and test experiments. Despite these observations, it is important to note that long range signal effect prediction of a genetic variant is only possible for long sequence context modelling, and hence the long sequence models were utilized for further analysis.

#### Supplemental Note 3. DNACipher's latent embeddings reflect biological relationships

To assess whether DNACipher learned biologically meaningful patterns, we extracted latent representations of cell types and assays from the DNACipher-Enformer model's input layers. We applied Leiden clustering to group the cell types and assays based on their latent embeddings, and annotated them based on biological themes (Figure S7,S8, Table S2). We then applied Uniform Manifold Approximation (UMAP) to visualize these embeddings in two dimensions (Figure S7,S8).

In the cell type latent space, we observed clusters reflecting expected biological relationships (Figure S7, Table S2). For example, cluster 4 was primarily composed of nervous system tissues (including brain regions) and was closely connected to cluster 16 that was primarily composed of neural progenitor cell types (Figure S7). Within the assay embedding space, we identified clusters that matched known regulatory complexes (Figure S8). For example, cluster 25 contained histone modifications and factors associated with the Polycomb repressive complex 2 (PRC2) including EZH2, SUZ12, and H3K27me3 ChIP-seq (Figure S8, Table S2) and cluster 1 highlighted AP-1 and stress-response TFs including ChIP-seq assays for JUN, FOS, NFIC, and RXRA (Figure S8, Table S2). This shows that DNACipher leverages biological relationships between assay types to infer missing assay measurements.

Overall, the latent representations learned by DNACipher align with known biology, even though the available training data included a high level of missing data for cell type and assay combinations.

#### Supplemental Note 4. DNACipher's imputation enables variant scoring that overcomes cell line ascertainment present in other models

To assess the effect of including additional context predictions from DNACipher for DVIM, we performed a sensitivity analysis comparing the DVIM method utilising DNACipher-Enformer with either the targeted contexts in specific T1D relevant biosamples (T cells, B cells, NK cells, and pancreatic tissue; N = 36) or utilizing all of the possible outputs ("DNACipher-Enformer all"; N = 38,582) (Extended Data Figure 7A). In addition, we performed the same analysis using Borzoi with the available experiments that matched the DNACipher-Enformer targeted analysis (N = 34), and all of the Borzoi outputs (N = 7,611). Because of differences in multiple hypothesis testing (MHT) burden between each of these analyses and vastly different experiment outputs, we compared the mean posterior inclusion probability (PIP) for credible set variants ranked by quantiles of N significant effects detected across outputs and at different significance thresholds (z-cutoff, Extended Data Figure 7A). We then computed the area under the curve (AUC) for the mean PIP as a measure of performance of each approach for ranking likely causal variants by number of significant effects for each significance threshold cutoff (Extended Data Figure 7A).

This approach revealed that DNACipher-Enformer with all outputs had the best performance across all significance thresholds, followed by DNACipher-Enformer targeted, Borzoi targeted, and Borzoi with all outputs (Extended Data Figure 7B-C). Hence we observe that adding additional assay predictions balanced across all cell/tissue types represented in the DNACipher output improves ranking of likely causal variants, while including all Borzoi contexts had a negative effect. This is because including all Borzoi outputs are strongly biased toward cell lines like K562, HEK293, and HepG2 cell line that are irrelevant for T1D, where we expect immunity-related cell/tissue types to be relevant given the auto-immune origin of T1D[6] (Extended Data Figure 7D). The DNACipher model has outputs reflecting predictions that are balanced across all tissue types due to the imputation capability, which enabled the prioritization of variants by immune related cell/tissue types (Extended Data Figure 7D).

This analysis suggests that including additional contexts from DNACipher is beneficial due to the imputation capability, but is not so for models where the outputs reflect strong ascertainment toward potentially irrelevant cancer cell lines. Because of the MHT burden of additional tests and compute for additional context predictions, our larger scale analysis remains on the focused contexts.

### Supplementary Figures

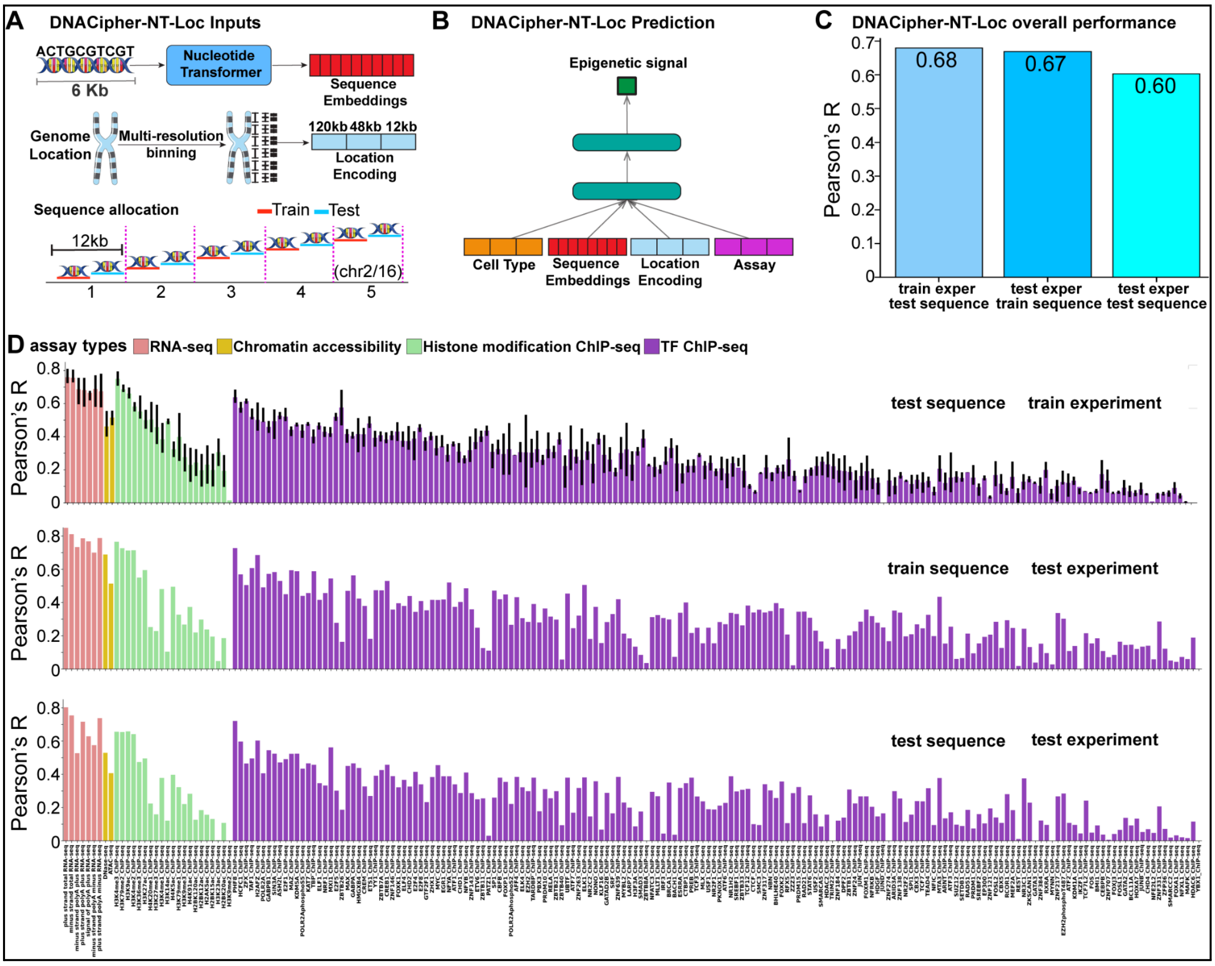


**Figure S1. DNACipher-NT-Loc signal prediction performance across diverse assay types. A.** Schematic of the DNACipher-NT-Loc model inputs, which include sequence embeddings for 6 kb sequences from the Nucleotide Transformer model, location of the input sequence in the genome (provided as a one-hot-encoding at 120 kb, 48 kb, and 12 kb bins within each chromosome), and the allocation of train and test sequences within each 12 kb genome bin on chromosomes 2 and 16. **B.** Depiction of the DNACipher-NT-Loc model, which takes the inputs depicted in A, to predict the mean genomic signal in the centre 300 bp of the inputted 6 kb sequence. **C.** Pearson's correlation (R) between the predicted and observed genomic signals stratified by training sequences in testing (unobserved) experiments, test sequences in training (observed) experiments, and test sequences in test experiments. **D.** Pearson's correlations of predicted versus actual signal values when stratified by assay type (x-axis) and by testing stratification. Error bars indicate the interquartile range of correlations across cell types. For test experiment stratifications, there is one held-out experiment per assay type, so error bars are not depicted.

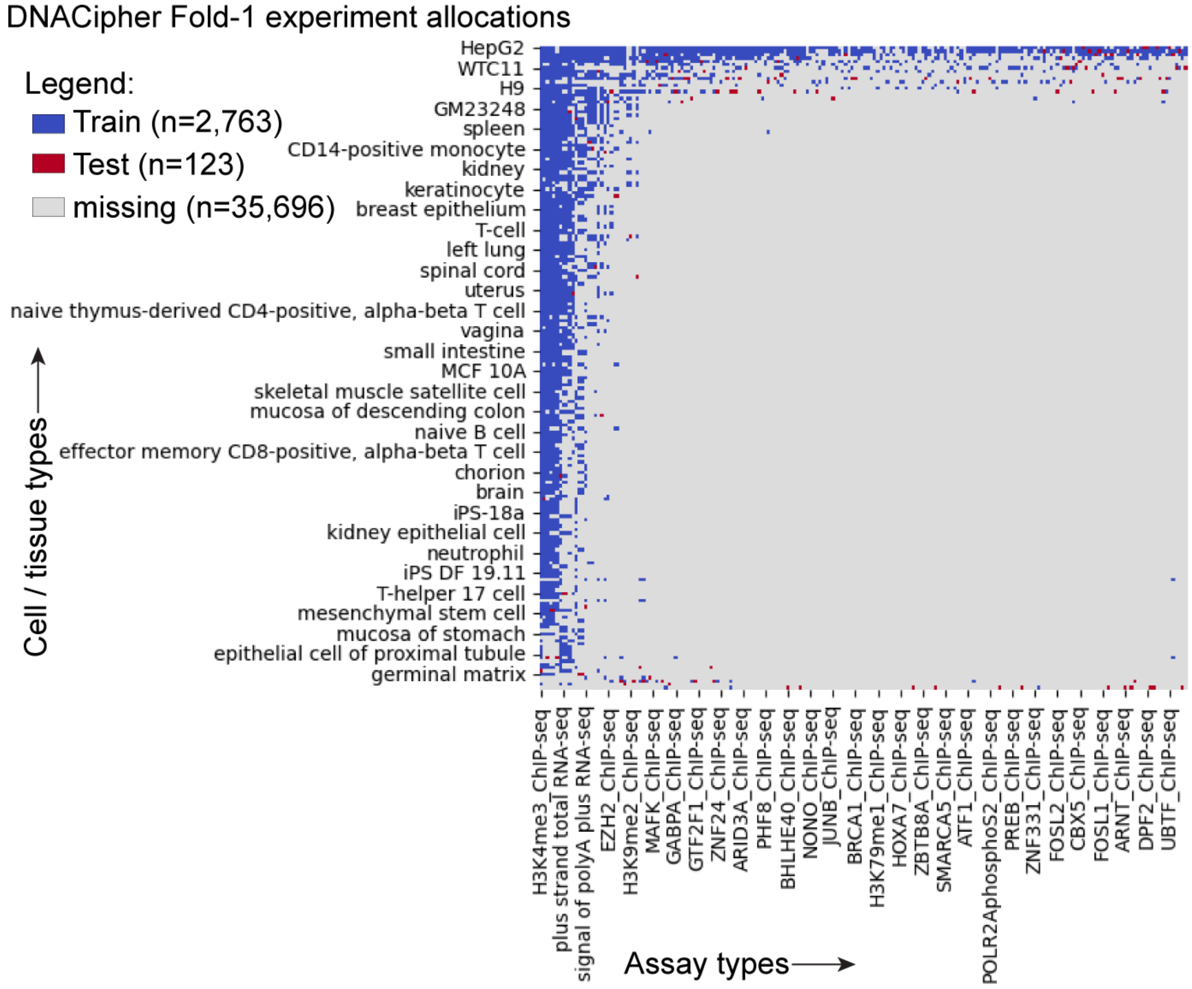


**Figure S2. DNACipher train and test experiment allocation for fold 1 of 5 cross validations.** The cell type/assay matrix of observed (train) and unobserved (test) experiments from ENCODE for fold-1 of the DNACipher 5-fold cross-validation. This shows two important points; 1) the high ascertainment bias for assays performed on particular cell types such as K562 cells, and 2) the approach to select test cell types/experiments for each assay that have the fewest other assays sampled for that cell type, thereby representing performance for the majority of cell types and assays which have a sparse number of experiments.

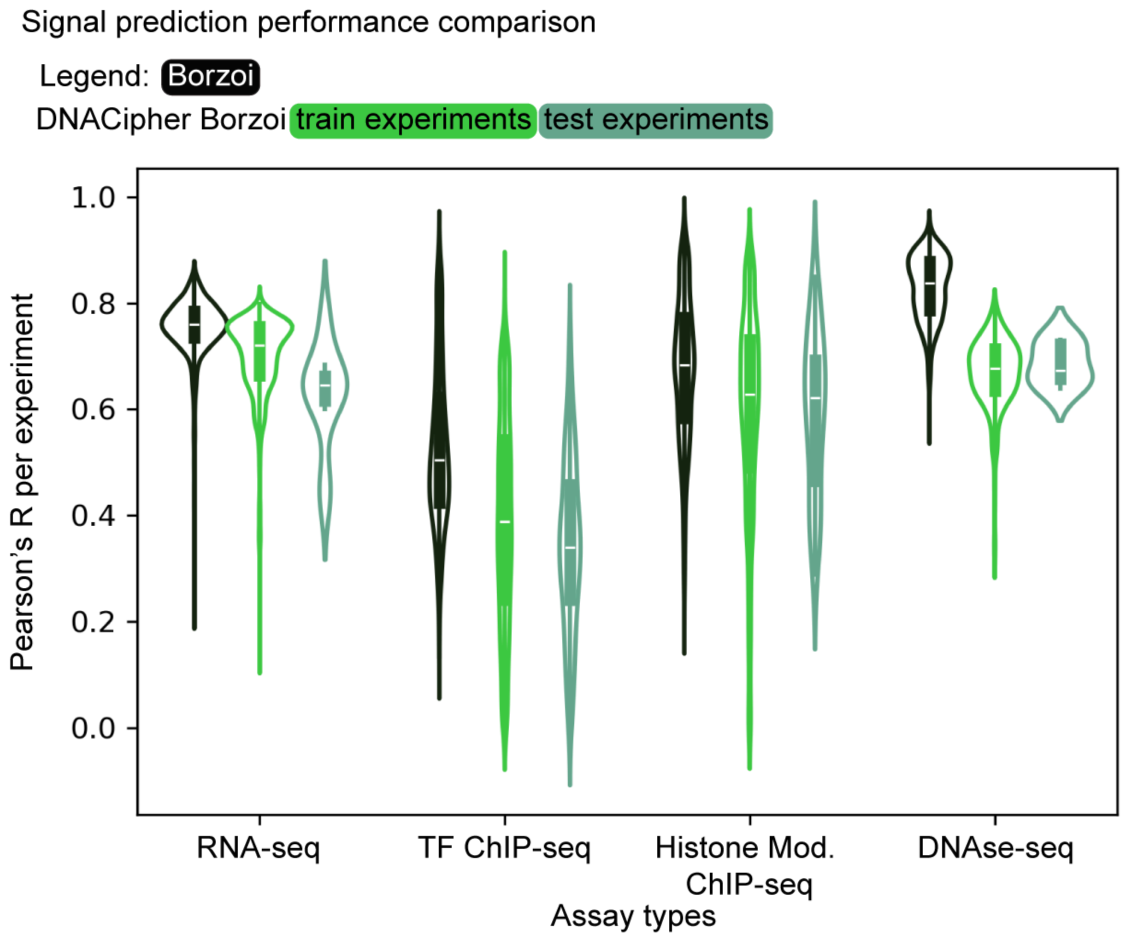


**Figure S3. DNACipher-Borzoi signal prediction performance compared with Borzoi.** Violin plots show the distribution of Pearson's R (y-axis) between predicted and observed signal values across all experiments for different assay types (x-axis), with predictions from Borzoi, DNACipher-Borzoi for training experiments, and DNACipher-Borzoi for unobserved (test) experiments.

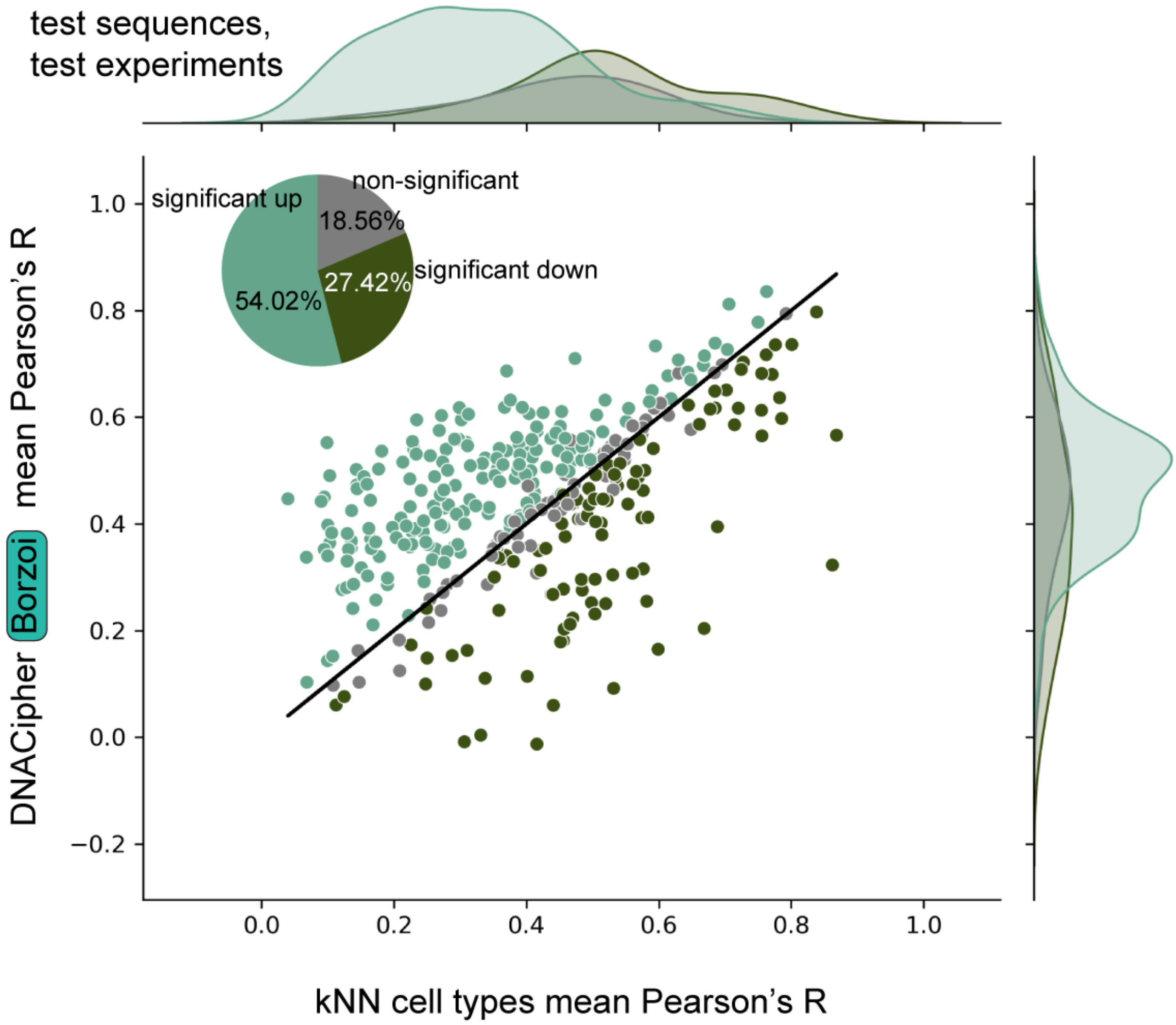


**Figure S4. DNACipher signal imputation outperforms a k-nearest neighbour (kNN) baseline.** Scatter plot; the x-axis is the mean Pearson's R (mean across N = 1,350 random batches of 256 sequences) from correlating the mean signal of kNN cell types (at most k = 10, and varies by assay type due to missing experiments) that are measured for a given assay against the observed signals for the same assay in a given cell type (test experiments only) for test sequences. The y-axis is the same metric, except using signal predictions from DNACipher-Borzoi. Test experiment correlations from all 5 cross validations of train/test experiment allocations are shown (N = 361), which differs from Figure 2C where each cross-validation is shown separately. The diagonal line indicates the expectation if the x- and y- axis values were the same. The overlaid pie chart indicates the proportion of assays where DNACipher-Borzoi correlations across batches (N = 1,350 of 256 sequences) were significantly higher than the kNN baseline correlations, significantly lower than the kNN baseline, or non-significantly different to the kNN baseline correlations. Significance was determined by a two-sided paired Wilcoxon rank-sum test with Benjamini-Hochberg false discovery rate $p < 0.05$. Density plots are depicted adjacent to the relevant axis, and depict distributions of the values for the axis stratified by experiments significantly greater than, significantly less than, or non-significantly different to the kNN baseline.

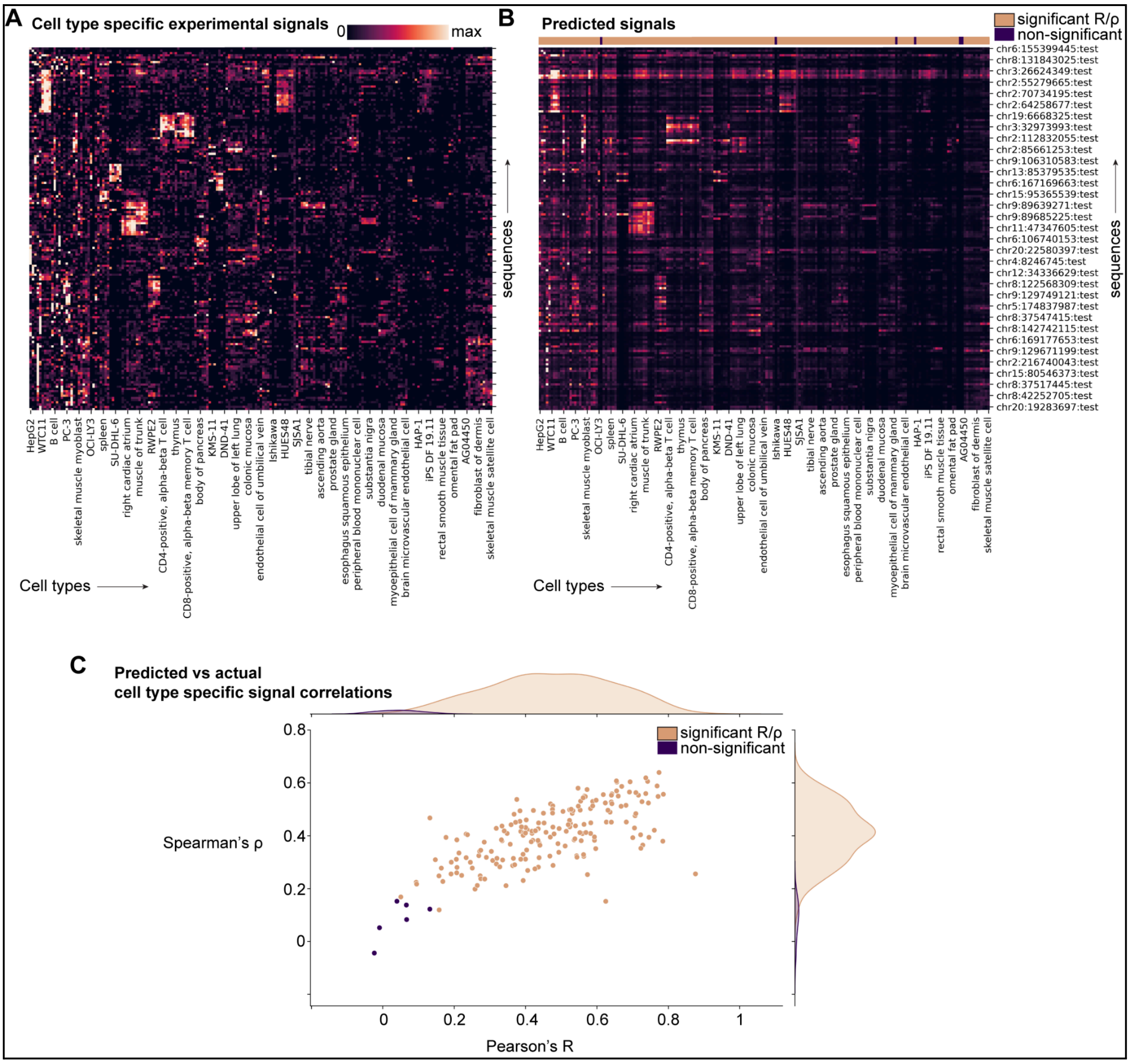


**Figure S5. DNACipher-Borzoi predicts cell type specific signals. A.** Heatmap where the rows are selected test sequences (unobserved during DNACipher training) with high cell-type specific signals (N = 162), and the columns are cell types (N = 191). Values represent the sum of signals across all assays available for that particular cell type. **B.** The equivalent to A, except the signals predicted using DNACipher-Borzoi. Columns are annotated by cell types where the signal predictions across all test sequences were significant or non-significantly correlated to the observed signals using both Pearson's R and Spearman's ρ. P-values were determined using a paired two-sided test with the beta-distribution for Pearson's R and the t-test for Spearman's ρ (default scipy.stats parameters), and taken as significant if the Benjamini-Hochberg false discovery rate was smaller than 0.05 for both correlation statistics. **C.** Scatterplot showing Pearon's R (x-axis) and Spearman's ρ (y-axis) for each cell type (N = 191) with labels for significant correlations by both statistics. Density plots are depicted adjacent to the relevant axis, and depict distributions of the values for the axis stratified by cell types with significant correlations for both R and ρ or not.

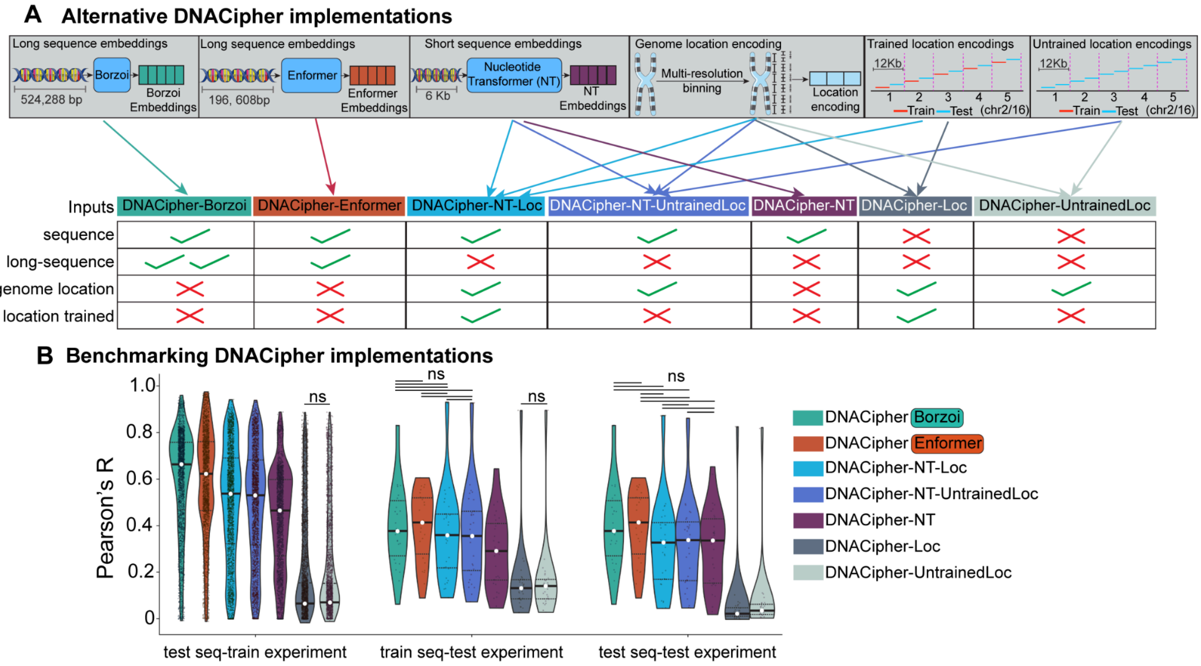


**Figure S6. Benchmarking alternative DNACipher implementations. A.** Schematic of alternative DNACipher implementations, with different possibilities for encoding the DNA sequence, including long versus short sequencing embeddings (Enformer or Nucleotide Transformer (NT)), encoding the position of the genome along with DNA sequence, and whether held out test sequences have adjacent training sequences. Colored arrows between the possible input combinations to the corresponding DNACipher implementation name indicate the combination of inputs used for each model. **B.** Violin plots indicating Pearson's R on the y-axis, separated by train/test sequence and test/train experiment combinations. Each point is the Pearson's R for a single experiment within the respective stratification. Colors indicate the particular DNACipher version used to make the signal predictions. Models that are non-significantly different from one another are indicated (Benjamini-Hochberg adjusted $p \geq 0.01$, paired t-test). Means are indicated with a white point in the centre of each violin plot.

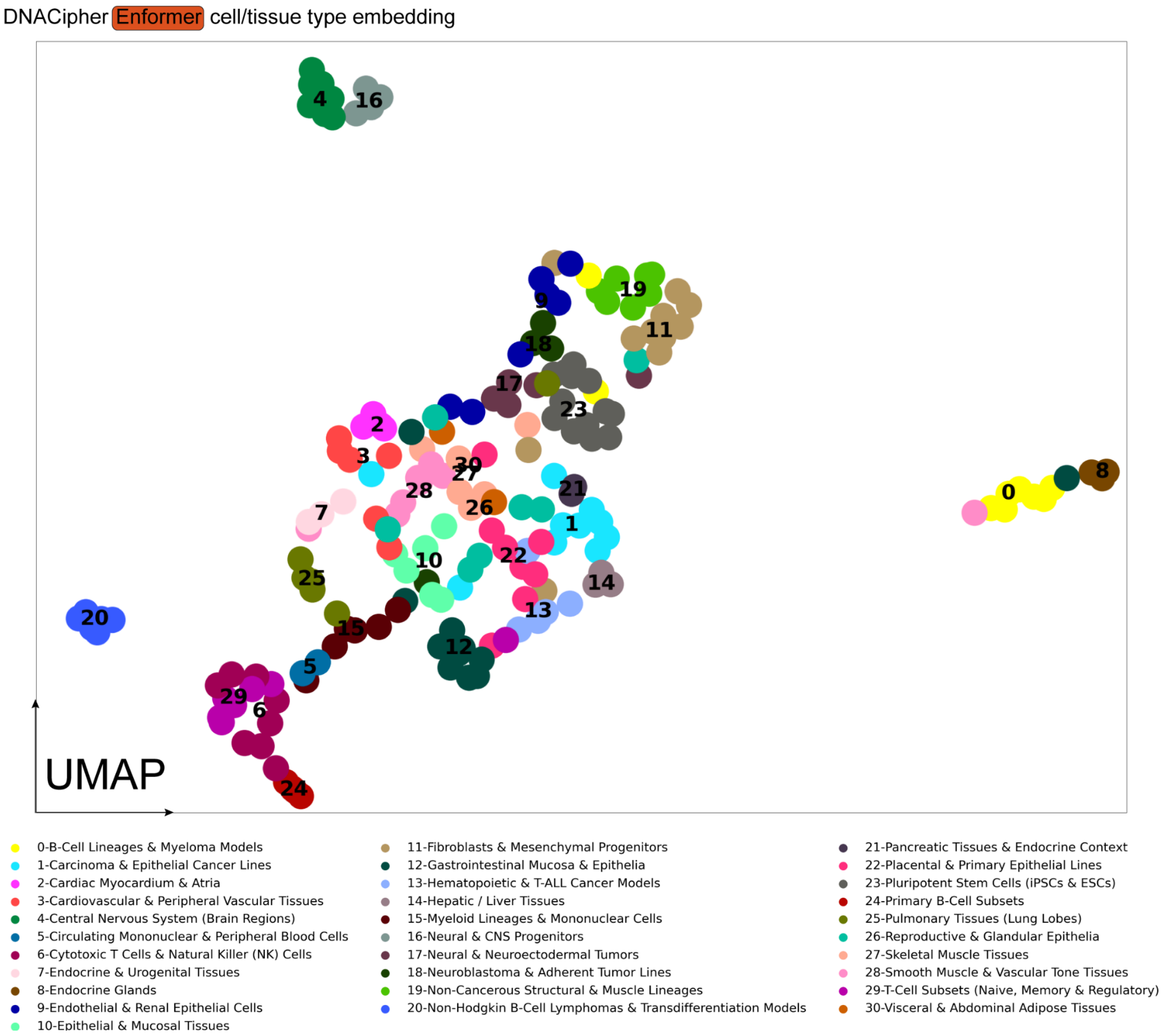


**Figure S7. DNACipher learns cell type relationships to predict specific genomic signals.** UMAP of 191 cell / tissue types clustered in the DNACipher cell type embedding space. Clusters reflected similar biology, as annotated in the legend. Exact cell/tissue types within each cluster are listed in Table S2.

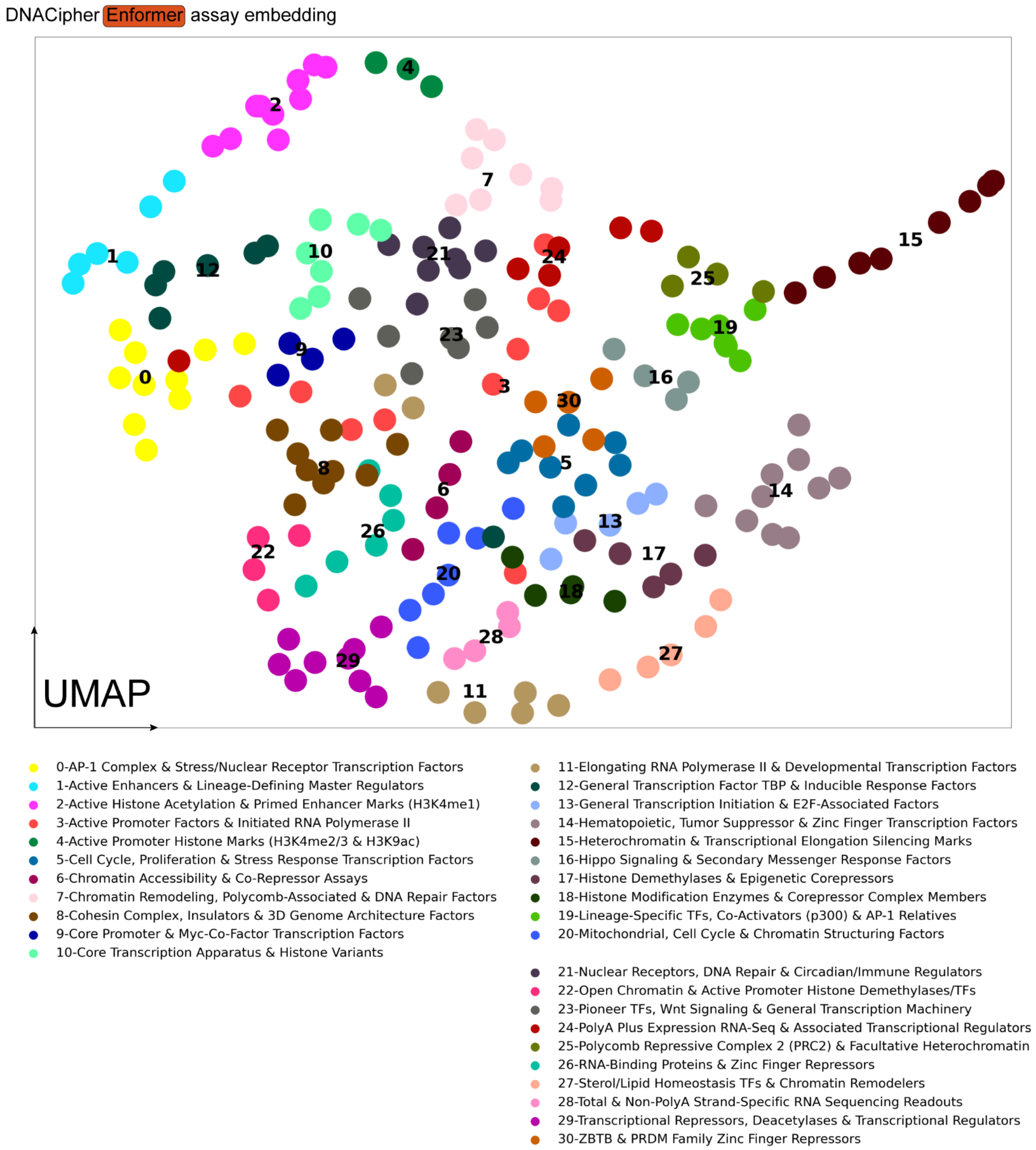


**Figure S8. DNACipher learns assay relationships to predict specific genomic signals. A.** Uniform Manifold Approximation (UMAP) based on cosine distances between assays in the DNACipher assay embedding space. Clusters represent assays with similar DNA regulatory activity or function, as annotated in the legend. Exact assays within each cluster are listed in Table S2.

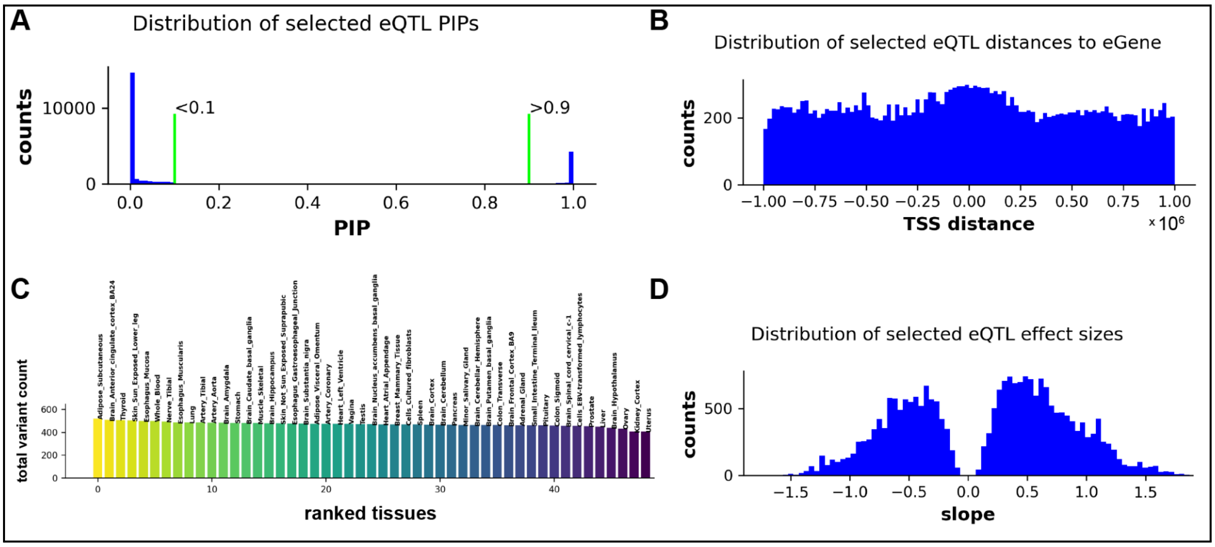


**Figure S9. Selected GTEx variants for molecular effect detection benchmark have equal representation across tissues and distances from associated gene (eGene) transcription start sites (TSS). A.** Distribution of selected Genotype-Tissue Expression project (GTEx) v8 eQTL posterior inclusion probabilities (PIPs) from DAP-G fine-mapping. Vertical lines indicate the threshold used to label variants as non-causal (PIP<0.1) or likely causal (PIP>0.9). **B.** Distribution of distances between the selected eQTLs and the eGene TSS. **C.** Bar plot showing the number of selected eQTLs in 49 tissues collected from GTEx v8. The x-axis ranks the tissues by eQTL count, and the y-axis shows the total number of selected variants. **D.** Distribution of effect sizes for the selected eQTLs on the associated eGene.

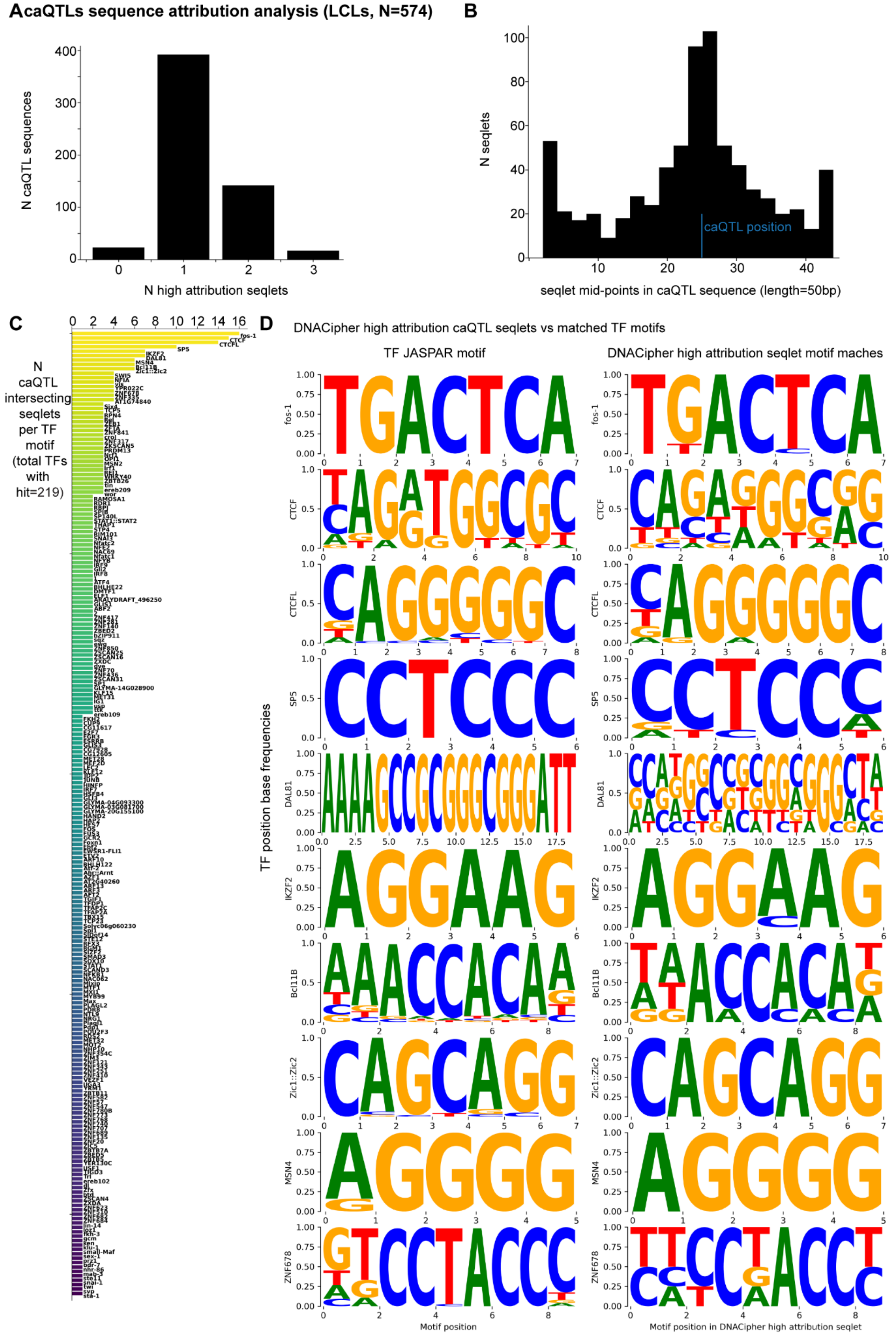


**Figure S10. Saturated *In silico* mutagenesis (ISM) of caQTL sequences highlights high**

**attribution TF motifs used by DNACipher-Enformer. A.** Bar chart showing the number of 50 bp sequences surrounding lymphoblast cell line (LCL) caQTLs (N = 574) with seqlets that have a high sum of predicted chromatin accessibility effects in LCLs (attribution) from DNACipher-Enformer using saturated in silico mutagenesis. Seqlets were defined using tangermeme[7] on the DNACipher-Enformer position attributions. **B.** Bar chart showing the number of high attribution seqlets detected across the 50 bp sequences surrounding caQTLs (y-axis) across the sequence positions (x-axis). **C.** Ranked bar chart showing JASPAR non-redundant core TF motifs ranked (y-axis) by the number of caQTL intersecting seqlets they were matched to via tangermeme[7] (x-axis). Blue vertical line indicates the caQTL positions in the centre of the sequences. **D.** For the top 10 TF motifs ranked in C, the left column shows sequence logos for the position frequency matrix from JASPAR, depicting at each TF motif position (x-axis) the proportion of each DNA nucleotide observed at that position. The right column shows the equivalent, except based on DNACipher-Enformer high attribution caQTL intersecting seqlets that matched to the same TF motif. This shows high correspondence between DNACipher-Enformer predicted important sequences and known TF motifs.

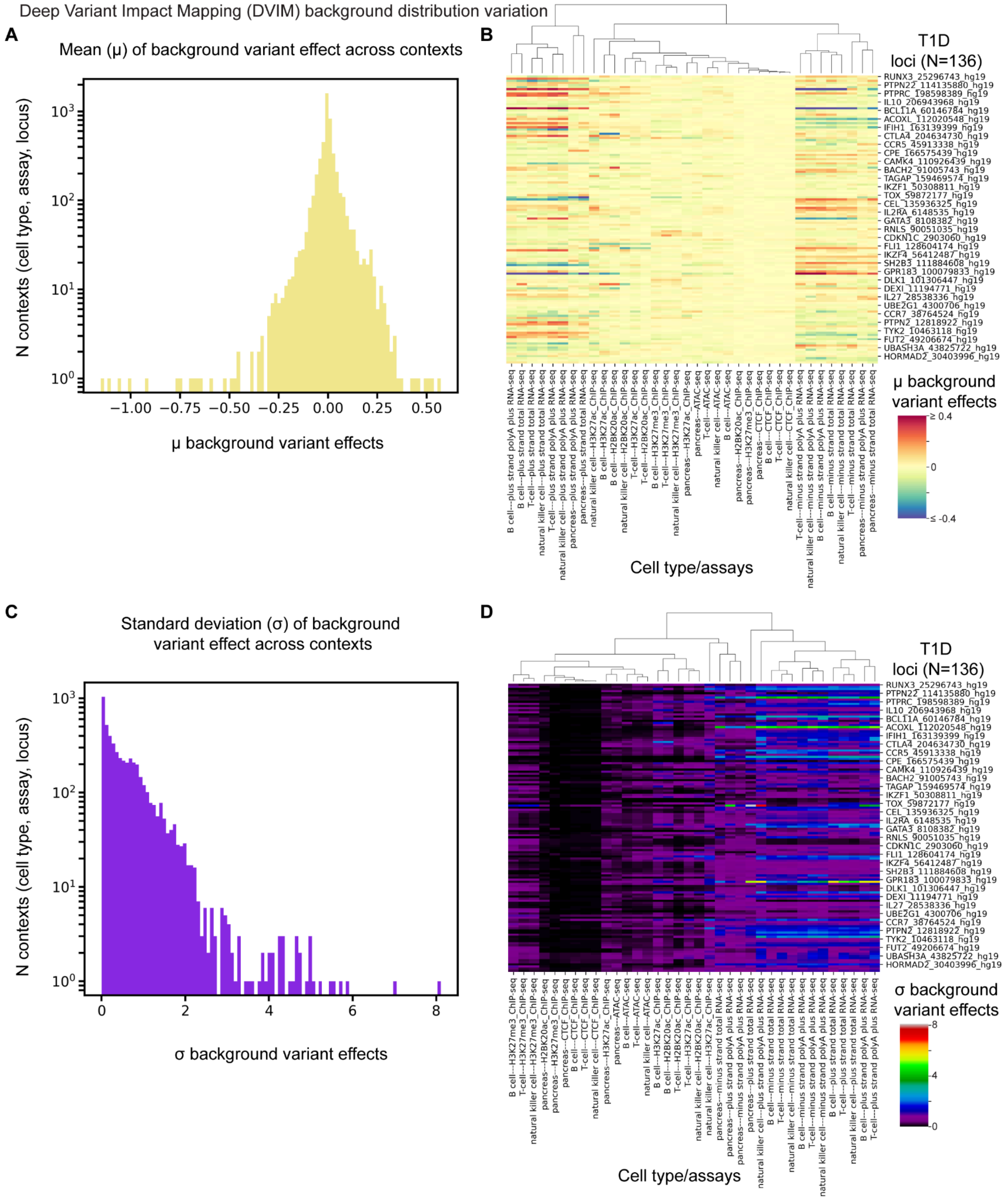


**Figure S11. Variation in background distribution of cell type, assay, and locus predicted effect sizes from DNACipher-Enformer. A.** Distribution of the mean of predicted non-trait associated variant ('background' variant) effects for a specific context (cell type, assay, and T1D GWAS locus), showing high variance depending on the precise context. **B.** Heatmap where rows are T1D loci (N = 136), and columns are cell type/assays (relevant T1D contexts predicted, N = 36), and values are the mean predicted background variant effects, illustrating high locus, cell type, and assay specificity in background variant effect means. **C.** Distribution of the standard deviation (σ) of predicted background variant effects for a specific context (cell type, assay, and T1D GWAS locus), showing high variance in σ depending on the precise context. **D.** Heatmap where rows are T1D loci (N = 136), and columns are cell type/assays (relevant T1D contexts predicted, N = 36), and values are the σ of predicted background variant effects, illustrating high locus, cell type, and assay specificity in background variant effect σ. Overall, this indicates the distribution of background variant effects is highly context dependent, supporting the statistical decision to utilize a background distribution that is specific to each cell type, assay, and locus.

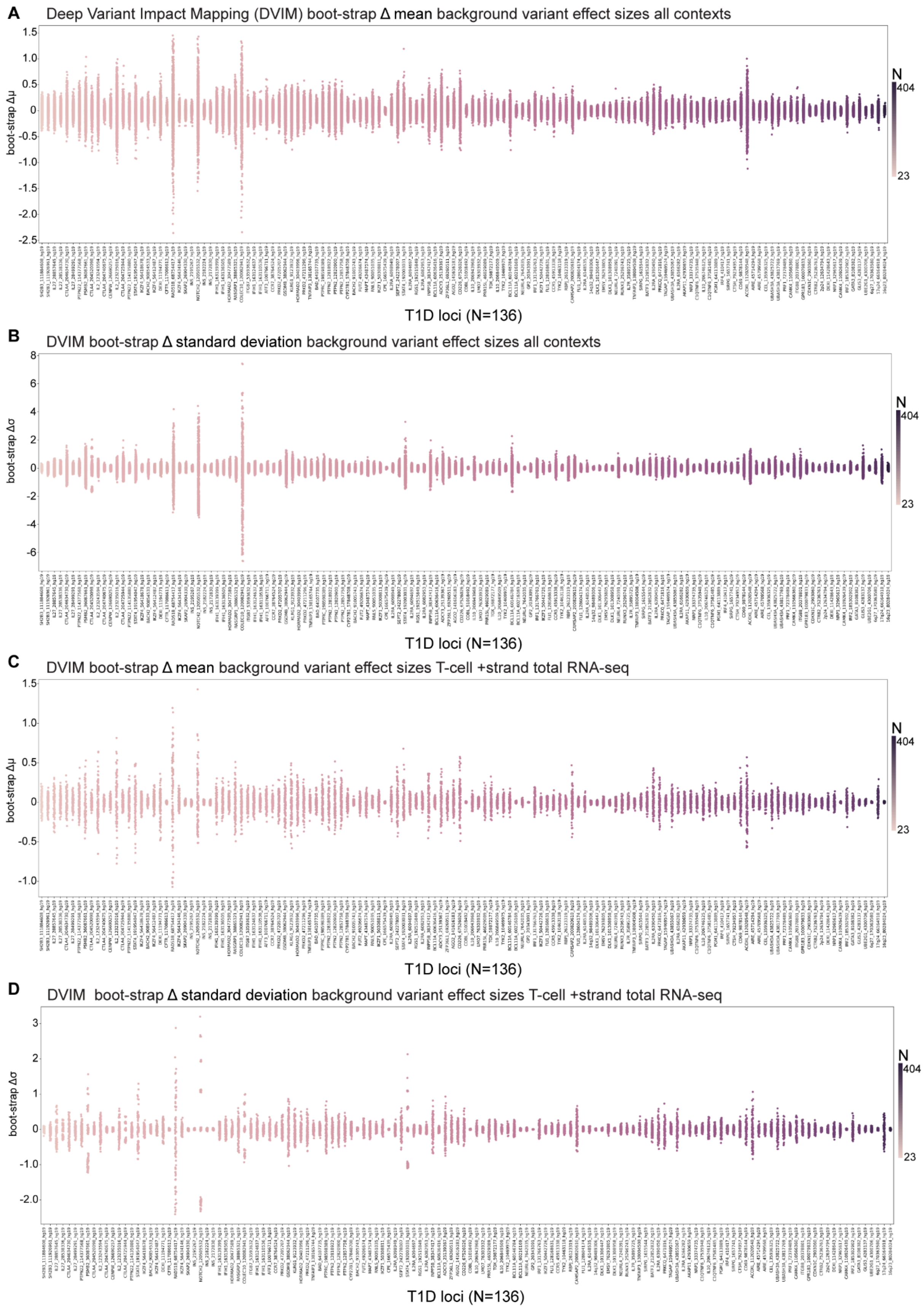


**Figure S12. Variation in estimated background distribution with 100 boot-straps. A.** Strip plots displaying T1D GWAS loci ranked (N = 136, x-axis) by the number of non-trait associated variants ('background variants') ±57,344 bp from the locus lead causal variant position. The y-axis is the change in mean predicted effects for a given cell type and assay (boot-strap Δμ) from boot-strapping the background variants (N = 100). This demonstrates high locus specificity in the uncertainty in the mean of background variant effects. **B.** Equivalent to A, except the y-axis is the change in standard deviation of predicted effects for a given cell type and assay (boot-strap Δσ) from boot-strapping the background variants (N = 100). This demonstrates high locus specificity in the uncertainty of the σ of background variant effects. **C,D.** Equivalent to A and B, respectively, except only showing for T-cell +strand RNA-seq predictions, showing the high specificity of the locus variability in background variant effects is still persistent even when considering a specific cell type/assay context. Overall, this justifies the boot-strapping approach taken to account for uncertainty in the cell type, assay, and locus specific background variant effect distributions.

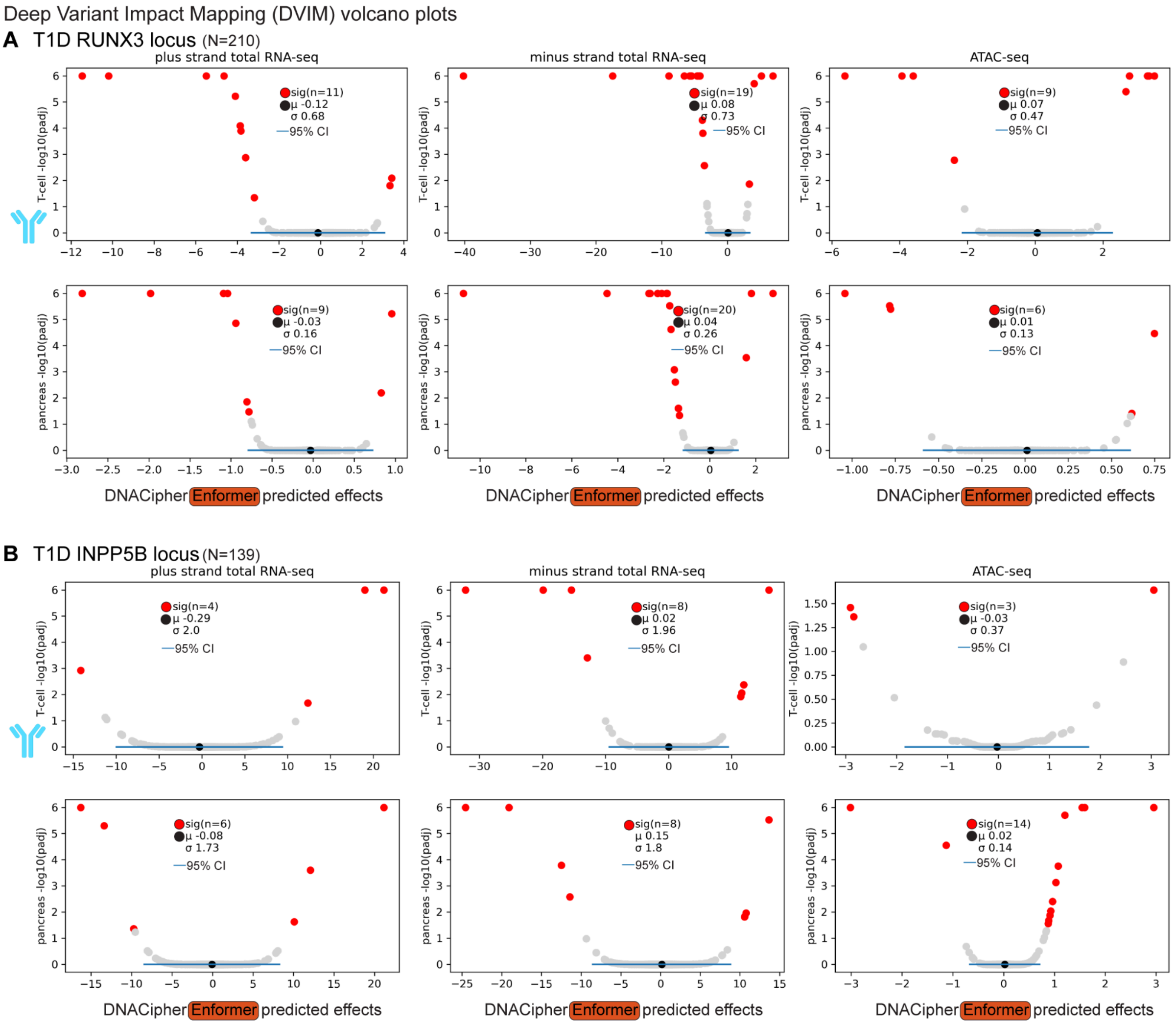


**Figure S13. Example volcano plots showing context-specific significant effects against background variant effect confidence intervals. A.** Volcano plots for different tested contexts from the T1D DVIM analysis, arranged so that each column of volcano plots represents a different assay (plus strand total RNA-seq, minus strand total RNA-seq, and ATAC-seq), and each row represents a different cell/tissue type (T- cells and pancreas). Each point of the plots are DVIM tested variants (common or rare) at the *RUNX3* locus (N = 210), and the y-axis of each plot is the -log10(adjusted p-value) of significance from the

DVIM boot-strapping based test, and the x-axis is the predicted DNACipher-Enformer effects. Variants that were significant for the particular context are highlighted in red, and non-significant variants in grey. The black dot is the mean of the predicted effects in the same context for background variants, and the horizontal blue line is the 95% confidence intervals of those background variant effects. The standard deviation of the background variant effects are also indicated in text. **B.** Equivalent to A, except for the *INPP5B* T1D GWAS locus.

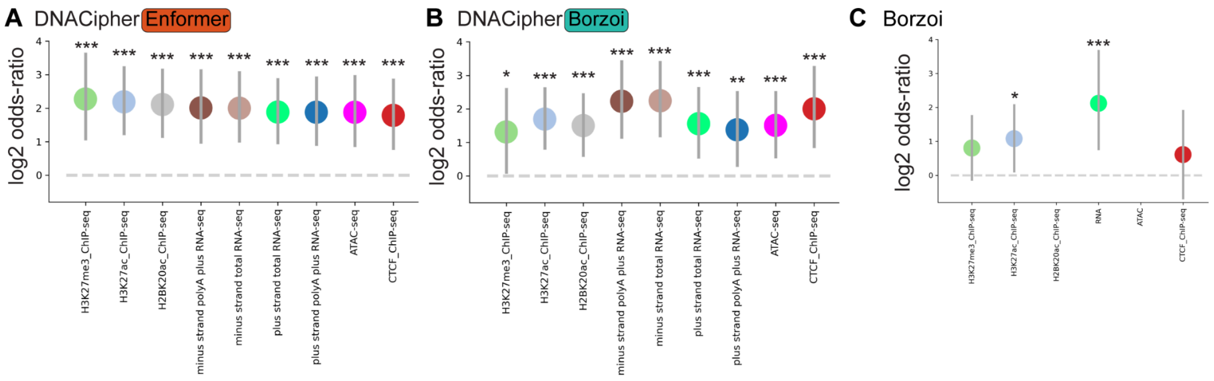


**Figure S14. T1D impact variants are enriched in accessible chromatin of cell types predicted to be significantly impacted by DNACipher across assays. A.** Dot plot with the log2-odds-ratio of overlap between the cell types predicted to be affected by imVars from DNACipher-Enformer DVIM within each molecular assay type, and variants found within accessible chromatin of the same cell type. The horizontal dotted line indicates the random expectation, and the error bars indicate the adjusted 95% confidence intervals for the log2-odds-ratio. P-values are determined based on Fisher's exact test between imVars with significant effects for the relevant assay and found in accessible chromatin of the cell type predicted to be affected. Benjamini-Hochberge false discovery rate (FDR) was used to correct for multiple hypothesis testing. ***FDR < 0.001, **FDR < 0.01, *FDR < 0.05. **B.** Equivalent to A, except for DNACipher-Borzoi predictions. **C.** Equivalent to A, except for Borzoi predictions. Exact N, p-values, contingency tables and other summary statistics are provided in Table S3.

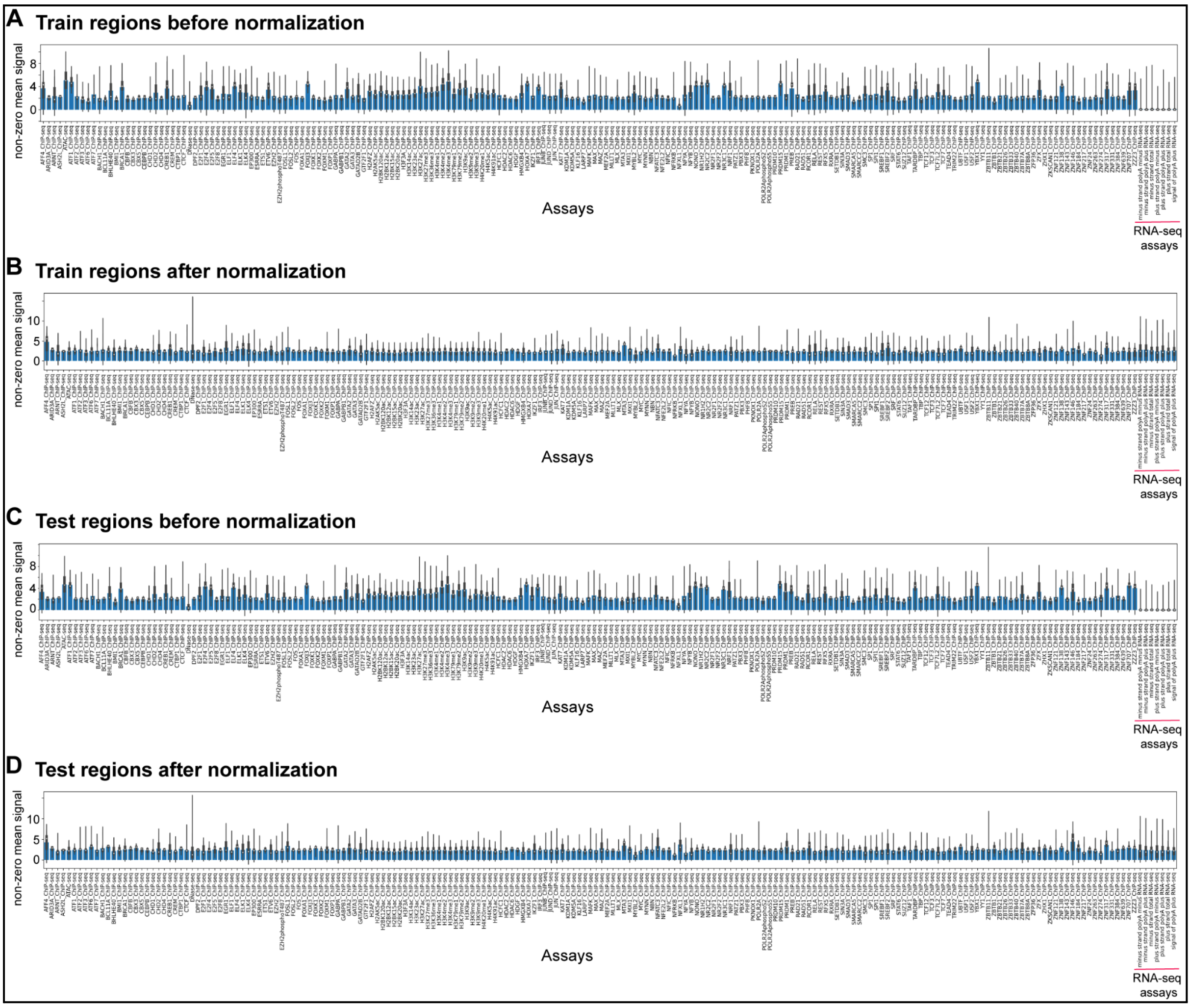


**Figure S15. Scaling factors and a custom RNA-seq transformation normalize signal measurements to the same mean non-zero signal. A.** Bar-charts showing the mean un-normalized non-zero mean signals (y-axis) across training regions for each assay (x-axis). RNA-seq assays are highlighted. Overlaid box-plots display the median value, edges are 25th and 75th percentiles, and whiskers are 1.5X the interquartile range. **B.** Same as panel A, except after applying (1) a custom transformation to the RNA-seq assays and (2) a global re-scaling of all assays to align the same mean non-zero signal. **C.** Equivalent to A, but for the test regions. **D.** Equivalent to B, except the RNA-seq transformation and subsequent scaling factors learnt on the training data are applied to the test data here to control for potential leakage of information between train and test data via normalization.

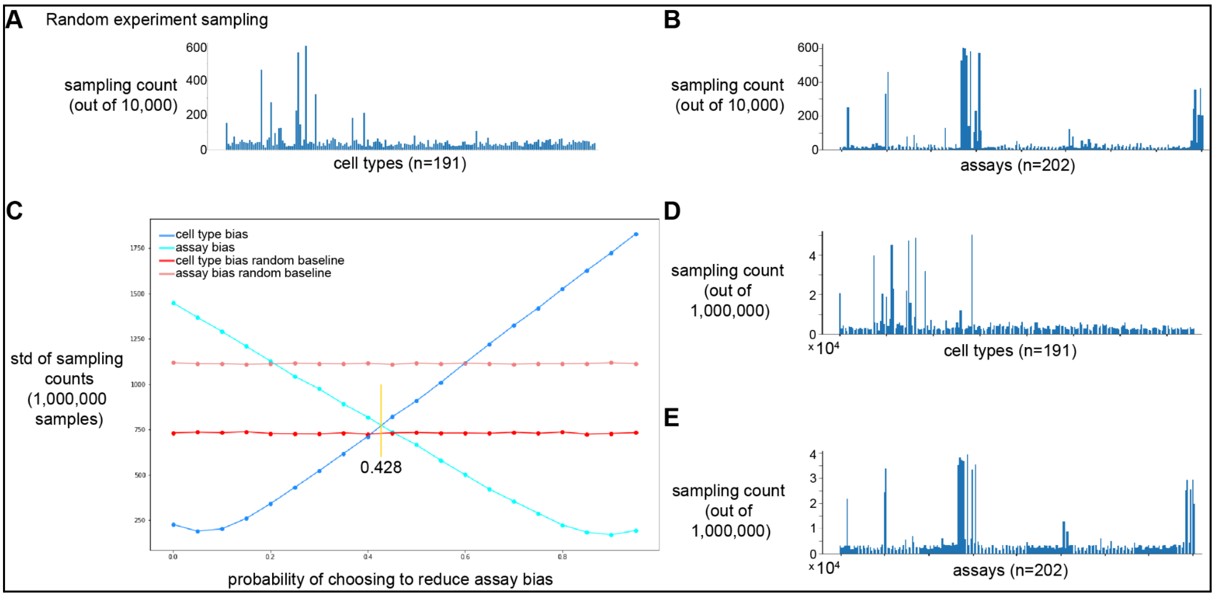


**Figure S16. Sampling optimization in ENCODE data identifies optimal experiment weights to minimize cell type and assay ascertainment bias. A.** Bar chart indicating the count of each cell type sampled by randomly sampling experiments (N = 10,000 times) from a total of 2,882 experiments. This indicates non-uniform representation of cell types across experiments. **B.** Equivalent to A, except displaying the count of each assay type, also indicating non-uniform representations of assays amongst experiments. **C.** Line plot where the y-axis is the standard deviation of sampling counts across cell types or assays (using 1,000,000 random experiment samples). The x-axis indicates the probability of choosing to minimise assay bias or cell type bias. If choosing to minimize assay bias, experiments are sampled with a probability inversely proportional to the frequency of the assay across all experiments. At each probability, the standard deviation indicates both the cell type ascertainment bias and assay ascertainment bias observed for the sampling procedure. Random sampling is also indicated for cell types and assays as a baseline comparison. An optimal probability to reduce assay bias is indicated at 0.428, minimizing both cell type and assay bias. **D,E.** Sampling distribution across cell types (**D**) and assays (**E**) under the optimized probability of 0.428.

### Supplementary References


1. Schreiber, J., Durham, T., Bilmes, J. & Noble, W. S. Avocado: a multi-scale deep tensor factorization method learns a latent representation of the human epigenome. *Genome Biol.* **21**, 81 (2020).
2. Hermann, L., Fiedler, T., Nguyen, H. A., Nowicka, M. & Bartoszewicz, J. M. Beware of data leakage from protein LLM pretraining. *bioRxiv* **261**, 106–116 (2024).
3. Avsec, Ž. *et al.* Effective gene expression prediction from sequence by integrating long-range interactions. *Nat. Methods* **18**, 1196–1203 (2021).
4. ENCODE Project Consortium *et al.* Expanded encyclopaedias of DNA elements in the human and mouse genomes. *Nature* **583**, 699–710 (2020).
5. Dalla-Torre, H. *et al.* Nucleotide Transformer: building and evaluating robust foundation models for human genomics. *Nat. Methods* **22**, 287–297 (2025).
6. Mauvais, F.-X. & van Endert, P. M. Type 1 Diabetes: A Guide to Autoimmune Mechanisms for Clinicians. *Diabetes Obes Metab* **27 Suppl 6**, 40–56 (2025).
7. Schreiber, J. tangermeme: A toolkit for understanding cis-regulatory logic using deep learning models. *bioRxiv* (2025) doi:10.1101/2025.08.08.669296.
